# Origin of exponential growth in nonlinear reaction networks

**DOI:** 10.1101/2020.08.18.254524

**Authors:** Wei-Hsiang Lin, Edo Kussell, Lai-Sang Young, Christine Jacobs-Wagner

## Abstract

Exponentially growing systems are prevalent in nature, spanning all scales from biochemical reaction networks in single cells to food webs of ecosystems. How exponential growth emerges in nonlinear systems is mathematically unclear. Here, we describe a general theoretical framework that reveals underlying principles of long-term growth: scalability of flux functions and ergodicity of the rescaled systems. Our theory shows that nonlinear fluxes can generate not only balanced growth, but also oscillatory or chaotic growth modalities, explaining non-equilibrium dynamics observed in cell cycles and ecosystems. Our mathematical framework is broadly useful in predicting long-term growth rates from natural and synthetic networks, analyzing the effects of system noise and perturbations, validating empirical and phenomenological laws on growth rate, and studying autocatalysis and network evolution.

**Significance:** Natural systems (e.g., cells, ecosystems) generally consist of reaction networks (e.g., metabolic networks, food webs) with nonlinear flux functions (e.g., Michaelis-Menten kinetics, density-dependent selection). Despite their complex nonlinearities, these systems often exhibit simple exponential growth in the long term. How exponential growth emerges from nonlinear networks remains elusive. Our work demonstrates mathematically how two principles, multivariate scalability of flux functions and ergodicity of the rescaled system, guarantee a well-defined rate of growth. By connecting ergodic theory, a powerful branch of mathematics, to the study of growth in biology, our theoretical framework can recapitulate various growth modalities (from balanced growth to periodic, quasi-periodic or even chaotic behaviors), greatly expanding the types of growing systems that can be studied.

## Introduction

Reaction networks are fundamental structures of living systems. They describe biochemical networks (consisting of metabolites, chemical reactions, and macromolecules), ecological networks (based on species and foraging activities), and economic systems (composed of materials and production processes) (1–5). Such networks can exhibit a capacity for growth under suitable conditions. A growing system requires uptake of external components, conversion reactions between internal components, and energy-generating pathways. These processes often involve nonlinear dependencies (3). For example, in cells, many biochemical reactions (e.g., Michaelis-Menten kinetics, feedback inhibition, promoter activation) translate into nonlinear differential equations. Yet, the entire system (total biomass) often converges to exponential growth in the long term, which is a typical property of linear differential equations. Similarly, multi-species communities can often undergo exponential expansion (6, 7). Yet, many interactions in ecosystems, including density-dependent competition and mutation-selection processes, follow nonlinear equations (4). The emergence of exponential growth from a multivariable nonlinear network is not mathematically intuitive. This indicates that the network structure and the flux functions of the modeled system must be subjected to constraints to result in long-term exponential dynamics.

Current growth models, including those based on flux balance analysis (8), proteome partition analysis (9) and general reaction networks (10–12), have been successful at predicting biomass growth. A common limitation of these models is that they assume *a priori* that the system has long-term exponential growth. In addition, they assume that the system exhibits *balanced growth* (Fig. 1A). That is, all components of the system under study are assumed to exponentially increase in amount at the same constant rate, such that their ratios remain fixed during growth of the system. However, in many biological systems, the components are not in balance and display oscillatory or even non-periodic behaviors caused by nonlinear fluxes in the systems. For example, the levels of metabolites and proteins can oscillate during cell growth (13–15). The abundance of species in ecosystems can also oscillate (6) or exhibit non-periodic fluctuations (7). Yet, the biomass of these systems increases exponentially (6, 7, 16). This raises a fundamental question: are there general principles that underlie long-term exponential growth in complex reaction networks?

**Figure 1:**
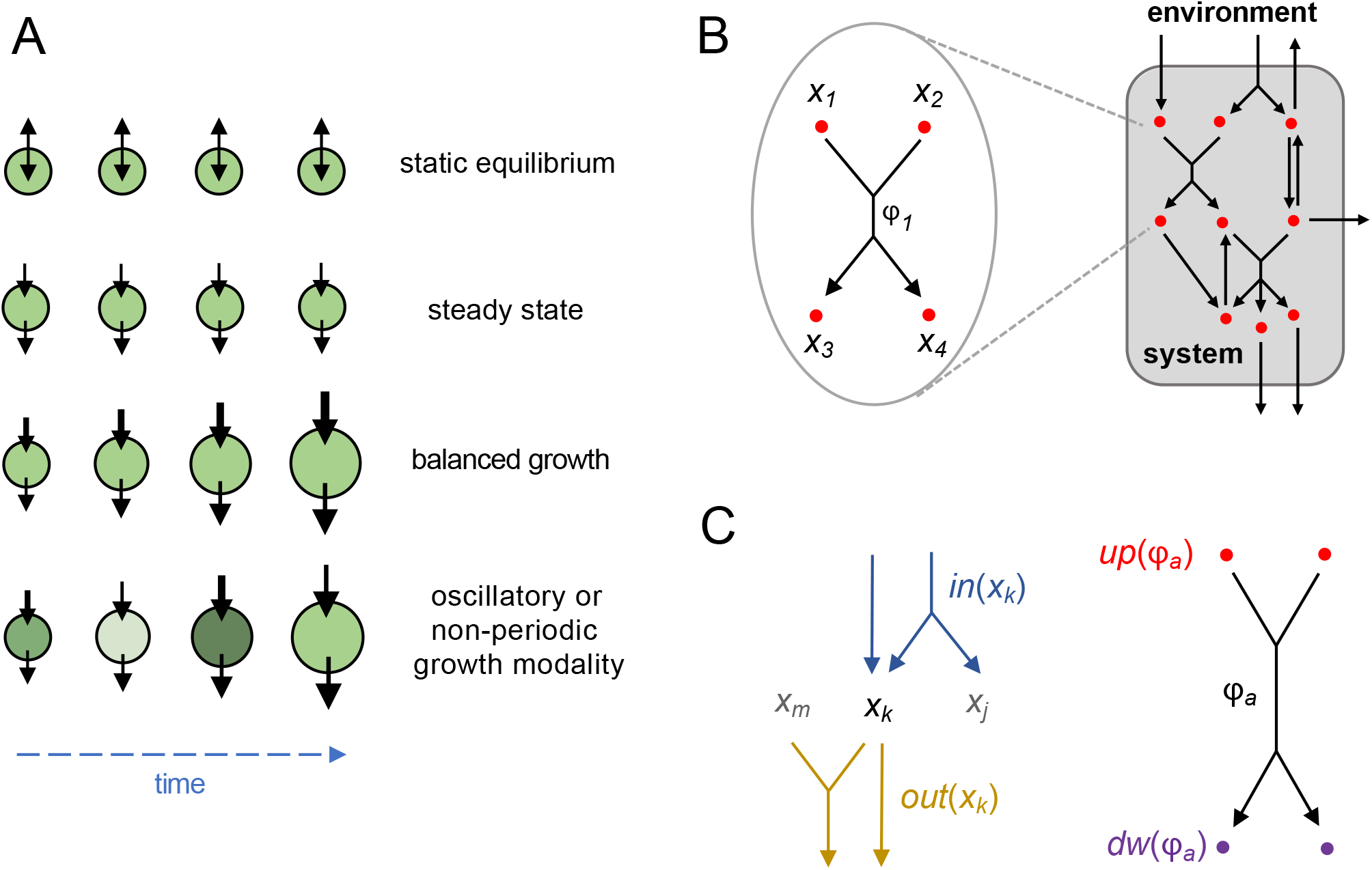
Mathematical formulation of a scalable reaction network. (**A**) Growth modalities for various types of open systems. System size is represented by different sizes of circles; a darker green color represents higher metabolite concentration. Flux magnitude is indicated by the thickness of the black arrows. Static equilibrium indicates that the system is at equilibrium with the environment, with the net flux magnitude equal to zero. Steady state means that influx and efflux are of the same magnitude, and the system is maintained at steady-state with no growth. Balanced growth indicates that the influx is greater than the efflux, and the system is growing proportionally to the fluxes. For oscillatory and non-periodic growth modalities, the influx is greater than the efflux and the system is growing, but the relative proportions of influx and efflux, as well as the metabolite concentration, can vary over time. (**B**) A reaction network is composed of a collection of nodes and reactions. Each reaction describes an interconversion process between upstream and downstream nodes and between these nodes and the environment. (**C**) Terminology describing relations between node *x* and reaction *ϕ*. Given a node *x*_*k*_, the collections of all reactions having *x*_*k*_, as the upstream and downstream nodes are denoted by *in*(*x*_*k*_) and *out*(*x*_*k*_), respectively. Given a reaction *ϕ*_*a*_, the collections of all upstream and downstream nodes are denoted by *up*(*ϕ*_*a*_) and *dw*(*ϕ*_*a*_).

Addressing these questions has been a major challenge due to the lack of a generalizable theoretical framework that goes beyond balanced growth. Ergodic theory is a powerful mathematical tool that has been successfully applied in the fields of fluid dynamics and statistical mechanics for studying the long-term behavior of physical systems (17). The ergodic condition is met when the time average of a quantity is equal to a statistical average over all possible outcomes weighed by their likelihood, also known as a space average. For example, the distance between two children running around a playground independently is a stochastic, time-dependent function; yet, the average distance between them over the long term does not depend on time and can be calculated from each child’s probability distribution over space. In reaction networks, the relevant ‘space’ consists of the set of all possible configurations or states of the entire system, and the overall dynamics constitute a time-evolution process in this state space. A basic requirement for the application of ergodic theory is stationarity of the process under consideration, which means that the likelihood of events does not change with time. Biological systems whose biomass increases over time violate stationarity, and ergodic theory, therefore, cannot be applied directly to study their long-term growth properties.

Here, we demonstrate that ergodic theory can be applicable in growing systems whose dynamics may be properly rescaled such that the growth process is decoupled from the rest of the dynamics. We show that many nonlinear biological reaction networks ―which we define and refer to as *scalable reaction networks* (SRNs)*―* satisfy this condition. By applying ergodic theory on rescaled systems, we show mathematically that scalability and ergodicity ensure that a large class of reaction networks have well-defined long-term growth rates (*λ*), which can be positive (exponential growth), negative (exponential decay), or zero (sub-exponential dynamics). This theoretical framework opens the door to the study and characterization of processes that exhibit different growth modalities, including not only static equilibrium, steady state, and balanced growth, but also oscillatory and non-periodic growth (Fig. 1A). The theory is applicable both for deterministic and stochastic dynamics. It enables one to construct scalable networks of arbitrarily high complexity, predict the growth rate and other dynamical features of the system, and identify autocatalytic networks associated with positive exponential growth.

## Results

### Reaction Network Modeling and Long-Term Growth Rates

We consider a system represented by a collection of nodes within an environment, *E*, that serves as an ideal reservoir for unlimited supply and removal of materials. A *reaction network* is defined by a set of nodes {*x*_1_, …, *x*_*n*_}, reactions {*ϕ*_1_, …, *ϕ*_*m*_}, and flux functions {*J*_1_, …, *J*_*m*_}, which specify the flux magnitude of each reaction. Each reaction represents an interconversion process that consumes materials from upstream nodes and produces materials in downstream nodes (Fig. 1B). The stoichiometric coefficients of interconversion reactions are given by an *n* × *m* stoichiometry matrix *S* (8) (see Supplementary Information (SI)), with positive or negative matrix element *S*_*ka*_ indicating that the reaction *ϕ*_*a*_ acts as an *influx* or *efflux* of node *x*_*k*_, respectively (Fig. 1C). In real-world applications, such networks can describe cellular components (e.g., metabolites and metabolic reactions), cellular populations (e.g., cellular states and transitions), or ecological structures (e.g., species and competitive interactions).

To model the system dynamics, we define a biomass vector 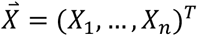 as the total amount of material in each node (in units of mass, e.g., grams), and denote the *system size* by *N* ≡ *X*_1_ + ⋯ + *X*_*n*_. The reaction flux 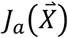 is a multivariate function that specifies the rate of reaction *ϕ*_*a*_, which can depend on any number of components of 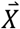, and can be highly nonlinear depending on, for example, the order of the reaction, whether or not it is enzyme-catalyzed, etc. For each node, summing all incoming and outgoing fluxes weighted by the stoichiometric coefficients yields a system of ordinary differential equations that governs the network’s dynamics,

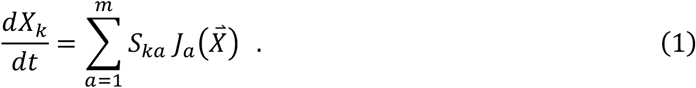

Such systems of nonlinear equations generally cannot be solved analytically and are typically analyzed using numerical algorithms. Currently, the only way to ensure or verify that a reaction network exhibits long-term exponential growth is to run large-scale numerical computations, combined with massive scans of parameter space, and check the results on a case-by-case basis. We therefore sought to identify fundamental features of nonlinear networks that give rise to exponential growth.

We consider the instantaneous growth rate of the system, *μ*(*t*) ≡ (*d*/*dt*) log *N*(*t*), a function that specifies the relative rate of change of the population size at time *t*. If the system grows exponentially at long times, then the long-term time average of *μ*(*t*) yields the system’s exponential growth rate,

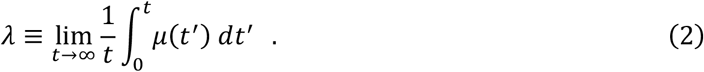

Note that the long-term dynamics of Eq. 2 could result in a system size that grows exponentially (*λ* > 0), decays exponentially(*λ* < 0), or displays sub-exponential dynamics (*λ* = 0). Since most of our discussion is related to growing systems, we will refer to *λ* as long-term growth rate but the reader should keep in mind that the same formula is equally applicable to systems with *λ* ≤ 0.

Mathematically, given an arbitrary set of nonlinear reactions there is no guarantee that the time average in Eq. 2 will converge to a well-defined limit. For example, if the instantaneous growth rate increases linearly with time, its long-term average will diverge, or if *N*(*t*) decays to zero in a finite amount of time, *λ* has no well-defined meaning. More complex behaviors can also emerge in reaction networks, such as periodic, quasi-periodic, and chaotic dynamics, and in such cases the relation of the network dynamics to *λ* is not clear.

We wish to establish general conditions such that the time average given in Eq. 2 converges, implying that the system will grow exponentially. These conditions should (a) ensure that the dynamical system of Eq. 1 has a well-defined solution 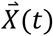 for all initial conditions, (b) imply that the system’s composition can exhibit stationarity even as population size may grow without bound, and (c) enable the time average in Eq. 2 to be computed as a space average over the system’s stationary distribution. We examine the system’s biomass composition vector, 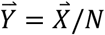, whose components specify the fraction of the total biomass present in node *x*_*i*_. By substituting *X*_*k*_ = *NY*_*k*_ into Eq. 1, we find

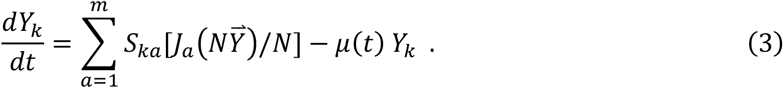

Here, we see that if all fluxes scale up proportionally with *N*, i.e., if the flux functions are *extensive* in the system size, then the term within the brackets would be simply 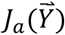, and independent of *N*. Summing both sides over *k* yields the expression for the growth rate,

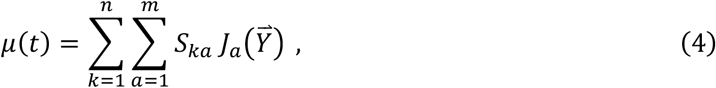

which would likewise be independent of *N*. Then, the system’s composition 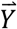 evolves independently of *N* according to the dynamical system in Eq. 3, and the composition alone determines the instantaneous growth rate according to Eq. 4. Hence, we can write 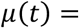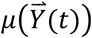(see Fig. S1). Below, we formally specify the conditions that ensure that all fluxes are extensive and that Eq. 3 has a well-defined, physical solution. We will refer to such systems as scalable reaction networks (SRNs).

### Building SRNs: Conditions and Examples

We state the formal conditions that define an SRN and discuss some practical implications and examples. We denote by *Q*^*n*+^ ≡ {*X*_*j*_ > 0} the positive quadrant and by *Q*^*n*^ ≡ {*X*_*j*_ ≥ 0} the non-negative quadrant. A flux function 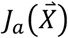 is *scalable* if it satisfies three conditions:

i. 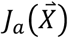is positive on *Q*^*n*+^ and continuously differentiable on *Q*^*n*^\{0} ;
ii. If node *x*_*k*_ is an upstream node of *J*_*a*_(*X*), then *J*_*a*_(*X*) = 0 whenever *X*_*k*_ = 0 ; and
iii. 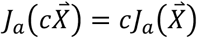 for any *c* > 0.

A reaction network is an SRN if all of its flux functions are scalable.

For an SRN, condition (i) guarantees that the system given in Eq. 1 will have a unique solution. Condition (ii) requires the flux functions to be *upstream-limited*, that is, whenever an upstream node is depleted, its connected efflux must be zero. This ensures that solutions are physical, i.e. the trajectories remain within *Q*^*n*^. Condition (iii) requires the flux magnitude to be extensive in the system size, which enables projection onto the simplex and the application of ergodic theory as detailed above.

We would like to emphasize that a multivariate flux function satisfying condition (iii) does not necessarily need to be linear. There is a wide range of commonly occurring flux functions that are not linear and yet scalable (Table 1), which one can verify by checking conditions (i) to (iii) directly.

More generally, biochemical fluxes are typically written as nonlinear functions of metabolite concentrations 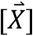 (mass per volume). If, in addition, the system volume (*V*) scales with the total biomass *N*, i.e., *V* = *bN* with constant *b*, the fluxes become scalable. To see this, we denote by 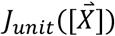 the metabolic flux magnitude per unit volume. Metabolic fluxes usually obey a mass-action law, e.g. *J*_*unit*_ ∝ [*X*_1_]^*A*^[*X*_2_]^*B*^ or quasi-steady-state kinetics, e.g. 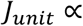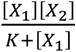. The total flux magnitude 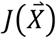 scales with volume and can be expressed as

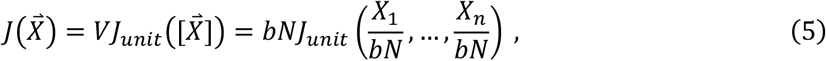

which satisfies the scaling condition (iii). Therefore, under the volume-biomass scaling assumption, biochemical flux functions belong to the class of scalable flux functions. When volume-biomass scaling is not exact, a larger class of networks, which we refer to as *asymptotically scalable*, is often applicable with similar results (see SI and Table 1).

### Ergodicity and Dynamics in SRNs

We are now positioned to apply ergodic theory to evaluate the time average of Eq. 2. The scalable condition of SRNs allows us to rescale the system and focus on the long-term dynamics of the composition vector 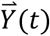 in the unit simplex Δ^*n*−1^, the space of all vectors (*Y*_1_, …, *Y*_*n*_)^*T*^ satisfying 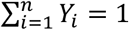 with *Y*_*i*_ ≥ 0 for all components *i*. The distribution on the simplex space, or likelihood of 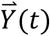 being located in a given region of Δ^*n*−1^, can be regarded as a probability measure *ω* defined on Δ^*n*−1^, satisfying the normalization condition 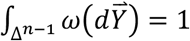. We are interested in probability measures *ω* that are invariant under the time evolution of Eq. 3, i.e., probability distributions with respect to which 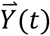 is stationary. Invariant measures are sometimes decomposable, meaning they are weighted sums of invariant measures supported on disjoint invariant sets. Those that cannot be decomposed are called *ergodic measures*. Ergodic measures are guaranteed to exist for continuous flows on a compact metric space such as Δ^*n*−1^, and they have the important property that time averages of observables can be equated with space averages, a property we now use to compute exponential growth rates.

Given an ergodic measure *ω*, we can apply the Birkhoff ergodic theorem to evaluate Eq. 2, yielding

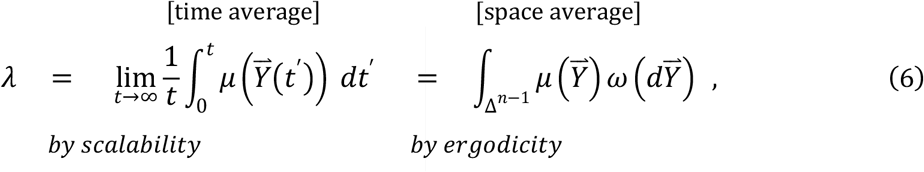

where the second equality holds for almost every initial condition 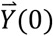, except possibly on a set of measure zero with respect to *ω*. Since Δ^*n*−1^ is a compact space, 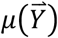 is bounded hence the space average in Eq. 6 is finite. Thus, the long-term growth rate *λ* defined in Eq. 2 converges and its value is independent of initial conditions (except for a set of *ω*-measure zero, see SI, section 2 for an example and a detailed discussion). It is also straightforward to show that if a trajectory exhibits exponential growth, any other trajectory that converges to the first one will have the same long-term growth rate. Thus, depending on the structure of *ω*, a wide range of trajectories exhibit exponential growth with the same rate.

We illustrate the above point with a few concrete scenarios in low dimensional systems, though it is important to note that the real power of the results is that they hold for high dimensional SRNs, i.e., for networks with an arbitrary number of nodes. Here, we consider 3-node SRNs, where the composition vector lies in the two-dimensional simplex Δ^2^.

Let’s first discuss deterministic dynamics without noise. Flows exhibiting a limit cycle, bistability, or a heteroclinic cycle are shown in Fig. 2A-C (the support of the ergodic measures is shown in green, and any regions lying outside of the support have measure zero). In Fig. 2A, the ergodic measure is supported by a limit cycle, and thus *λ* converges along the limit cycle. All other trajectories converge to the limit cycle trajectory, which implies that the system eventually grows exponentially with the same rate *λ* regardless of the initial condition. In Fig. 2B, we show a bistable system in which two ergodic measures are shown, each supported on a different fixed point. A trajectory that remains at one fixed point will grow exponentially, but the rate of growth can be different for each fixed point; all trajectories that converge to a given fixed point will grow exponentially with the same rate. In Fig. 2C, ergodic measures exist at each of the simplex vertices, and a heteroclinic cycle lies on the boundary. Trajectories starting in the interior of the simplex spend progressively longer amounts of time near each of the vertices without converging to any one of them. In this case, although the long-term growth rate would generally not converge, exponential growth will be achieved for increasingly long periods of time while the trajectory remains in the vicinity of each fixed point.

**Figure 2:**
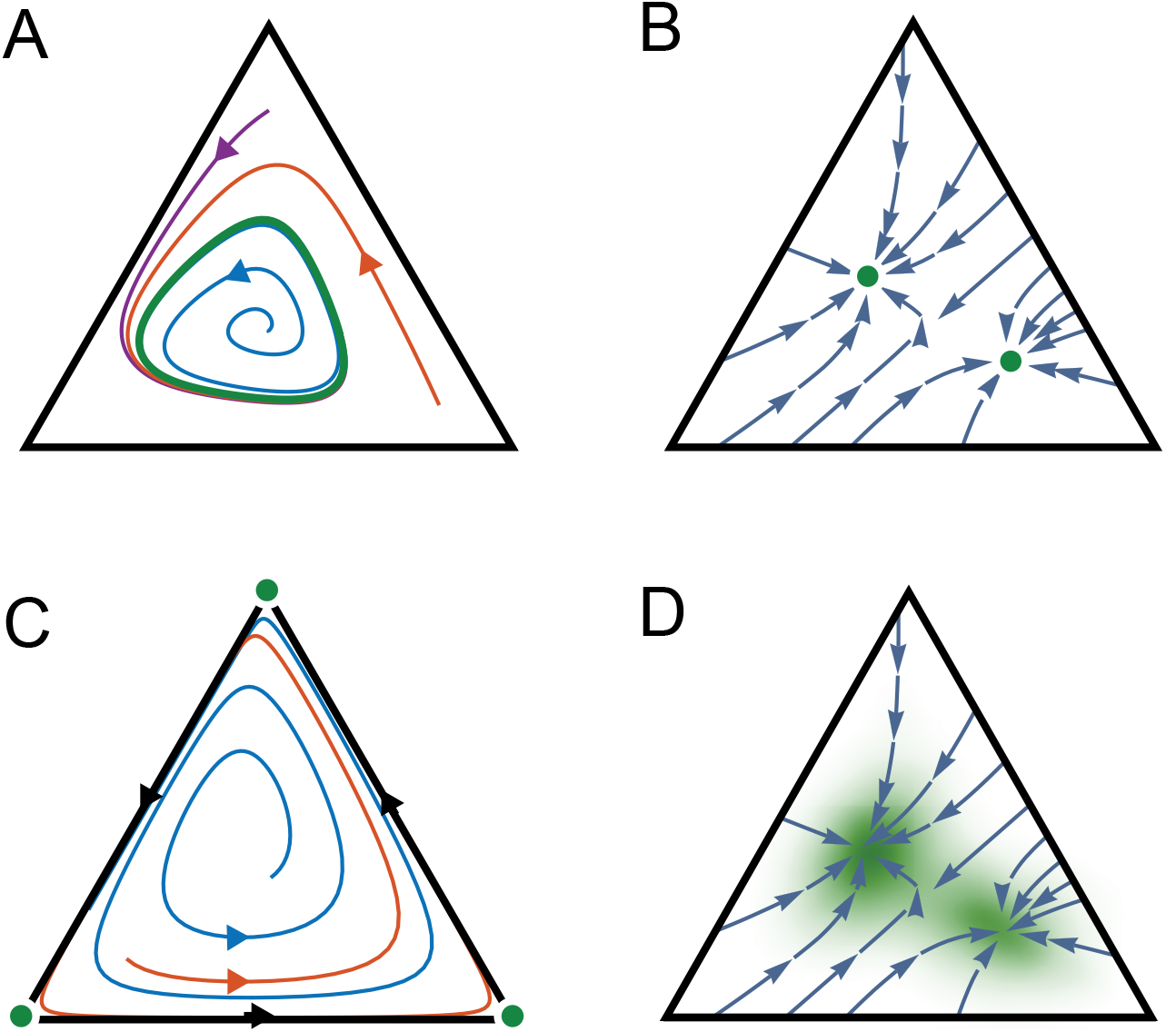
Dynamics of three-node SRNs and ergodic measures. Each panel shows representative trajectories of the composition vector 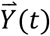 on the simplex Δ^2^, where vertices correspond to network composition states (1,0,0), (0,1,0), or (0,0,1), and any point in the simplex corresponds to a linear combination of these. Ergodic measures are shown in green. Dynamics with a stable limit cycle; the ergodic measure is supported on the limit cycle. (**B**) A bistable system with two stable fixed points; two ergodic measures are shown, supported on each of the stable fixed points. (**C**) Dynamics with a heteroclinic cycle on the boundary; ergodic measures exist supported at each of the vertices which are unstable fixed points. Trajectories approach each vertex for a progressively longer time before moving close to the next vertex, and so forth. (**D**) Stochastic dynamics were simulated for the bistable system of and the ergodic measure is shown as a green density within the simplex; the density peaks near the fixed points of the deterministic flows shown in blue.

We see that while ergodic measures determine the long-term growth rates of SRNs, complex scenarios involving multiple ergodic measures are possible even in low dimensional systems, and the situation only gets more complicated in higher dimensions. However, in the presence of random noise, one generally obtains dynamics that are better behaved, and are described by a single ergodic measure. All of the ergodic theory results described above carry through for random dynamical systems, specifically when Eq. 3 is formulated as a stochastic differential equation to model noise in the system’s composition. Fig. 2D shows the bistable network of panel 2B with the addition of noise. A single ergodic measure exists on the interior of the simplex, concentrated near the two fixed points but also exhibiting non-zero measure between them, indicating that stochastic trajectories spend time sampling both fixed points. Thus, an SRN whose deterministic dynamics exhibit distinct growth rates dependent on the initial conditions can, due to the presence of noise, grow exponentially with a single rate for all trajectories. The timescale for a stochastic trajectory to converge to an ergodic measure depends on the magnitude of diffusion and drift on the simplex (18). Mathematical techniques, such as large deviations theory and mean first passage time calculations (e.g., 19, 20), can be applied for estimating the timescale for convergence.

In summary, ergodic measures underlie exponential growth modalities in SRNs. They can be used to analyze these modalities for both deterministic and stochastic dynamics. We found two general criteria for the long-term growth rate of an SRN to converge (see SI, Supplementary text for proof):

- For deterministic dynamics, if a trajectory 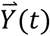 converges to a fixed point, a periodic or a quasiperiodic orbit, then it exhibits a well-defined long-term growth rate *λ*, which is determined by the ergodic measure for the given attractor according to Eq. 6.
- For stochastic dynamics, if one adds random diffusion to the composition vector, and if SRNs can always regenerate any component that goes to zero from other components (we call these *regenerative* SRNs), then all trajectories have the same exponential growth rate regardless of the initial conditions.

Our results extend also to chaotic dynamics, as we show below. Further details and generalizations are provided in SI. In the following sections, we use concrete examples to illustrate the versatility of the SRN framework. This framework allows us to implement nonlinear flux functions in reaction networks to study growth dynamics beyond balanced growth and perform systematic parameter scans for long-term growth rate analysis.

### Growth Modalities of SRNs

SRNs that display exponential growth can exhibit various growth modalities. The simplest is *balanced growth*, in which all components of the system grow at the same unchanging rate (Fig. 1A). This modality corresponds to exponential growth along a constant vector 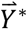, and according to Eq. 6, it is a special case in which an ergodic measure is concentrated entirely at 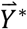, i.e. 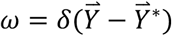 and 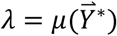. In general, *ω* can have forms that correspond to various growth modalities. They include periodic growth as in metabolic cycles (14, 21), cell cycles (22–24), oscillating populations (6, 25), and economic business cycles (26) as well as different types of aperiodic growth such as chaos in microbial food webs (7). Such systems cannot be modeled according to a balanced growth assumption, yet their ability to exhibit exponential growth over long timescales follows the general principles of ergodicity and scalability described above.

To demonstrate the utility of SRNs for the study of non-balanced growth, we consider two explicit examples with various growth modalities for which long-term growth rates cannot be calculated by using a balanced-growth approach. We first constructed an SRN consisting of a repressilator-type regulatory circuitry. The repressilator is an autonomous nonlinear genetic circuit that is well-known for generating oscillations of protein expression in bacterial cells (27). To create a growing system, we incorporated the repressilator circuit into an autocatalytic network (Fig. 3A). In this model, the synthesis fluxes *J*_2_, *J*_3_ and *J*_4_ are subject to the repression of nodes *x*_3_, *x*_4_ and *x*_2_, respectively. These sigmoidal flux functions depend on the Hill coefficient (*θ*) and the repression strength (*K*). By varying *θ*, we found that this network is able to change growth modes. For small *θ* (e.g., *θ* = 1), the system exhibits balanced growth (Fig. 3B), while larger *θ* (e.g., *θ* = 4) causes the system to display a periodic growth modality (Fig. 3C). Here, the bifurcation happens around *θ* ≈ 2 upon a change in the fixed-point stability of Eq. 3. Interestingly, the oscillation period and growth rate are strongly coupled in this model. For example, under high repression strength, growth rate *λ* sharply increases around *θ* ≈ 2 (Fig. 3D), where the attractor of 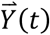 transitions from a fixed point to a limit cycle (Fig. 3E). The transition from fixed point toward limit cycle is accompanied with fast increase of growth rate. This intriguing behavior is due to a change in the network composition at the transition, such that the relative biomass fraction of nodes *x*_2_, *x*_3_, *x*_4_ sharply increases promoting greater autocatalysis influx (*J*_1_ in Fig. 3A), which yields higher values of *λ*.

**Figure 3:**
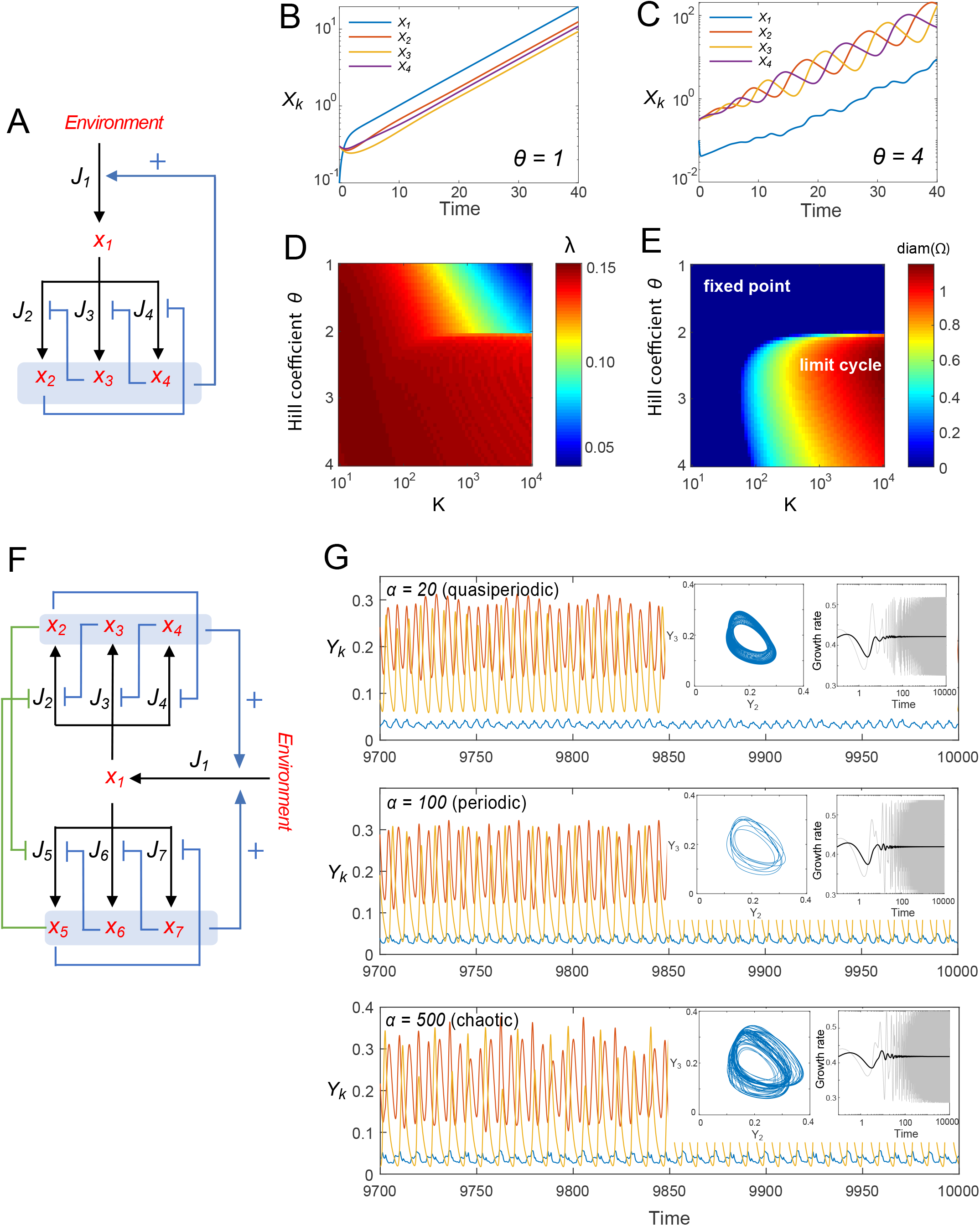
Scalable reaction networks with various growth modes. (**A**) An autocatalytic, single-repressilator SRN. Blue lines ending with an arrow represent flux activation, whereas blue lines ending with bars indicate flux repression. The flux functions *J*_2_, *J*_3_, *J*_4_ have sigmoidal forms, e.g., 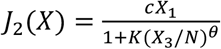. (**B**) Balanced growth (*θ* = 1, *K* = 500) for the network shown in (A). (**C**) Periodic growth modality (*θ* = 3, *K* = 500) for the network shown in (A). (**C**) Phase diagram of *λ* in the parameter space (*θ*, *K*) for the network shown in (A). (**e**) Phase diagram of the diameter (diam) of the attractor (Ω) for the network shown in (A). In this regime, Ω is either a fixed point or a limit cycle in Δ^3^. The diameter of Ω is defined by 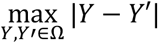; it is zero for fixed points and positive for limit cycles. (**F**) An autocatalytic double repressilator SRN. The parameter *α* is the ratio of repression strengths between the two oscillators (see SI for details). (**G**) Growth modalities of the double repressilator. Dynamics can be quasiperiodic (*α* = 20), periodic (*α* = 100), or chaotic (*α* = 500). Left insets: projection of attractors on *Y*_2_ − *Y*_3_ coordinates. Right insets: instantaneous growth rate *μ*(*t*) (gray) and long-term average 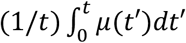 (black).

By combining two mutually-inhibiting repressilator networks (Fig. 3F), we obtained an SRN capable of exhibiting additional growth modalities, ranging from quasiperiodic to non-periodic, chaotic growth (Fig. 3G, see Fig. S3 for the full bifurcation diagram and the largest Lyapunov exponent signature). Under a chaotic growth modality, the trajectory of 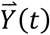 is intrinsically unpredictable in the long term. However, in the presence of noise, this regenerative SRN is guaranteed to have a single ergodic measure with density in the interior of the simplex, and therefore a well-defined long-term growth rate, which we calculate using Eq. 6 in the small-noise limit (Fig. S3C). Numerical simulation indicates convergence of *λ* (Fig. 3G, insets). This example demonstrates that the long-term growth rate of SRNs can be robust, even when the components of the network fluctuate indefinitely.

### Exponential growth of ecosystems with interspecies competition and cross-feeding

In ecological and population dynamic contexts, SRNs provide a basis for understanding how exponential growth can emerge from nonlinear interactions between different species, such as in natural expansion of ecosystems. In addition, the SRN model is also applicable for turbidostat and chemostat experiments. In a turbidostat, nutrients typically remain in excess at all times and cell density is maintained constant by a feedback loop that adjusts the dilution rate. In such a bioreactor, the long-term average dilution rate is equal to the long-term growth rate of the population and therefore can be predicted by the SRN model. In a chemostat, an essential nutrient is in limiting concentration while the dilution rate is held a constant value set by the user. We provide an SRN model that enables the long-term dynamics of chemostats to be analyzed under the SRN framework (see SI, section 2.8).

To illustrate these points, we used an SRN framework to construct a three-species community in which the species compete through a classical Lotka-Volterra-type interaction (28) (Fig. 4A). In addition, in this model, each species (*x*_1_ to *x*_3_) secretes a metabolite (*m*_1_ to *m*_3_) that can cross-feed the other two species. The efficiency of cross-feeding (CF) and the competitive interactions are allowed to be asymmetric. To mimic the diversity of natural interactions among species, we took advantage of the SRN framework to scan 150,000 different parameter sets of species interactions and studied long-term species coexistence (see SI, Methods).

**Figure 4:**
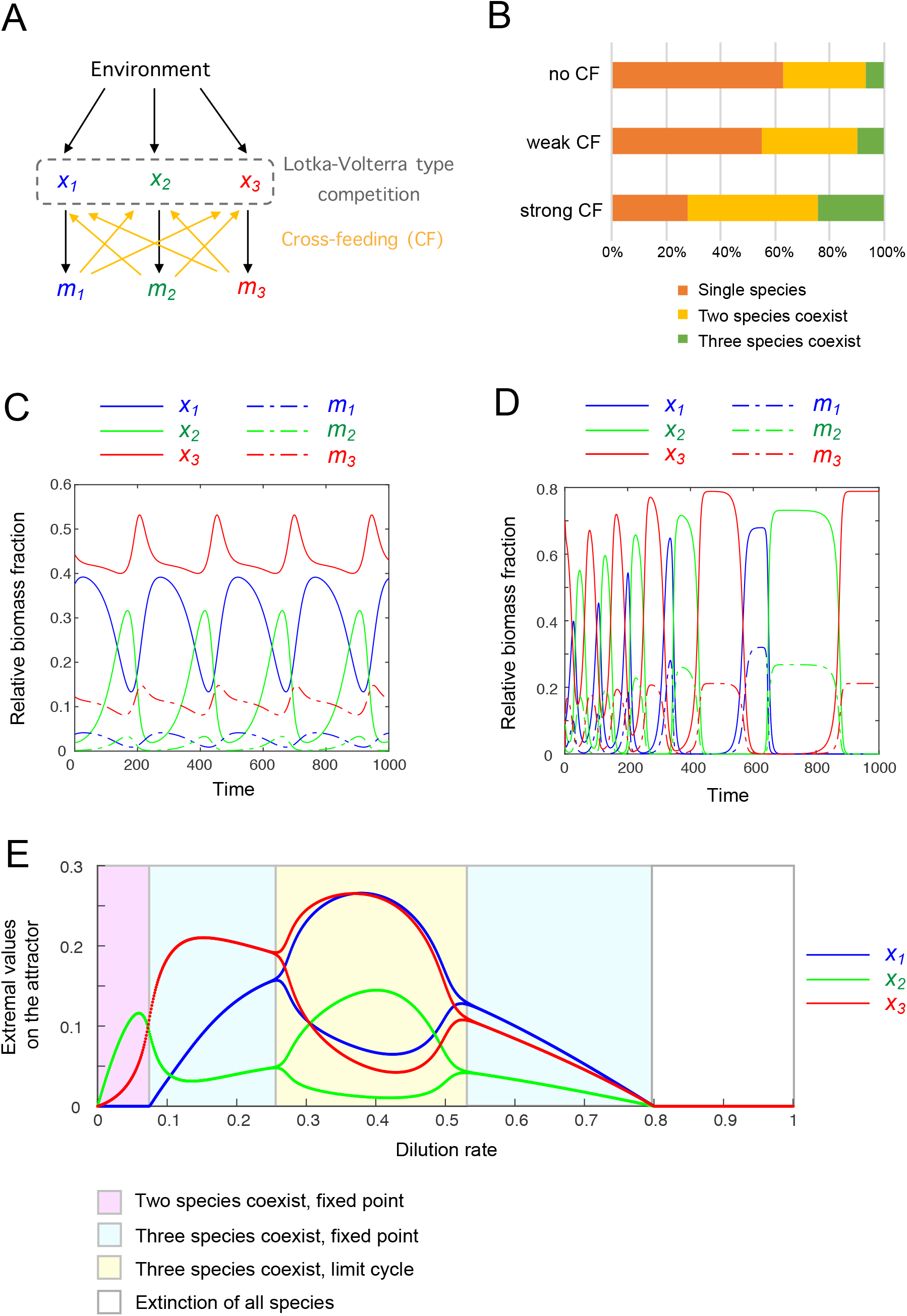
Models of three species that mutually compete and cross-feed under the condition of unlimited (turbidostat-like) or limited (chemostat-like) external resources. See SI (Methods) for model details. (**A**) Cross-feeding reaction network with three species (*x*_1_ to *x*_3_) and three secreted metabolites (*m*_1_ to *m*_3_). The Lotka-Volterra-type competition is described by quadratic functions, and the cross-feeding fluxes follow a Michaelis-Menten function. (**B**) Species coexistence statistics when cross-feeding is absent, weak (cross-feeding efficiency between 0 and 0.1) or strong (cross-feeding efficiency between 0 and 0.5) for a turbidostat-type system. The three-species coexistence includes growth modalities of a fixed point, a limit cycle and a heteroclinic cycle. (**C**) An example from (B) where the cross-feeding system has a limit cycle growth modality. (**D**) An example from (B) where the cross-feeding system has a heteroclinic cycle growth modality. (**E**) Long-term growth dynamics of the cross-feeding population (shown in A) for a chemostat-type system as a function of the dilution rate, *D*.

We initially performed our analysis assuming that the environmental nutrient is unlimited (turbidostat-type systems). For a first dataset, we simulated random competitive interactions without cross-feeding. We found that, in the long term, more than 60% of the simulated systems consist of single species, with the remaining systems consisting of two or all three species (Fig. 4B). For a second dataset, we simulated random competitive interactions plus random weak cross-feeding (with efficiency parameters between 0 and 0.1; see SI, Methods). Increasing the efficiency of cross-feeding increased the chance of species to coexist (Fig. 4B). For a third data set, we allowed stronger cross-feeding to occur (with efficiency parameter between 0 and 0.5), which further increased the probability of species coexistence (Fig. 4B). We found that different growth modalities can emerge when three species coexist. They include not only examples of balanced growth (fixed point), but also examples of limit cycles (periodic growth, Fig. 4C) and heteroclinic cycles (growth with increasing period, Fig. 4D). As shown mathematically (see above and SI), a rescaled system with a fixed point or limit cycle has a well-defined growth rate. In the case of an ecosystem with a heteroclinic cycle, one of the three species will dominate alternately with increasing periods. Exponential growth is achieved when the trajectory *Y*(*t*) is at the vicinity of the saddle points of the heteroclinic cycle. For deterministic dynamics the heteroclinic cycle can be arbitrarily close to the simplex boundary. In stochastic dynamics of finite populations, a species would go extinct when the last individual is lost, and the system could end up at a fixed point.

Next, we considered the case when the concentration of a nutrient is limiting (chemostat-type systems) by modifying the cross-feeding model to a chemostat-type equation and including the limited nutrient into the equation (see SI, section 2.8). By scanning 120,000 parameter sets, we found that the cross-feeding model can, in specific cases, exhibit transitions in growth modality as the dilution rate is varied. This is illustrated in Fig. 4E where the long-term dynamics can be either two-species fixed point, three-species fixed point, three-species limit cycle or extinction depending on the dilution rate *D*. The bifurcations of this system can therefore be studied using the SRN framework in the future.

Our results provide a theoretical basis to understand how ecosystems can grow and expand exponentially even when exhibiting complex, unbalanced dynamics among not only species but also resources (e.g., due to cross-feeding, nutrient limitation). We envision that similar SRN-based models can be constructed for economic systems to study their growth (e.g., business cycles (26)).

### Exponential growth of biosynthesis with energy allocation

Often reaction networks are modeled to study cellular metabolism and biomass growth, which can be done using SNRs. As a proof of concept, we constructed a simplified biosynthesis model with 24 nodes using common metabolic flux functions such as Michaelis-Menten equations (Fig. 5A, also see SI for details). The advantage of using an SRN framework is that it enables unconstrained parameter scans of the system while automatically ensuring that various unphysical behaviors are avoided (e.g. the system size blowing up at a finite time, or biomass components becoming negative). There is no need to impose constraints based on empirical objective functions or phenomenological laws. Instead, as we illustrate below, the SRNs model can be used to uncover a phenomenological law.

**Figure 5:**
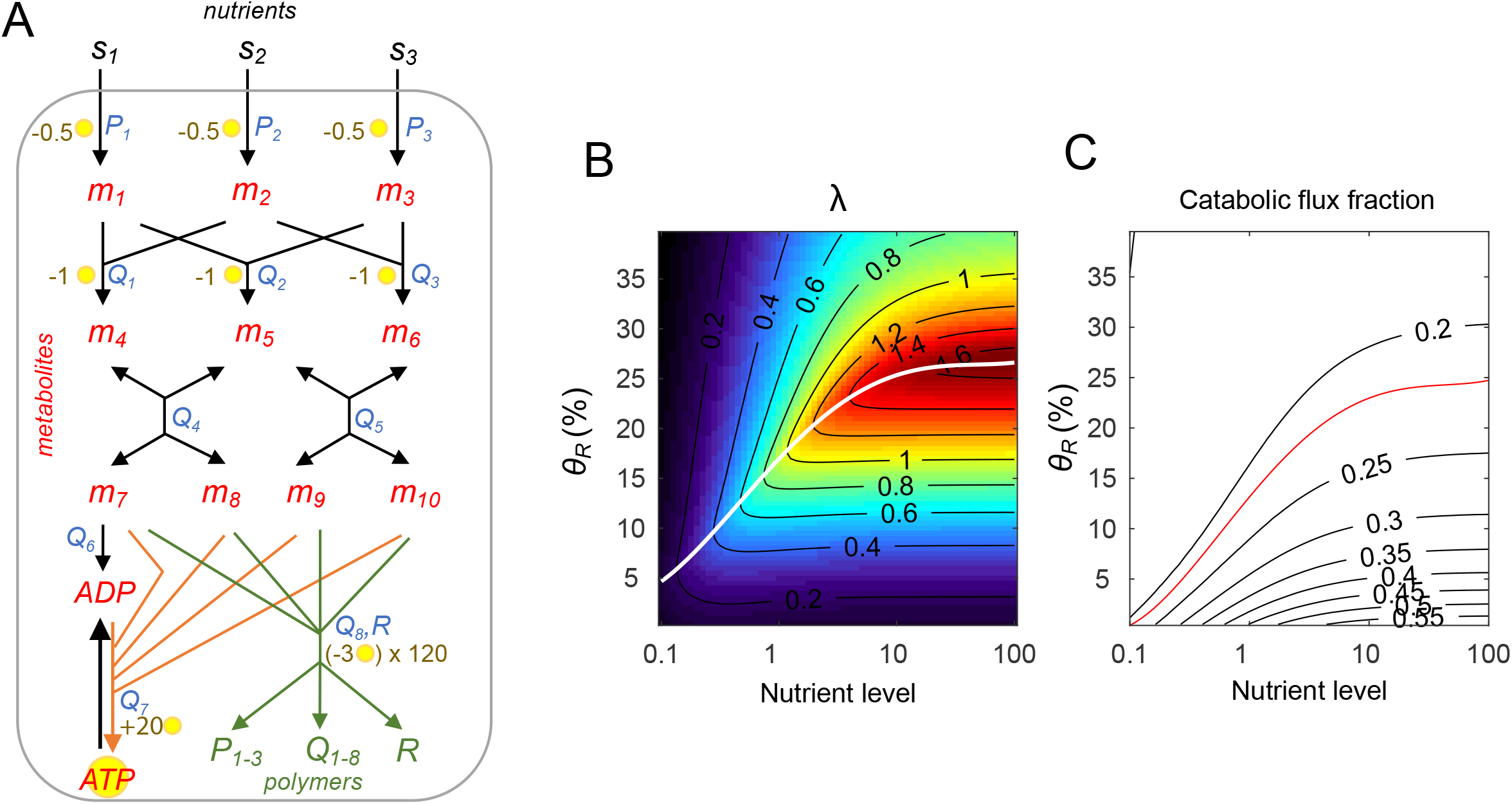
Toy model of an autocatalytic biosynthesis network. (**A**) Diagram of the biosynthesis network. Nodes labeled in red and green correspond to metabolites and polymers, respectively. The external resources *s*_1_, *s*_2_, *s*_3_ are imported into the system and converted into metabolites *m*_1_ to *m*_6_ through a series of reactions. These metabolites are used to produce amino acids *m*_7_ to *m*_10_, which can either enter anabolic pathways (green arrows) to produce polymers such as transporters (*P*_1-3_), enzymes (*Q*_1-8_), and ribosomes (*R*), or enter catabolic pathways (orange arrows) to replenish cellular energy in the form of ATP from ADP. Text in blue next to a reaction represents the catalytic enzyme of this flux. The yellow circle next to a reaction represents the number of ATP produced or consumed by this reaction. For details about the model and flux functions, see SI. (**B**) Growth rate under different ribosomal synthesis strengths (*θ*_*R*_) and varying external nutrient levels (set to be the same level for *s*_1_ = *s*_2_ = *s*_3_). The white line indicates the optimal growth condition under each external nutrient level. (**C**): Contour line of the catabolic flux fraction of amino acids (*m*_7-10_). The red line indicates the optimal condition predicted by the objective function (see main text).

We built a toy model in which the reaction network imports three types of external nutrients (*s*_1_ to *s*_3_) and converts them to amino acids (*m*_7_ to *m*_10_) through intermediate metabolites (*m*_1_ to *m*_6_) (Fig. 5A). The generated amino acids are then utilized in polymer synthesis (anabolic pathways, green) or energy production (catabolic pathways, orange). Biosynthesis produces transporters (*P*_1-3_), enzymes (*Q*_1-8_), and ribosomes (*R*), while energy production replenishes ATP from ADP. The polymers and energy molecules are, in turn, required for catalyzing the upstream fluxes (see SI, Methods (section 4) for details). Using parameter values that are compatible with physiological conditions (SI, Table S4), the system trajectory *Y*(*t*) converges to a fixed point or a limit cycle. Hence, the long-term growth rate can be calculated using Eq. 6, and the correlation between growth rate, metabolic levels and flux magnitudes can be analyzed for various parameters.

Using this model, we investigated how growth rate is affected by the balance between catabolism and anabolism, which is known to be essential to achieve optimal biosynthesis (29, 30). We set values of ATP consumption and production within a reasonable range (31). Specifically, consumption of one amino acid (*m*_7-10_) produces 20 ATP while the synthesis of an amino acid costs 2 ATP. Utilization of amino acids for polymer synthesis costs 3 ATP. Therefore, for each amino acid, the system either produces 18 ATP through the catabolic pathway (orange arrows) or consumes 5 ATP through the anabolic pathway (green arrows) for polymer synthesis (Fig. 5A). If the flux magnitudes of the catabolic and anabolic pathways have a ratio of 5:18, the production and consumption of ATP will be balanced. Heuristically, 5/(5+18) = 0.22 would be the *optimal catabolic flux fraction*. This value can be used as a target for an objective function on energy balance.

To determine whether this objective function predicts the optimal growth rate, we varied the external nutrient level and the ribosomal synthesis strength in simulations. In Fig. 5B, the condition with the highest *λ* across various nutrient levels is indicated by the white curve (left graph). In Fig 5C, the optimal objective function (catabolic flux fraction = 0.22) is given by the red curve. The similarity between the white and red curves shows that the objective function can indeed predict optimal growth. When we varied the ATP production stoichiometry to 10, 20 or 30 ATP per amino acid, the optimal catabolic flux fractions became 0.38, 0.22 or 0.15 in this model. In all cases, the optimal growth rate determined by simulation strongly correlated with the optimal growth rate predicted by the objective function (see Fig. S4).

Altogether, our results show that our biosynthesis toy model can be used to validate the objective function of ATP balance for optimal growth. By using an SRN formulation, this type of model can be easily generalized to include any number of nodes and reactions.

### Autocatalytic circuits for SRNs

For a system to grow autonomously (e.g., free-living organisms), it must contain a reaction network that is autocatalytic. What makes a network autocatalytic? Theorists have addressed this fundamental question and identified design principles of autocatalysis using discrete symbolic models (32, 33). As shown below, by using SRNs, we can bridge from symbolic models to continuous dynamical systems (described by differential equations), demonstrating that the existence of autocatalytic properties is a necessary condition for a positive long-term growth rate (*λ* > 0).

Here, we refer to a reaction *ϕ*_*a*_ as *maintained by node x*_*z*_, if *J*_*a*_(*X*) = 0 whenever *X*_*z*_ = 0, and designate the *maintenance set* of *ϕ*_*a*_ to be the collection of nodes that maintain *ϕ*_*a*_, which we denote as *mt*(*ϕ*_*a*_). Note that the maintenance set of the reaction *ϕ*_*a*_ only includes the components that directly affect the magnitude of the flux *J*_*a*_. For example, in a biochemical network, the immediately upstream precursors, the enzymes and the co-enzymes that are directly involved in the reaction are all essential for generating a positive flux. Hence, they all belong to the maintenance set of the given reaction.

A collection of reactions *K* is called an *autocatalytic circuit* if

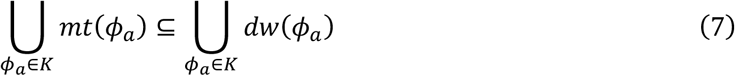

where *dw*(*ϕ*_*a*_) denotes the downstream nodes of reaction *ϕ*_*a*_ (see Fig. 1C). Intuitively, an autocatalytic circuit is a collection of reactions capable of *synthesizing their own maintenance set* (see Fig. S5 for an example). In biological systems, an autocatalytic circuit contains all central metabolic reactions and synthesis pathways for making essential macromolecules. We call a reaction network *autocatalytic* if it contains at least one autocatalytic circuit.

With the SRN formulation, we are able to establish a connection between the sign of *λ* (which is an analytical property given by Eq. 6) and the presence of an autocatalytic circuit (which is an algebraic property given by Eq. 7). Our main result is the following (see SI, Supplementary text (section 6) for proof):

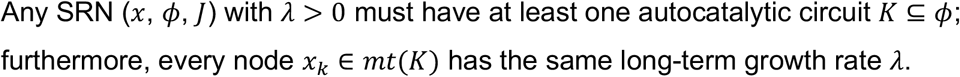

The above result provides the basic topological constraint: if an SRN does not have an autocatalytic circuit, it cannot exhibit positive long-term growth rate under any parameter set and any initial condition. This topological constraint can be used to identify autocatalytic circuits and to rule out non-growing topologies from thousands of random networks (see SI and Figure S6 for examples of random reaction networks with a power-law connection probability).

## Discussion

Long-term growth is an essential property of living systems. How this remarkable property is achieved and maintained is one of the most fundamental questions in biology. In this study, we identify an important class of biological reaction networks, SRNs, whose long-term growth properties can be studied using powerful ergodic theory tools. With these tools, we mathematically demonstrate two basic principles of exponentially growing systems: scalability of the underlying flux functions and ergodicity of the rescaled system. Our mathematical framework explains how various growth dynamics (balanced growth, oscillatory or non-periodic) driven by complex, nonlinear flux functions can converge to long-term exponential growth, which is prevalent in biological systems.

Our theory has a number of practical implications as it provides a rigorous mathematical foundation for modeling and probing natural and synthetic reaction networks with distinct advantages over existing methodologies. First, our approach does not *a priori* assume that a long-term growth rate exists; instead, this is a mathematically derived consequence of scalability and ergodicity. Second, as current methods are limited to the case of balanced growth, our theory considerably expands the type of systems that can be analyzed, including metabolic cycles (13, 14, 21, 23) and chaotic oscillations in communities (7). Third, once an SRN is constructed, there is no requirement for the dynamics of the system, 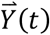, to be constrained by objective functions or phenomenological laws. In fact, with our approach, objective functions or phenomenological laws can be inferred or validated through large-scale simulation across the parameter space (Fig. 5 and Fig. S4). Fourth, biological processes are stochastic in nature and thereby inherently noisy, yet noise is often neglected in current growth models. The ergodic theory formulation enables generalization from ordinary differential equations to stochastic differential equations and allows rigorous consideration of how noise impacts long-term growth.

From a technical standpoint, our study bridges two fields that seldom interact: ergodic theory and growth modeling. The key concept underlying this connection is the scalability of multivariate functions in reaction networks. This allows one to leverage a powerful branch of mathematics in the study of biological growth processes. Interestingly, in the field of ergodic theory, noise has emerged as an important factor to consider, as its averaging effect tends to produce simpler, more tractable dynamics, while at the same time capturing the system’s essential features (34–36). Consistent with this observation, we find that noise can enable SRNs to achieve ergodicity and thus helps systems attain a robust long-term growth rate (see SI, Supplementary Text, and Fig. S3). This provides a novel perspective on the importance of noise in biology.

Living systems (e.g., cells) are highly complex in their reaction network structure. Hence, realistic models of such systems can include hundreds to thousands of components (nodes) and reactions (fluxes) (37). Simulating this level of complexity is computationally challenging, largely because of open problems regarding i) the stability of models, ii) the robustness of results to the choice of parameters and initial conditions, and iii) the behavior of the dynamics over long timescales. In our framework, an SRN regardless of its complexity can be projected into a bounded simplex space while preserving all of the key dynamical information. This results in well-behaved topological and analytical properties (simply connected space, straightforward parametrization, etc.), which will help address the aforementioned open problems.

Our mathematical framework is applicable for both exponential growth and decay. This allows one to study when a system transitions from growth to decay (or vice versa), which can be critical in the context of ecosystem management and economic development (38, 39). For a system to grow and expand exponentially, it must achieve *λ* > 0. Under the SRN framework, we prove that a sufficient condition for achieving *λ* > 0 is the existence of an autocatalytic circuit. Autocatalysis has been extensively studied, primarily using symbolic models (32, 33). Continuous dynamical models have also been reported (e.g., (40–42)). However, they are usually restricted to linear networks while the nonlinear network models are specialized for particular systems because of the absence of a theoretical framework. In contrast, our theory provides a generalizable and systematic way to connect symbolic and continuous models to analyze long-term growth rates from any scalable reaction network. We envision that our mathematical framework will facilitate future studies in subjects as diverse as network design and evolution, biochemical modeling of the cell, ecosystem growth and sustainability, and economic business cycles.

## Methods

See SI for detailed methods for theoretical derivation and numerical simulation.

## Acknowledgments

We thank Drs. Thierry Emonet, Hee Oh, and Alvaro Sanchez for discussions. We also thank Dr. Damon Clark, Markus Covert, Marco Cosentino Lagomarsino and the members of the Jacobs-Wagner laboratory for their input and critical reading of an earlier version of the manuscript. We are also grateful to Dr. Ned Wingreen for constructive suggestions that improved the manuscript. C. J.-W. is an Investigator of the Howard Hughes Medical Institute. This work was partially funded by NIH grant R01-GM120231 to E.K. and NSF Grant 1901009 to L.-S.-Y.

## Authors contributions

W.-H.L. conceived the idea and conducted the theoretical and computational studies. C.J.-W. and E.K. supervised the research. C.J.-W., E.K. and L.-S.Y. provided conceptual guidance. W.-H.L., E.K. and L.-S.Y. contributed to mathematical proofs. All authors wrote and edited the manuscript.

## Authors information

The authors declare no competing financial interest. Simulation code used to generate the results shown in this study can be found at https://github.com/JacobsWagnerLab/.

## Supplementary Information

## Supplementary text

### Mathematical theory of scalable reaction networks

#### 0. Introduction

A flux network is a fundamental structure that describes the interconversion of materials or resource in chemical, biological, and economic models. Although the transient dynamics for a general flux network can be extremely complicated, relatively simple long-term dynamics can emerge over a large timescale. For example, many biochemical reactions are complex nonlinear functions that depend on substrate and enzyme concentrations, yet the entire cell can follow a simple exponential growth at a large time scale.

Our goal is to characterize a mathematical class of flux networks that give rise to exponential dynamics in the long term. While a system evolves and increases its size, the relative proportion of each node also evolves. Since a growing process is not stationary, acquiring its long-term statistical averaging is generally not possible. Here, we explored the class of *scalable functions* (described in section 2) for constructing dynamical systems. The proportional scaling property allows us to decouple the *n*-dimensional dynamics into 1-dimensional “bulk dynamics” and (*n-1)*-dimensional “relative proportion dynamics”. The latter dynamics is confined in the (*n-1)*-dimensional unit simplex, which enables us to apply ergodic theory for long-term statistical averaging.

In section 1, we introduce the basic mathematical structure for flux networks and the concept of long-term growth rate. In section 2, we introduce *scalable reaction networks* and present the main result (Theorem 2.3). This result guarantees, via the use of ergodic theory, converged long-term growth rates for many (though not all) initial conditions of scalable networks. Examples are discussed. In section 3, we consider the addition of scalable noise and show that the presence of such noise, together with a regenerative condition on the networks, guarantees well-defined long-term growth rates for all initial conditions. Along with the theory, we investigate concrete examples to illustrate non-steady-state growth modes and rich behaviors (including chaotic dynamics) for growing reaction networks. In section 5, we generalize the class to *asymptotic scalable networks*, which replaces the stringent “proportional scaling” condition to a more relaxed “asymptotic scaling”. In section 6, we study the necessary structural motifs for the network to have a positive long-term growth rate.

#### 1. Flux network (*x*, *ϕ*, *J*) and long-term growth rate

As in thermodynamics formulations, we consider a *system* that is connected to an *environment*. The environment (denoted as *E*) acts as a reservoir and provides unlimited supply and removal of materials. The system is composed by multiple nodes *x*_1_, …, *x*_2_ and interconversion reactions *ϕ*_1_, …, *ϕ*_*m*_. A reaction can happen between the environment and system nodes, as well as among multiple system nodes. In general, a reaction *ϕ*_*a*_ can be represented by

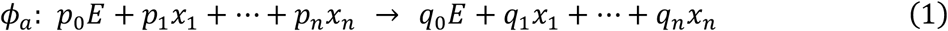

where *p*_*k*_ and *q*_*k*_ (*k* = 0, …, *n*) represent the *stoichiometry coefficients* in this conversion. Specifically, whenever the reaction happens, a proportional amount of *E*, *x*_1_, …, *x*_*n*_ is converted into another proportional amount of *E*, *x*_1_, …, *x*_*n*_, where both proportions follow the relative ratio of *p*_*j*_ and *q*_*j*_. It is clear that if *p*_*k*_ > *q*_*k*_, the reaction *ϕ*_*a*_ consumes *x*_*k*_ and decreases the amount of material on node *x*_*k*_.

The above description is simplified by classifying nodes that are increased, decreased, or unchanged during the conversion. If we denote *s*_*k*_ ≡ *p*_*k*_ − *q*_*k*_ and environment *E* ≡ *x*_0_, then the reaction *ϕ*_*a*_ can be rewritten as

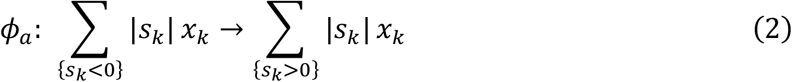

for *k* = 0,1, . ., *n*. Nodes *x*_*k*_ with *s*_*k*_ < 0 (or *s*_*k*_ > 0) are referred to be at the *upstream* (or *downstream*) of the reaction *ϕ*_*a*_. Conceptually, each conversion reaction can be regarded as an “action” that decreases the amount of its upstream nodes and increases the amount of its downstream nodes. This can be visualized as the *network diagram*:

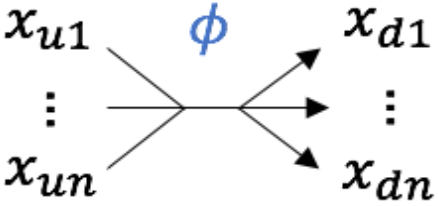

where {*x*_*u*_} and {*x*_*d*_} are the upstream and downstream nodes of the reaction *ϕ*, respectively. Note that the network diagram is more complex than the directed graph — it allows multiple upstream/downstream nodes.

A flux network can usually have multiple reactions {*ϕ*_1_, …, *ϕ*_*m*_}. The net stoichiometry coefficient of reaction *a* on node *k* is denoted by *S*_*ka*_. We define the *n*-by-*m stoichiometry matrix S* by [*S*]_*ka*_ = *S*_*ka*_. This way, each column of *S* represents the stoichiometry coefficient of one reaction. Here, the stoichiometry coefficient of environment (*x*_0_) is not included in *S*, since we assume unlimited supply/removal of materials from the environment.

##### Example

Consider the following flux network with *n* = 4 nodes and *m* = 3 reactions:

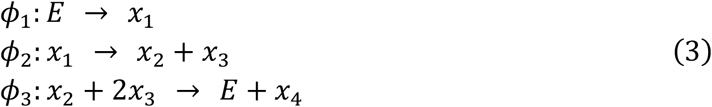

The stoichiometry matrix *S* and the network diagram are below:

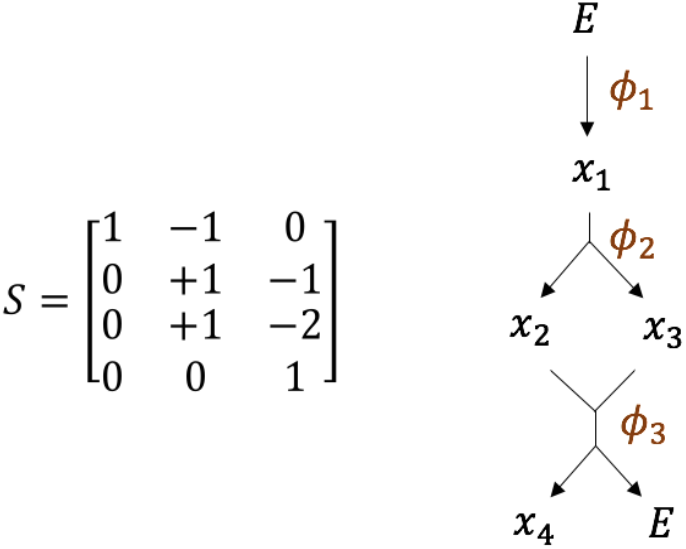

Above, we used the stoichiometry matrix *S*_*ka*_ to describe the topology of the reaction network. Each reaction *ϕ*_*a*_ is further associated with a flux function *J*_*a*_(*X*). The flux function can be linear such as *J*_*a*_(*X*) = *X*_1_ + *X*_2_ or nonlinear such as 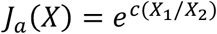. Below, *X*_1_, … *X*_*n*_ denote the *absolute amount* of material on nodes *x*_1_, …, *x*_*n*_, and *J*_1_, …, *J*_*m*_ denote the *flux* of conversion reactions *ϕ*_1_,. ., *ϕ*_*m*_.

##### Remark on notation

The *x*_*k*_ and *ϕ*_*a*_ are algebraic terms: *x*_*k*_ are node labels and *ϕ*_*a*_ are stoichiometry column vectors. They are used to describe the *network structure*. The *X*_*k*_ and *J*_*a*_(*X*) are multivariate nonnegative functions associated with *x*_*k*_ and *ϕ*_*a*_, respectively. They are used to model the *network dynamics*.

In a small time interval *Δt*, the reaction *ϕ*_*a*_ causes *X*_*k*_ to change by an amount of *S*_*ka*_*J*_*a*_*Δt*, where *S*_*ka*_ is the net stoichiometry coefficient of flux *a* on node *k*. Under a continuous time limit, the deterministic dynamics of a flux network follows a differential equation. Given a node *x*_*k*_, the rate of change of *X*_*k*_ is equal to the sum of all fluxes weighted by their stoichiometry coefficients on node *x*_*k*_:

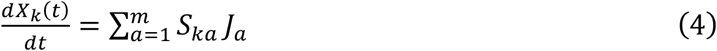

The flux magnitude *J*_*a*_ is usually a nonlinear function that depends on the amount or relative amount of materials in system nodes. In general, we write *J*_*a*_(*X*) where *X* = (*X*_1_, …, *X*_*n*_)^*T*^ is a column vector. For the general nonlinear function *J*_*a*_(*X*), there is no easy way to explicitly solve the equation. Our approach is to constrain the function form of *J*_*a*_(*X*) and study the long-term dynamics of the system.

Let *N* ≡ *X*_0_+… +*X*_2_ denote the total amount of material in the system, or the *system size*. We are interested in flux networks that have asymptotic exponential growth of *N*(*t*). We define long-term growth rate *λ* by the following limit (if it exists):

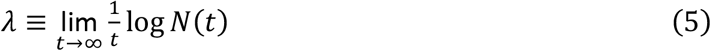

Note that for an arbitrary flux network, this limit may not exist. The asymptotic behavior of 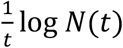 can be different, such as (1) *N*(*t*) becomes negative in the long term, and hence *λ* is not well-defined, (2) *N*(*t*) diverges at a finite time *t*_*c*_ and hence *λ* is not well-defined, and (3)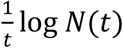 diverges to infinity as *t* → ∞, and (4) 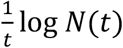 oscillates indefinitely and does not converge to a real number as *t* → *∞*. Therefore, it is clear that constructing flux networks with arbitrary flux functions may not give a well-defined long-term growth rate.

The long-term growth rate *λ* can be positive, negative or zero, which correspond to exponential growth, exponential decay, or other sub-exponential dynamics, respectively. The value of *λ* may also depend on the initial condition *X*(0) of the system. Each trajectory can have a different *λ*. We explicitly denote this by *λ*_*X*(0),_ if necessary.

Exponentially growing systems are often studied by starting with a system of equations 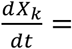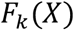 and assuming *a priori* that there exist (i) a long-term growth rate *λ* and (ii) a *balanced growth vector X*\* such that for *k* = 1, …, *n*

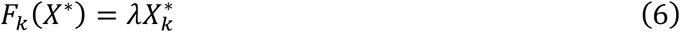

If the above equation has a solution (*λ*, *X*\*), then on the half-line {*X* = *cX*\*, *c* > 0} the system trajectory follows a *balanced growth solution X*(*t*) ~ *X*\* *e*^*λt*^. In biological models, the *λ* mentioned above is regarded as the *dilution rate* for the cells or the chemostat. Still, it is unclear how to find the proper *F*_*k*_(*X*) such that the above solution exists. To our knowledge, there are few empirical rules for constructing nonlinear, exponential growing networks. Moreover, the above method only identifies balanced growth solutions and no other types of growth dynamics, such as oscillations, which occur in many biological systems. In the following section, we focus on a class of flux functions *J*_*a*_(*X*) for which the convergence of *λ* can be rigorously analyzed.

#### 2. Scalable reaction networks

In this manuscript, we denote *Q*^*n*^: {*X* ∈ ℝ^*n*^ | *X*_*k*_ ≥ 0} as the non-negative quadrant and *Q*^*n*+^: {*X* ∈ ℝ^2^ ~ *X*_*k*_ > 0} as the positive quadrant. For all discussions, we will assume that the system *X*(*t*) has the initial condition *X*(0) ∈ *Q*^*n*^, since the amount of starting material (e.g., metabolites) should be nonnegative in living systems. Recall that a node *x*_*k*_ is *upstream* of flux *ϕ*_*a*_, if the stoichiometric coefficient *S*_*ka*_ < 0.

##### Definition 2.1

A flux function *J*_*a*_(*X*): *Q*^*n*^ → ℝ is *scalable* if it satisfies the following conditions:

(C1) (*differentiability*): *J*_*a*_(*X*) is positive in *Q*^*n*+^ and continuously differentiable in *Q*^*n*^ ∖ {0}.
(C2) (*upstream limited*): If node *x*_*k*_ is upstream of *ϕ*_*a*_, then *J*_*a*_(*X*) = 0 whenever *X*_*k*_ = 0.
(C3) (*proportional scaling*): *J*_*a*_(*cX*) = *cJ*_*a*_(*X*) for all scalar *c* > 0 and all *X* ∈ *Q*^2^.

##### Examples of scalable flux functions

Below, *x*_*j*_ represents an upstream node while *x*_*p*_ and *x*_*z*_ denote other nodes in the network, as exemplified here:

1. *rX*_*j*_, with *r* > 0.
2. 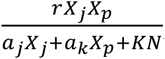, with *a*_*j*_, *J*_*k*_ ≥ 0, *r*, *K* > 0.
3. 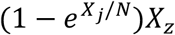
4. [*a* + *b sin*(*X*_*p*_/*N*)]*X*_*j*_, with *a* > *b* > 0.

Flux networks constructed by scalable functions are called scalable reaction networks (SRNs). SRNs have the nice property that the vector field satisfies the scaling property *F*_*k*_(*cX*) = *cF*_*k*_(*X*). Therefore, once we specify the vector field on the *n* − 1-dimensional unit simplex *Δ*^*n*-1^ ≡ {*X* ∈ *Q*^*n*^|*X*_1_+… +*X*_*n*_ = 1}, the entire vector field can be obtained from the scaling relation. Furthermore, as we will show below, this scaling property is closely related to the asymptotic exponential growth of the system.

To analyze the scalable network, we change variable from the *X*-coordinate to (*Y*, *N*)-coordinate where *N* = *X*_1_+… +*X*_*n*_ and *Y*_*k*_ = *X*_*k*_/*N* for *k* = 1, …, *n* (see Fig. S1). The vector *Y* = (*Y*_0_, …, *Y*_2_)^*T*^ is the *relative fraction* of each node in the system. In general, the differential equation 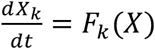 can be transformed to (*Y*, *N*)-coordinates as

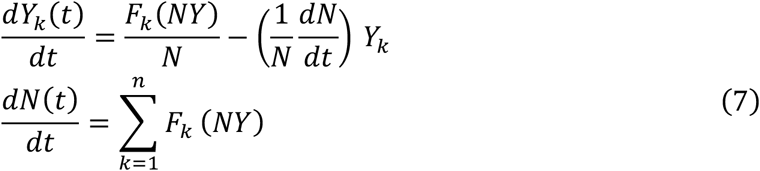

Next, we introduce the *instantaneous growth rate* of the system by 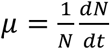. From the above equation, we have 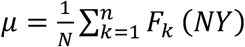. Now, we assume that all fluxes in the network are scalable. By the scaling property (C3), we have *F*_*k*_(*NY*) = *NF*_*k*_(*Y*) for all *k* = 1, …, *n*, and the above equation can be reduced to

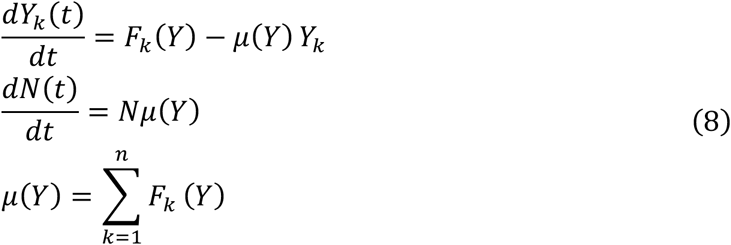

Note that the instantaneous growth rate *μ* now becomes a function of *Y* only and is independent of *N*. The dynamics of 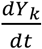 become a function of *Y* only and are independent of *N*.

Since *μ* is a function of *Y*, the dynamics of *μ*(*t*) is determined by the dynamics of *Y*(*t*). For scalable fluxes, the trajectory *Y*(*t*) is confined in the *n* − 1-dimensional unit simplex space *Δ*^*n*-1^ ≡ {*X* ∈ *Q*^*n*^|*X*_1_+… +*X*_*n*_ = 1}. To see this, note that the upstream-limited condition (C2) guarantees that whenever the upstream node of a flux is zero, the flux itself is also zero. Hence, the non-negative quadrant *Q*^*n*^ is forward invariant for 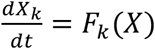. This implies that the unit simplex *Δ*^*n*-1^ is forward-invariant for 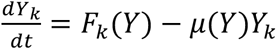.

Now, we can express the long-term growth rate as the *time average* of *μ*:

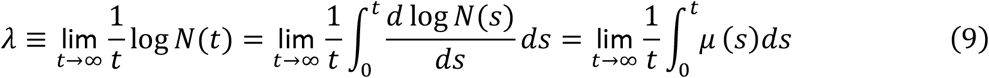

Our next step is to replace the time average by the *space average* on the space *Δ*^*n*-1^. To proceed, we need to introduce some concepts about ergodic theory. Below, we adopt the notation *F*(*X*) = (*F*_1_(*X*), … *F*_*n*_(*X*))^*T*^ as a vector function and denote *Y* = (*Y*_1_, …, *Y*_*n*_)^*T*^ as a vector on *Δ*^*n*-1^. We denote 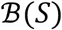 as the Borel sets of *S*, and write a probability measure *α*(*dY*) as *dα*.

###### Definition 2.2

Given a dynamical system *Y*′(*t*) = *G*(*Y*) on *S* ⊆ ℝ^*n*^, a Borel probability measure *ω* is called *invariant* if for every Borel set *A* ⊆ *S*, *ω*{*Y*(*t*) ∈ *A*} = *ω*{*Y*(0) ∈ *A*} for all *t* ∈ ℝ. An invariant probability measure *ω* is called *ergodic* if the system has no invariant subset of intermediate *ω*-measure, i.e., there does not exist *A* ⊆ *S* with 0 < *ω*(A) < 1 such that *Y*(0) ∈ *A* implies *Y*(*t*) ∈ *A* for all *t*.

We remark that every continuous dynamical system on a compact space admits at least one ergodic measure [Ref(43), Thm 1.8.1]. Given a dynamical system *Y*′(*t*) = *G*(*Y*) on *S* ⊆ ℝ^2^ and an ergodic measure *ω*, the ergodic theorem asserts that time averages are equal to space averages, as described below.

###### Birkhoff Ergodic Theorem

[Ref(43), Thm 1.2.1]. Let *Y*(*t*) and *ω* be as above. Then for *ω*-almost every *Y*(0) ∈ *S*, we have

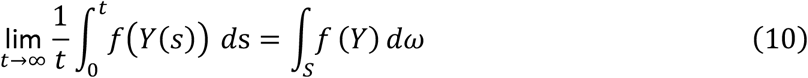

for every continuous function *f*(*Y*): *S* → ℝ.

A point *Y*(0) ∈ *S* is called *ω*–*regular*, or regular with respect to *ω*, if it satisfies Eq. (10) for every continuous function *f*(*Y*). We say *Y*(0) is regular if it is regular with respect to some ergodic measure. The following result is a direct consequence of Birkhoff Ergodic Theorem with *f*(*Y*) = *μ*(*Y*):

###### Theorem 2.3

Given a scalable reaction network, the long-term dynamics of *Y*(*t*) is governed by

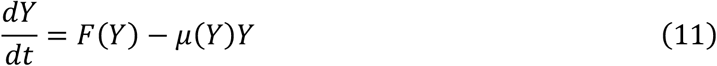

and is confined in *Δ*^*n*-1^. Furthermore, if *Y*(0) is regular with respect to an ergodic probability measure *ω*, then the long-term growth rate converges to

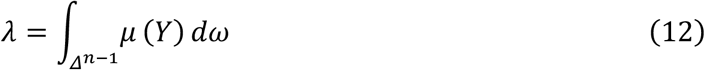

##### Note 1

Given an initial condition *Y*(0) that is *ω*-regular, the ergodic probability measure *ω* can be constructed as follows. Let χ_*B*_ be the *characteristic function* such that:

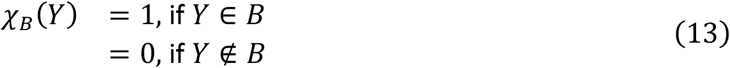

where *B* ⊆ *Δ*^*n*-1^ is a Borel set. We define a family of probability measures {*p*_*t*_}_*t*≥0_ by

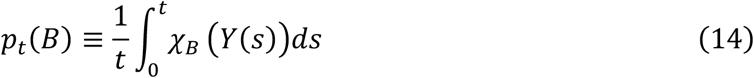

Intuitively, *p*_l_(*B*) is the fraction of time that *Y*(*t*) stays within *B* during [0, *t*]. Then, as *t* tends to infinity, *p*_*t*_ converges in the weak* topology to *ω*, i.e., for every continuous function *f*(*Y*): *S* → ℝ, the ergodic probability measure *ω* is the weak* limit such that

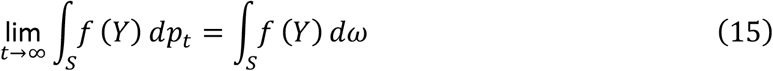

Conversely, for any initial condition *Y*(0), if *p*_*t*_ converges to an ergodic measure *ω*, then *Y*(0) is *ω*-regular. We note, however, that *p*_*t*_ may not converge for some *Y*(0), and that the limit measure (even when it exists) need not be ergodic and may depend on *Y*(0).

##### Note 2

If 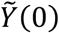 is another initial condition with 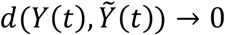 as *t* → ∞, then the growth rate at 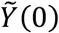 is also well-defined and equal to *λ*.

##### Note 3

It is possible that for some initial conditions (within a set of *ω*-measure zero) the growth rate does not converge to the value given by Eq. 12. Here, we analyze a deterministic example to illustrate this possibility.

We consider a deterministic case on the simplex space *T*^*n*-1^ with *n* = 3 (see figure at right). The simplex space contains two fixed points: *Y*_*a*_ is unstable and *Y*_*b*_ is stable. For this system, we find two ergodic measures on *T*^2^: the delta measures *δ*(*Y* − *Y*_*a*_) ≡ *ω*_*a*_ and *δ*(*Y* − *Y*_*b*_) ≡ *ω*_*b*_. Since *Y*_*b*_ is a stable attractor, all trajectories with the initial condition *Y*(0) ≠ *Y*_*a*_ converge asymptotically to the constant trajectory *Y*(*t*) = *Y*_*b*_ and therefore have *λ* = *μ*(*Y*_*b*_). The trajectory with *Y*(0) = *Y*_*a*_ has *λ* = *μ*(*Y*_*a*_). Thus, if *μ*(*Y*_*a*_) ≠ *μ*(*Y*_*b*_) then with respect to the ergodic measure *ω*_*b*_ we have that {*Y*_*a*_} is a set of measure zero for which *λ* does not converge to the value given by Eq. 12; and the same is true, respectively, for the ergodic measure *ω*_*a*_ and the set *T*^2^\{*Y*_*a*_}.

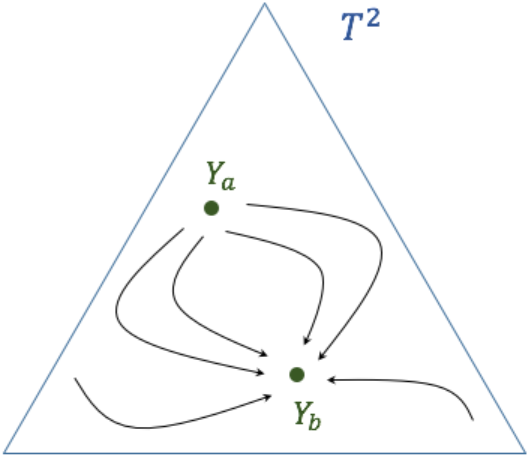

This discussion, together with Theorem 2.3, shows that for a scalable network, many initial conditions *X*(0) are guaranteed to have well-defined long-term growth rates. This includes all *X*(0) for which the rescaled initial conditions *Y*(0) are regular.

We now consider a few illustrative examples:

1. For a flux network with *Y*(*t*) converging to a fixed point *Y*\*, the ergodic probability measure is a Dirac *δ*-measure supported on *Y*\*. The ergodic averaging for the long-term growth rate is reduced to *λ* = *μ*(*Y*\*). The long-term dynamics converges to balanced exponential growth on the half line {*X* = *cY*\*, *c* > 0}, with *X*(*t*) → *Y*\**e*^*μ*(*Y*\*)*t*^.
2. For a flux network with *Y*(*t*) converging to a limit cycle *C*, the ergodic probability measure is supported by the limit cycle. The long-term growth rate can be calculated by the contour integral *λ* = ∮_*c*_ *μ* (*Y*) *dω*, and the dynamics converges to exponential growth with periodic functions *u*_*k*_(*t*) as prefactors, namely, *X*_*k*_(*t*) → *u*_*k*_(*t*)*e*^*λt*^.
3. There are attractors of *Y*(*t*) that *do not* have an ergodic probability measure. One example is the heteroclinic cycle attractor, where the trajectory *Y*(*t*) oscillates between *M* saddle points {*Y*_*s*1_, …, *Y*_*sM*_} with an increasing period. In this case, *λ* oscillates between the values {*μ*(*Y*_*s*1_), …, *μ*(*Y*_*sM*_)} with an increasing period and does not converge to a real number.

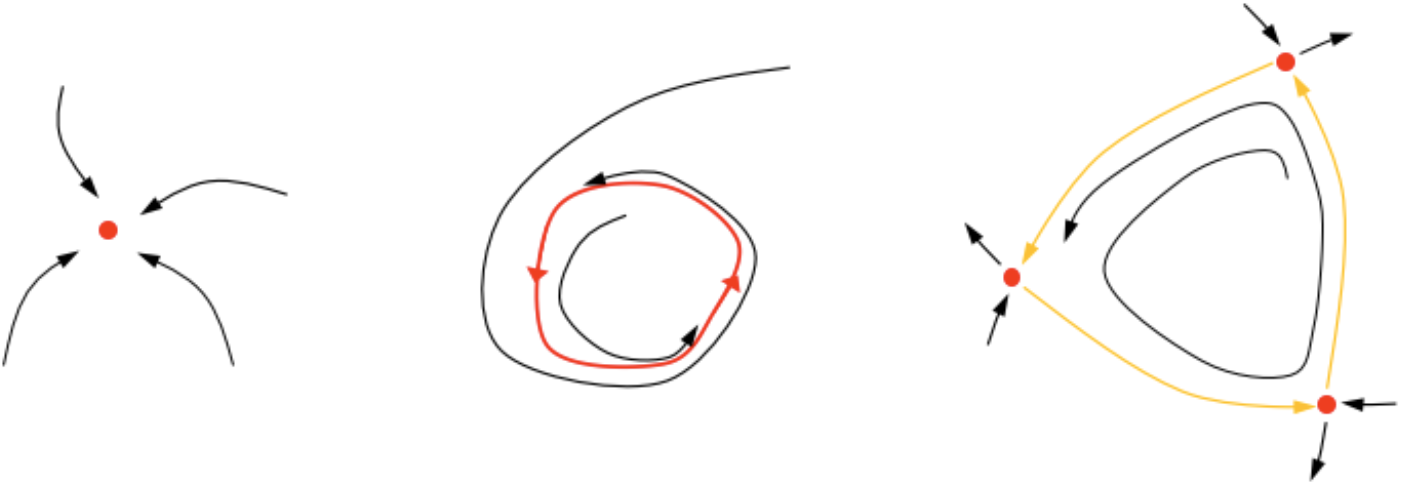

Different types of attractors and invariant measures. Red color indicates invariant sets in the system. Left: a stable fixed point attracting all nearby trajectories. Middle, a stable limit cycle attracting all nearby trajectories. Right, a heteroclinic cycle (union of red saddle points and yellow curves). The saddle points support ergodic measures. For the trajectory *Y*(*t*) shown, the measures *p*_l_ in the construction above either do not converge or converge to a measure that is a linear combination of the ergodic measures supported on the three saddles. Such a measure is not ergodic. The initial condition *Y*(0) is not regular and does not have a well-defined long-term growth rate.

As we have seen, balanced exponential growth belongs to one of the special (and simplest) types of exponentially growing networks. Additional growth modalities exist, which correspond to different types of attractors of *Y*(*t*), as shown in the following two examples of SRNs.

###### Example 2.4: The autocatalytic repressilor network

In this example, we modify the repressilator model by Elowitz and Leibler [Ref(44)] to be an autocatalytic, scalable network. The system has four nodes, where *x*_1_ represents the upstream metabolite and *x*_2_, *x*_3_, *x*_4_ represent three mutually repressing components:

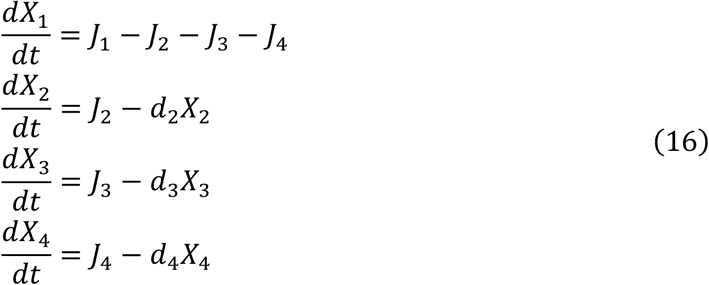

In the repressilator, the synthetic flux of nodes *x*_2_, *x*_3_, *x*_4_ are repressed by node *x*_3_, *x*_4_, *x*_2_, respectively. We implemented this by defining the synthetic flux function *J*_2_, *J*_3_, *J*_4_ as decreasing functions of *X*_2_, *X*_3_, *X*_4_. With *Y*_*k*_ denoting the fraction of *X*_*k*_, the flux functions are: 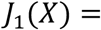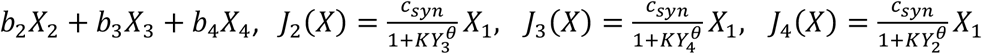.

The autocatalytic repressilator network can exhibit different growth modes, depending on the cooperativity coefficient *θ* (see Fig. 3A-E in main text). For *θ* = 1, *Y*(*t*) converges to a stable fixed point, while for *θ* = 3, *Y*(*t*) converges to a limit cycle, which corresponds to a periodic growth mode.

###### Example 2.5: Modified May-Leonard system

In this example, we study density-dependent growth of three species, which illustrates a case where *Y*(*t*) is a heteroclinic cycle and *Y*(*t*) is not regular with respect to any ergodic probability measure. The model is modified from the classical May-Leonard model [Ref(45)], with scalable flux functions as birth and death processes. Let *x*_1_, *x*_2_, *x*_3_ denote the three species, the growth equation is

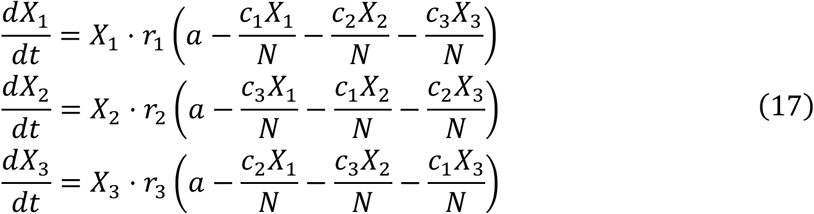

This system can be mapped to a network diagram by defining 12 scalable flux functions (see Fig. S2a), three for the birth process and nine for the death process. The death process is driven by a nonlinear flux function in the form of (*X*_j_*X*_*k*_)/*N*.

In our simulation, we find that the trajectory *Y*(*t*) of the flux network exhibits similar behavior as in the original May-Leonard system. Consistent with the analytical results of the May-Leonard system, varying *c*_2_ and *c*_3_ changes the system behavior drastically (see Fig. S2):

1. For *c*_2_ + *c*_3_ < 2 (Fig. S2B), the system has balanced growth and three species co-exist.
2. For *c*_2_ > 1 and *c*_3_ > 1 (Fig. S2C), species *X*_1_ is dominant and grows exponentially, while the other species *X*_2_ and *X*_3_ go extinct. The system still undergoes balanced growth, with fixed point *Y*\*(*t*) = (1,0,0) on the boundary of *Δ*^2^.
3. For *c*_2_ + *c*_3_ > 2 and *c*_2_ < 1 < *c*_3_ (Fig. S2D), *Y*(*t*) asymptotically converges to a heteroclinic cycle on the boundary of the system. The heteroclinic attractor *C* ≡ {*Y* ∈ *Δ*^2^|*Y*_1_*Y*_2_*Y*_3_ = 0} is the omega-limit set of *Y*(*t*), but *Y*(*t*) is not regular with respect to any ergodic probability measure. In this case, each component *Y*_*k*_(*t*) oscillates between 0 and 1 while each period becomes exponentially longer. On the other hand, the overall system *X*(*t*) grows unboundedly without the long-term growth rate *λ* converging to a real number (see Remark (3) in Proposition 2.4).

###### Definition 2.6

A *quasi-periodic trajectory Y*(*t*) with *q frequencies* is a trajectory that is parametrized by *Y*(*t*) = *h*(*A*(*θ*_1_*t*, …, *θ*_*q*_*t*)), where *h* is a continuous function from torus *T*^*q*^ to *S*, and *A*: *T*^*q*^ → *T*^*q*^ is a continuous map between torus *T*^*q*^ with rationally independent frequencies {*θ*_1_, …, *θ*_*q*_} (i.e., 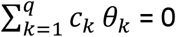 has no integer solution except *c*_1_ =… = *c*_*k*_ = 0).

###### Proposition 2.7

Consider an SRN with the initial condition *X*(0) ∈ *Q*^*n*^. If the trajectory of *Y*(*t*) converges to a fixed point, a periodic cycle, or an invariant torus with quasi-periodic dynamics, then *Y*(0) is regular for some ergodic probability measure.

*Proof:* For *Y*(*t*) converging to a fixed point *Y*\* the ergodic probability measure is *δ*(*Y* − *Y*\*). For the q-frequency torus (including *q* = 1 case for periodic trajectory), the trajectory is parametrized by equation *Y*(*t*) = *A*(*θ*_1_*t*, …, *θ*_*q*_*t*), which follows

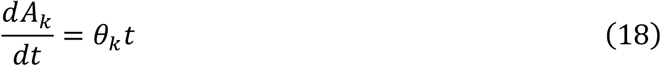

with *k* = 1, …, *q*. The flow *A*(*t*) on the torus *T*^*q*^ is uniquely ergodic with respect to Lebesgue measure on *T*^*q*^ [Ref(43), Thm 3.1.2], Hence *A*(*t*) has an ergodic probability measure on *T*^*q*^. Since both *h* and *A*(*t*) are continuous, this implies *Y*(*t*) also has an ergodic probability measure on *Δ*^*n*-1^.

###### Example 2.8: SRN-based models for chemostat-type systems

In this section, we study the relation between the SRN framework and chemostat-type equations. The general SRN framework assumes unlimited external resources, which conflicts with chemostat conditions in which an essential nutrient is in limited amount. However, for a chemostat equation, there exists an equivalent SRN model that gives rise to the same dynamics. Below, we use classical chemostat equations [Ref(46)] for demonstration.

Let *s*(*t*) and *x*(*t*) represent the concentrations of an upstream-limited nutrient and the microorganism in the chemostat culture vessel, respectively. The dynamics of *s*(*t*) and *x*(*t*) follows the chemostat equations:

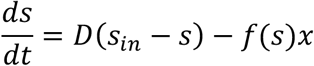

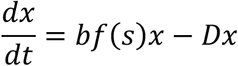

where *D* is the dilution rate of the culture, *s*_*in*_ is the constant level of the external nutrient supply, *b* is the stoichiometry constant for the biomass yield, and 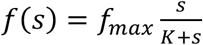 is the Monod’s function, which describes the dependence of the nutrient uptake rate (influx) on the extracellular nutrient level *s*(*t*).

The above chemostat equations are not scalable (e.g. *f*(*s*) does not satisfy *f*(*cs*) = *cf*(*s*)). However, since the equations are formulated in terms of concentrations *s*(*t*) and *x*(*t*), the system dynamics should be independent of the chemostat volume. Intuitively, scaling the chemostat volume to different sizes should yield identical dynamics of concentrations. Therefore, if we reformulate the equation in terms of the total number of molecules, the system should be an SRN.

To rigorously show this, we include the solvent molecule species *w* in the equations. Let *S*, *X*, and *W* be the total numbers of nutrient molecules, cells, and solvent molecules in the system, and let τ_*S*_, τ_*X*_, τ_*W*_ be their unit volumes. The <u>total culture volume</u> *V* > 0 is a time-independent positive constant and is given by:

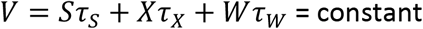

This imposes a constraint among *S*, *X*, and *W*, and we have the relations *s* = *S*/*V*, *x* = *x*/*V*, and *w* = *W*/*V*. We refer to the quantities (*s*, *x*, *w*) as *intensive variables* and (*S*, *X*, *W*) as *extensive variables*, similar to thermodynamics definitions.

Since *V* is a constant, we have

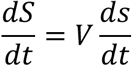

Note that the variable 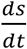, which is a function of intensive variables (*s*, *x*), is scale-invariant under scaling (*S*, *X*, *W*) → *c*(*S*, *X*, *W*). To see this, note that

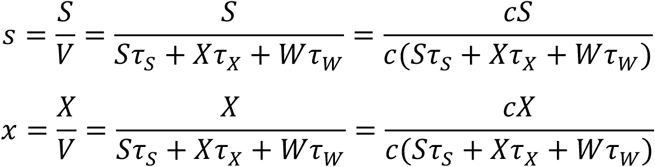

Therefore, *S*′(*t*) = *V s*′(*t*) is a scalable function. Similarly,

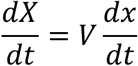

The variable 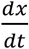, which is a function of intensive quantities (*s*, *x*), is also scale-invariant under scaling (*S*, *X*, *W*) → *c*(*S*, *X*, *W*). Therefore, *X*′(*t*) = *V x*′(*t*) is a scalable function.

For the solvent variable *W*(*t*), by the constraint of constant total volumes *V*, we have

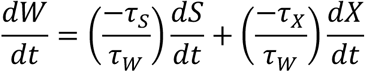

Since both *S*′(*t*), *X*′(*t*) are scalable, *W*′(*t*) is also scalable. Therefore, the ODE of the new variables (*S*, *X*, *W*) is an SRN and is equivalent to the chemostat equations. The same mapping can be applied for studying more complex chemostat dynamics, including multi-species interactions (as shown in Fig. 4E and described in SI in the Method section).

##### Remark

For the above equivalent SRN model mentioned above, the long-term growth rate of the system (*S*, *X*, *W*) is zero. This is due to the constraint that the total volume 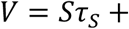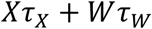 is a positive constant, and hence it is impossible to have *λ* > 0 (*V* diverges to infinity) or *λ* < 0 (*V* converges to 0).

The SRN framework provides additional insight into how biomass growth rate, *bf*(*s*), is related to the dilution rate *D*. We define

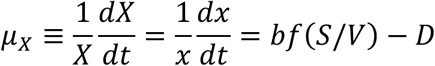

If the dilution rate is higher than the maximal biomass growth rate (i.e., *D* > *bf*_*max*_), then *μ*_c_ < 0 and all cells in the chemostat will be washed away. For cells to persist in the chemostat culture, the time average of *μ*_*X*_ must equal *λ*. For the equivalent SRN model, we have *λ* = 0 as mentioned above. Using the SRN formulation, we integrate *μ*_*X*_ with respect to an ergodic measure *ω*, which yields

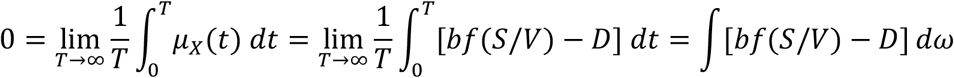

and hence

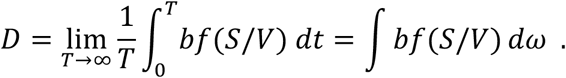

In general, the constant dilution rate *D* is equal to the average biomass growth rate *bf*(*S*/*V*) with respect to an ergodic measure *ω*. For balanced growth (as a special case), we have *D* = *bf*(*S*\*/*V*\*) where (*S*\*, *X*\*, *W*\*) is the fixed point of the system. The same relation holds for more complex chemostat dynamics. For example, when a multi-species chemostat exhibits oscillatory behavior, the limit cycle has the property that the average biomass growth rate on the limit cycle is equal to the dilution rate. This also applies for dynamics which include a limit torus or chaos.

#### 3. Growth rates in scalable reaction networks with noise

For scalable reaction networks, ergodic theory offers a conceptual understanding of long-term growth rates. Because the rescaled system takes values in a compact region, long-term growth rates are guaranteed to exist for many initial conditions, namely those that are regular with respect to some ergodic probability measure. Given a particular initial condition, however, growth properties are not always easy to determine, because not every initial condition is regular and more complicated systems often have multiple ergodic measures each with their own set of regular points and growth rates. This less-than-ideal state of affairs is improved greatly in the presence of random noise. Real-world systems are inherently noisy and mathematically stochastic terms are often added to model uncontrolled fluctuations in physical and biological systems. As we will show in this section, the averaging effect of random noise leads to a simpler and more tractable dynamical picture.

To investigate how flux noise affects the system’s behavior, we generalize the ordinary differential equation 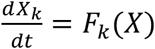 into a stochastic differential equation (SDE). As before, we let *Q*^*n*^: {*X* ∈ ℝ^*n*^ | *X*_*k*_ ≥ 0}, and *Q*^*n*+^: {*X* ∈ ℝ^*n*^ | *X*_*k*_ > 0}.

##### Definition 3.1

A *scalable reaction network with scalable noise* is a systems SDE

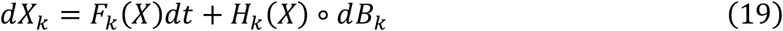

for *k* = 1, …, *n*. Let *S* denote the constant *stoichiometry matrix* and define the deterministic flow by *F*_*k*_(*X*) = ∑_*a*_ *S*_*ka*_ *J*_*a*_(*X*). Every flux *J*_*a*_ needs to satisfy the following properties:
(C1) *J*_*a*_(*X*) is positive in *Q*^*n*+^ and continuously differentiable in *Q*^*n*^ ∖ {0}.
(C2) If the node *x*_*k*_ is one of the upstream nodes of *ϕ*_*a*_, then *J*_*a*_(*X*) = 0 whenever *X*_*k*_ = 0.
(C3) *J*_*a*_(*cX*) = *cJ*_*a*_(*X*) for arbitrary scalar *c* > 0 for all *X* ∈ *Q*^*n*^.

The noise function *H*_*k*_(*X*) satisfies:

(N1) *H*_*k*_(*X*) > 0 on {*X* ∈ ℝ^2^ | *X*_*k*_ > 0} and twice continuously differentiable in *Q*^*n*^ ∖ {0}.
(N2) *H*_*k*_(*X*) = 0 whenever *X*_*k*_ = 0.
(N3) *H*_*k*_(*cX*) = *cH*_*k*_(*X*) for arbitrary *c* > 0.

Here, *dB*_*k*_ represents the standard Brownian motion, with integral in Stratonovich-sense.

In the above SDE, the deterministic part of the equation, i.e., *F*(*X*), has identical requirements as in scalable flux functions. We consider a noise function *H*_*k*_ which also scales with the system size. This is motivated by the following reasons:

i. In nature, the environment is never truly static. For example, the environmental temperature has nonzero fluctuations and this globally affects all enzyme activities. Consider a flux *J* = *rX*_*k*_ where the rate constant *r* = *r*(*T*) is a function of temperature *T*. Fluctuations of the environmental temperature *δT* create noise *δr* ≈ *r*′(*T*)*δT* on the rate constant *r*. Hence, this creates noise on flux *δJ* ≈ (*δr*)*X*_*k*_ = (*δr*/*r*)*J*. This noise scales with the flux magnitude and therefore scales with system size.
ii. Consider a system with finite space, for example, a bacterial culture in a turbidostat, the population is kept at a constant size *C* while the excess of growing bacteria is discarded by medium dilution. The dilution process usually creates a sampling noise 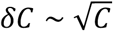. In this case, the growth dynamics is equivalent to an exponentially growing population with *N*/*C* copies of subsystem with size *C*. If the exponential growing population has size *N* and a noise level 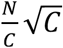, then each subsystem has a fluctuation 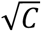. In this case, the equivalent exponentially growing population has a noise level that scales with system size.

In addition to scalable noise, our formulation allows a system with *λ* > 0 to have other types of noise that grow slower than the system size. These “sub-scalable noises” will decay to zero as the system grows to infinity. Therefore, only the scalable noise term remains.

To establish the existence and uniqueness of solutions for all *t* > 0 for the SDE in Eq. (19), we convert our SDE from Stratonovich to the Ito formulation (see [Ref(47)]). Since *H*(*X*) is a diagonal matrix, hence (*H*)_*kl*_ = *δ*_*kl*_*H*_*k*_, this simplifies to

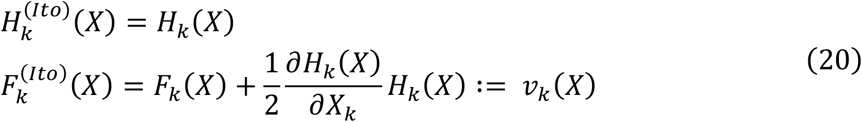

Existence and uniqueness are then checked by appealing to classical results (e.g. [Ref(47)], Theorem 5.6.1). Details are given in the Appendix.

As in the deterministic case, we now consider the rescaled SDE *Y*(*t*) defined by *Y*_*k*_ ≡ *X*_*k*_/|*X*|, |*X*| = *X*_1_+… +*X*_*n*_. We begin with a result asserting that starting from an initial condition where *X*_*k*_ > 0 for all *k*, i.e., all the substances in the network are present in positive amounts, no substance will be depleted in finite time.

##### Proposition 3.2

The trajectory *Y*(*t*) of Eq. (19) is confined in the unit simplex *Δ*^*n*-1^. For each component *k*, if *Y*_*k*_(0) > 0, then *Y*_*k*_(*t*) > 0 for all *t* > 0.

*Proof:* One way to prove this is to produce a Lyapunov function *V* on the interior of *Δ*^*n*-1^ with *V*(*Y*) → ∞ as *Y* tends to ∂*Δ*^*n*-1^, the boundary of *Δ*^*n*-1^. The function *V* is to have the property 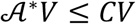 for some fixed *C* > 0 where 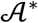 is the diffusion operator of the SDE; it controls the speed with which solutions can approach the boundary. A complete proof is given in the Appendix.

The SDE (19) generates a time-homogeneous Markov process on *Δ*^*n*-1^. For *Y*_0_ ∈ *Δ*^*n*-1^ and *s* > 0, we use *P*_*s*_(*Y*_0_, ∙) to denote the *transition probability* starting from *Y*_0_, that is, if *B* ⊆ *Δ*^*n*-1^ is a Borel set, then *P*_*s*_(*Y*_0_, *B*) is the probability that *Y*(*s*) ∈ *B* given that *Y*(0) = *Y*_0_. A probability measure *ω* is said to be *stationary* if *ω*(*B*) = ∫ *P*_*s*_(*Y*, *B*)*ω*(*dY*) for all Borel sets *B* and all *s* > 0. A stationary probability measure *ω* is called *ergodic* if there is no subset *A* ⊂ *Δ*^*n*-1^ with 0 < *ω*(*A*) < 1 such that *P*_*s*_(*Y*, *A*) = 1 for *Y* ∈ *A* and *s* > 0. The Ergodic Theorem in section 2 applies in this setting as it does in the deterministic case.

##### Corollary 3.3

(*Dichotomy for ergodic measures*): Consider an SRN with noise as defined by Eq. (19), and let *ω* be an ergodic probability measure of the rescaled system. Then we have either *ω*(∂*Δ*^*n*-1^) = 0 or 1.

*Proof*: This follows immediately from Proposition 3.2, which says that the interior of *Δ*^*n*-1^ is an invariant set, so it must have measure 0 or 1 by the ergodicity of *ω*.

Below, we give a sufficient condition for the existence of long-term growth rates for noisy scalable networks. Following the notation in Eq. (20), we call a system *regenerative* if for every *k* = 1, …, *n*, the *k*th component of the drift vector *v*_*k*_(*Y*) > 0 whenever *Y*_*k*_ = 0. Mathematically, this means the drift term in the SDE points into the interior of the simplex *t*^*n*-1^ at all points on its boundary.

##### Theorem 3.4

Consider an SRN with noise as defined by Eq. (19) and assume that the system is regenerative. Then, there is a number *λ* with the property that starting from any initial condition (0) ∈ *Δ*^*n*-1^, the long-term growth rate converges almost surely to *λ*.

*Proof*: First, ergodic stationary measures exist because the process is confined to a compact region and the transition probabilities *P*_*s*_(*Y*,∙) vary continuously with *Y*. Let *ω* be an ergodic probability measure. By Corollary 3.3, *ω*(∂*Δ*^*n*-1^) = 0 or 1. The regenerativeness of the network implies *ω*(∂*Δ*^*n*-1^) = 0, because the positivity of *v*_*k*_ pushes all initial conditions in {*Y*_*k*_ = 0} into the interior of *Δ*^*n*-1^ immediately. Next we observe that *ω* has a density, i.e., *ω*(*dY*) = ρ(*Y*)*dY* for some integrable function ρ(*Y*). This is because *ω* = ∫ *P*_*s*_(Y,∙)*ω*(*dY*) and *P*_*s*_(Y,∙) has a density for all *Y* in the interior of *Δ*^*n*-1^ by condition (N1) above. Observe that ρ(*Y*) > 0 on all of *Δ*^*n*-1^ by standard properties of diffusions. The uniqueness of *ω* follows, as all ergodic probability measures have strictly positive densities and distinct ergodic measures are supported on disjoint sets. Finally, the Birkhoff Ergodic Theorem applied to the function *μ*(*Y*) gives the result that the long-term growth rate of every *Y*(0) ∈ *Δ*^*n*-1^ exists almost surely and is equal to *λ* = ∫ *μ*(*Y*)*ω*(*dY*).

We remark that even though the invariant density ρ(*Y*) is positive on all of *Δ*^*n*-1^, it is in general much more concentrated in some regions of *Δ*^*n*-1^ than in other regions, that is to say, certain network configurations (in terms of proportions of the substances present) are much more prevalent even though there is a small theoretical probability of exploring other parts of the phase space. What makes the situation in Theorem 3.4 more tractable than the noise-free case is the uniqueness of ergodic measure and the fact that every initial condition is regular with respect to the unique ergodic measure.

##### The general case (of not-necessarily-regenerative networks)

Not all realistic biological networks are regenerative. Not being regenerative means that there is no explicit mechanism in place for each substance to restore itself as it heads towards relative depletion. To be precise, relative depletion here means the rescaled amounts *Y*_*k*_(*t*) (but not necessarily the actual amounts *X*_*k*_(*t*)) become arbitrarily small. Ergodic measures always exist for the same reasons as before, but unlike the regenerative case, there may exist ergodic measures supported on ∂*Δ*^*n*-1^. For example, if *v*_*k*_ ≡ 0 on {*Y*_*k*_ = 0}, then starting from *Y*(0) with *Y*_*k*_(0) = 0, almost surely *Y*_*k*_(*t*) = 0 for all *t* > 0 (see condition (N2)), so there is a subsystem defined on the (n-2)-dimensional simplex *Δ*^*n*-1^ ∩ {*Y*_*k*_ = 0} with its own ergodic measures.

In this general case, let us consider only initial conditions starting from the interior of *Δ*^*n*-1^. There is the following dichotomy:

*Case 1.* Existence of an ergodic measure with a density ρ(*Y*) > 0 on all of *Δ*^*n*-1^. In this case, long-term growth rates of all *Y*(0) in the interior of *Δ*^*n*-1^ converge almost surely to *λ* = ∫ *μ*(*Y*)ρ(*Y*)*tY*, as before.
*Case 2.* All ergodic measures are supported on ∂*Δ*^*n*-1^. In this case, starting from *Y*(0), in the interior of *Δ*^*n*-1^, one or more of the substances will come close to relative depletion (though none can deplete in finite time according to Proposition 3.2). More precisely, given any *Y*(0), for any *ε* > 0 we have Prob{*t*(*Y*(*t*), ∂*Δ*^*n*-1^) > *ε*} → 0 as *t* → ∞. We summarize our findings below, leaving mathematical proofs to be published elsewhere.

Since trajectories spend nearly 100% of their time near ∂*Δ*^*n*-1^, their large-time behaviors are dictated by the ergodic measures supported on ∂*Δ*^*n*-1^. Now ∂*Δ*^*n*-1^ is a finite union of (n-2)-dimensional simplices and an analysis similar to that for *Δ*^*n*-1^ can be made for each of these simplices. After successively reducing dimension, we arrive at the conclusion that i) the system has at most a finite number of ergodic invariant measures, each supported on a lower dimensional simplex and ii) in this lower dimension, the system has a density with respect to the Lebesgue measure.

It may happen that *Y*(*t*) converges almost surely to one of these ergodic measures *ω*, in which case its long-term growth rate is given by *λ* = ∫ *μ*(*Y*)*ω*(*dY*). Another possibility is for *Y*(*t*) to explore multiple ergodic measures one after another (in a way similar to the heteroclinic loop example in section 2). In this case, there may not be a single number that correctly describes the trajectory’s long-term growth rate.

#### 4. An exponentially growing system with chaotic dynamics

In the following example, we study a flux network converging to a nonperiodic trajectory and investigate the effect of noise on its long-term growth rate.

##### The double repressilator network

We construct a flux network with two repressilators repressing each other (see Fig. 2F and Fig. S3). The double repressilator network consists of seven nodes: *x*_1_ represents the common precursor, *x*_2_, *x*_3_, and *x*_4_ form the first repressilator, and *x*_5_, *x*_6_, and *x*_7_ form the second one. In addition, the two repressilators are coupled by mutual repression, where nodes *x*_2_ and *x*_5_ repress the influxes of each other. The full differential equation is:

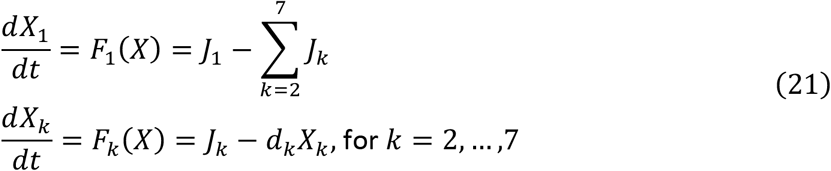

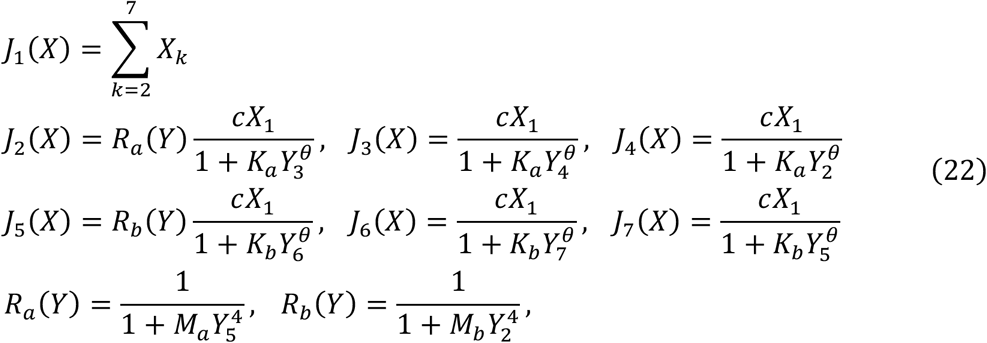

Note that this flux network satisfies the regenerative condition in Theorem 3.4. The parameters *K*_*a*_ and *K*_*b*_ are the repression strengths of repressilators {*x*_2_, *x*_3_, *x*_4_} and {*x*_5_, *x*_6_, *x*_7_}, respectively. We define *α* ≡ *K*_*a*_/*K*_*b*_ as the relative repression strength between the two repressilators. By varying *α*, the system exhibits complex behaviors, including periodic, quasi-periodic, and non-periodic trajectories of *Y*(*t*). For different values of parameter *α*, the omega-limit set of the *Y*(*t*) can have complex geometry such as a limit torus (*α* = 20), a complicated limit cycle (*α* = 100) and a strange attractor (*α* = 500) (Fig. 2G).

To show how the attractor of *Y*(*t*) transitions from a limit cycle to a limit torus to a more complex attractor, we project the attractor on the *Y*_2_ − *Y*_3_ plane and get a Poincare section at {*Y*_2_ = 0.2}. For various *α*, we see that the system exhibits rich behavior, including period doubling and transition to chaos (Fig. 2G, insets). For the full bifurcation diagram, see Fig. S3A.

To quantitatively analyze the attractor, we numerically calculated the largest Lyapunov exponent (LLE) of the trajectories Fig. S3B). LLE quantifies how small perturbations propagate along the solution trajectory, and is a useful indicator for chaotic dynamics: positive LLE indicates strange attractor while LLE=0 indicates a limit torus, a limit cycle or a fixed point. We found that for *α* with values between 400-500, LLEs are significantly positive, which is consistent with what is qualitatively observed in the bifurcation diagram. There are also sporadic “chaotic islands” for *α* = 120.

For comparison, we introduce a linear noise *H*_*k*_(*X*) = *h*_*k*_*X*_*k*_ on the SDE. We find that the LLEs with and without noise (black versus light blue lines in Fig. S3B) are comparable, indicating that the chaotic behavior is robust under perturbation. Although it is difficult to show the convergence of *λ* in a nonperiodic, deterministic system, there is noise associated with fluxes in most realistic systems. By introducing small noise, we can infer whether the long-term behavior observed in the deterministic system is robust. By Theorem 3.4, *λ* converges under arbitrarily small *h*_*k*_. We simulate long-term growth rate with *h*_*k*_ = 0 and *h*_*k*_ = 0.1 for various *α* (Fig. S3C). For non-chaotic regimes, noise has little or no effect on *λ*, while at the border between periodic and chaotic regimes (*α*~420), *λ* substantially differs whether noise is present or not. The noise seems to affect the transition between different types of attractors.

The chaotic growing system indicates that an exponentially growing system can have a complex trajectory that is unpredictable in the long-term. Still, the long-term growth rate of the system converges. We conclude that while the trajectory is sensitive to small perturbations, the long-term growth rate, which is a statistical averaging on the projected system, can be still be robust.

#### 5. Asymptotically scalable reaction networks

The condition (C3) of scalable flux requires that the flux function satisfies *J*(*cx*) = *cJ*(*x*) for a scalar *c* > 0. This is a relatively strong condition that reduces the degrees of freedom of the function by one dimension. Here, we show that a more general class of flux functions, referred to as *asymptotically scalable functions*, gives a similar convergence property on long-term growth rate.

##### Definition 5.1

A flux function *J*_*a*_(*x*): *Q*^*n*^ → ℝ is *asymptotically scalable*, if:

(C1)(*differentiability*): *J*_*a*_(*X*) is positive in *Q*^*n*+^ and continuously differentiable in *Q*^*n*^ ∖ {0}.
(C2)(*upstream limited*): If the node *x*_*k*_ is one of the upstream nodes of *ϕ*_*a*_, then *J*_*a*_(*X*) = 0 whenever *X*_*k*_ = 0.
(C3’)(*asymptotic scaling*): For every scalar *c*_1_ and *c*_2_ > 0 and every vector *x* ∈ *Q*^*n*^,

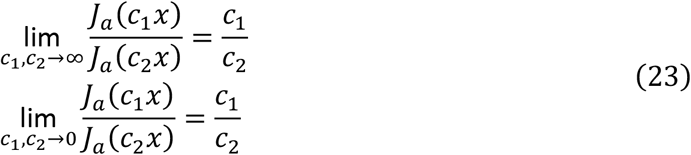

Note that the (C1) and (C2) conditions are identical to the scalable function definition, while condition (C3’) is more general than condition (C3).

##### Examples of asymptotic scalable flux functions

(*x*_*j*_ denotes the upstream node)

1. 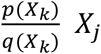, where *p*(*X*), *q*(*X*) are polynomials *c*_*i*_*X*^*i*^+… +*c*_*n*_*X*^*n*^ with *c*_*k*_ > 0 for *k* = *i*, …, *n*.
2. 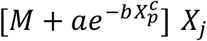, with *M*, *a*, *b* > 0, *c* ≥ 1.
3. [*M* + tan^−1^(*aX*_*k*_)] *X*_*j*_, with *M* > π/2.
4. 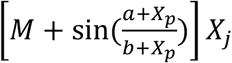, with *M* > 1, *a*, *b* > 0.

In the following section, we denote 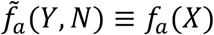 to be the function *f* in (*Y*, *N*) coordinates.

###### Proposition 5.2

If *J*_*a*_(*X*) is asymptotically scalable, then there exists two limiting functions *J*_*a*,0_(*y*): *Δ*^*n*-1^ → ℝ and *J*_*a*,∞_(*y*): *Δ*^*n*-1^ → ℝ such that the following point-wise convergence holds:

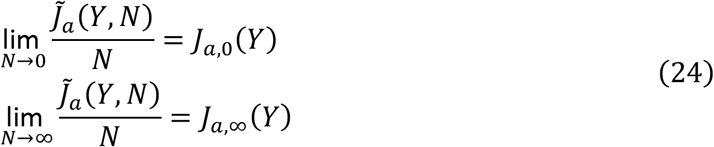

*Proof*: Consider a fixed vector *Y*\* ∈ *Δ*^*n*-1^ and define a function *h*_*Y*\*_ (*N*) ≡ *J*_*a*_(*NY*\*)/*N* on the ray {*bY*\*}, *b* > 0. We have

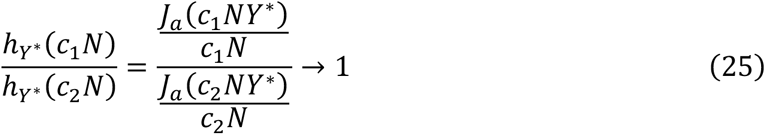

as *c*_1_, *c*_2_ → 0, by property (C3’). Hence, *h*_*Y*\*_ (*N*) approaches a constant as *N* → 0, as denoted by *J*_*a*,0_(*Y*\*). Since the above argument works for every direction *Y*\*, this defines the function *J*_*a*,0_(*Y*) on *Δ*^*n*-1^. A similar argument holds for *J*_*a*,∞_(*Y*).

Assume that the limiting functions *J*_*a*,0_(*Y*), *J*_*a*,∞_(*Y*) are continuously differentiable, which can be checked directly. Using the same stoichiometry matrix *S*, we define

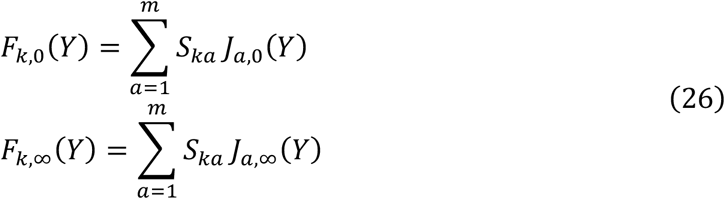

and let 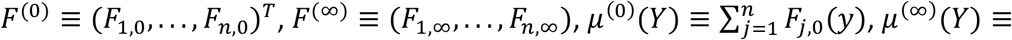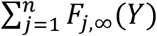. We obtain the following proposition, which is analogous to Theorem 2.3:

###### Proposition 5.3

Given an asymptotically scalable reaction network with the initial condition *X*(0) ∈ *Q*^*n*^, consider the long-term dynamics of *Y*(*t*):

i. If limsup_*t*→∞_*Y*(*t*) = liminf_*t*→∞_*Y*(*t*) = ∞, then the long-term dynamics are governed by

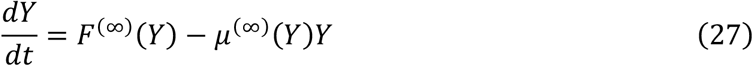
ii. If limsup_*t*→∞_*Y*(*t*) = liminf_*t*→∞_*Y*(*t*) = 0, then the long-term dynamics are governed by

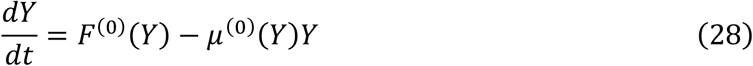
iii. If 0 < liminf_*t*→∞_*Y*(*t*) ≤ limsup_*t*→∞_(*t*) < ∞, then we have *λ* = 0.

##### Remark

The case (i) in the above proposition indicates that the system trajectory *X*(*t*) = (*Y*, *N*)(*t*) diverges to infinity along the *N*-coordinate. Hence, the long-term behavior of *Y*(*t*) is asymptotically determined by the limiting function *F*^(∞)^. If *Y*(0) is regular with respect to an ergodic probability measure *ω*, then 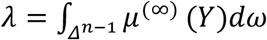. Similar arguments are also applicable for case (ii), where the trajectory (*Y*, *N*)(*t*) converges to 0 along the *N*-coordinate. For case (iii), we obtain *λ* = 0 directly since *N*(*t*) is bounded.

Finally, there are cases not listed above, which are 0 = liminf_*t*→∞_*Y*(*t*) < limsup_*t*→∞_*Y*(*t*) and liminf_*t*→∞_*Y*(*t*) < limsup_*t*→∞_*Y*(*t*) = ∞; in these cases, convergence of *λ* is inconclusive.

#### 6. Autocatalytic flux networks

One of the essential features of living systems is the existence of autocatalysis in suitable environments. Scalable reaction networks with positive growth rates can be used to address a range of questions related to autocatalytic systems. For example, what are the fundamental motifs in an autocatalytic flux network, and can we draw inferences about the long-term growth dynamics of a network by examining its topological structure?

In an SRN, one can consider the growth rates of individual nodes or fluxes as *λ*[*X*_*p*_] ≡ lim_*t*→∞_(1/*t*)log *X*_*p*_ or *λ*[*J*_*a*_] ≡ lim_*t*→∞_(1/*t*)log *J*_*a*_, when the limit exists. Given an SRN with *λ* > 0, not every node *x*_*j*_ in the network would be expected to grow as fast as the system size *N*. If a node *x*_*j*_ grows slower than the system, we have *Y*_*j*_ → 0 as *t* → ∞. In this case, we say that the node *x*_*j*_ does not contribute to the growth of the system. Our goal is to characterize the nodes that contribute to system growth and to study the mutual dependence of growth rate between fluxes and nodes.

To describe the structure of a flux network (*x*, *ϕ*), we adopt the following terminology: A node *x*_*k*_ is *upstream* of a reaction *ϕ*_*a*_, if *S*_*ka*_ < 0. A node *x*_*k*_ is *downstream* of a reaction *ϕ*_*a*_, if *S*_*ka*_ > 0. Given a reaction *ϕ*_*a*_, let *fp*(*ϕ*_*a*_), *dw*(*ϕ*_*a*_) denote the collection of its upstream and downstream nodes, respectively. A reaction *ϕ*_*a*_ is an *influx* of node *x*_*k*_, if *x*_*k*_ ∈ *dw*(*ϕ*_*a*_). A reaction *ϕ*_*a*_ is an *efflux* of node *x*_*k*_, if *x*_*k*_ ∈ *up*(*ϕ*_*a*_). Given a node *x*_*k*_, let *in*(*x*_*k*_), *out*(*x*_*k*_) denote the collection of its influxes and effluxes, respectively.

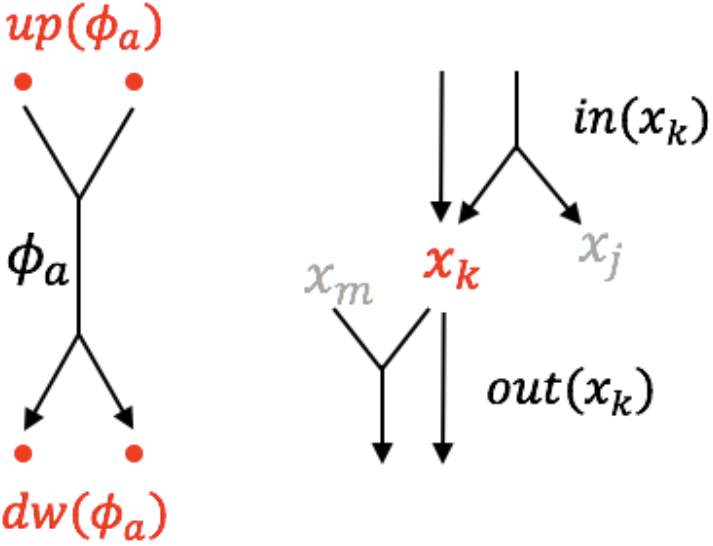

The above definition only involves the network connection; it is not related to the flux function set *J*. If we associate a flux function set to a flux network (*x*, *ϕ*, *J*), then we can introduce the following concept of “maintenance set”:

##### Definition 6.1

Let *x* = {*x*_1_, …, *x*_*n*_} denote nodes in the system. We say a node *x*_*k*_ *maintains* the reaction *ϕ*_*a*_, if *X*_*k*_ = 0 implies *J*_*a*_(*X*) = 0. The collection of nodes that maintains *ϕ*_*a*_ is called the *maintenance set* of reaction *ϕ*_*a*_, denoted by *mt*(*ϕ*_*a*_).

##### Remark

One reaction can have zero, one, or multiple maintenance nodes. For a scalable reaction flux, all of its upstream nodes are in the maintenance set (by condition C2). Still, there may be additional nodes that are not upstream of a reaction but do maintain it, such as, for example, an enzyme that is necessary for a reaction flux but is not consumed in the reaction. For SRNs, the only possible scenario for a reaction *ϕ*_*a*_ to have no maintenance set is the case where the only upstream node is the environment {*E*}. For example, consider the reaction *ϕ*_*a*_: {*E*} → *x*_3_, and the associated flux function *J*_*a*_(*X*) = *X*_1_ + *X*_2_. Biologically, *X*_1_ and *X*_2_ are redundant transporters that import *x*_3_ from the environment into the system. For this case, *mt*(*ϕ*_*a*_) is empty since *x*_1_ and *x*_2_ are redundant in the flux function. In this case, if we split the reaction *ϕ*_*a*_ into two reactions *ϕ*_*a*1_ and *ϕ*_*a*2_ and associate two flux functions *J*_*a*1_ ≡ *X*_1_, *J*_*a*2_ ≡ *X*_2_, then we have *mt*(*ϕ*_*a*1_) = *x*_1_ and *mt*(*ϕ*_*a*2_) = *x*_2_.

Practically, we can always modify the reaction set and split the flux function such that the maintenance set is nonempty under this new representation. This can be done by noticing that *J*_*a*_(*X*) = *J*_*a*_(*Y*)*N* = *J*_*a*_(*Y*)*X*_1_ + ⋯ + *J*_*a*_(*Y*)*X*_*n*_ and by setting *J*_*aj*_(*X*) = *J*_*a*_(*Y*)*X*_*j*_. It can be checked that *J*_*aj*_(*X*) are scalable functions and *mt*(*ϕ*_*aj*_) is nonempty. In general, the method to split a flux function is not unique, and we would like to choose the representation that best fits the biological intuition. In the following discussion, we assume that all reactions have a nonempty maintenance set.

###### Definition 6.2

Given a scalable reaction network (*x*, *ϕ*, *J*), a collection of fluxes *K* ≡ {*ϕ*_*a*1_, …, *ϕ*_*an*_} is called an *autocatalytic circuit*, if

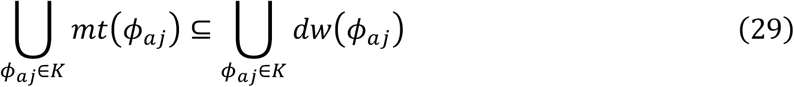

We can examine every reaction *ϕ*_*a*_ and list its maintenance set and its downstream nodes. For a flux collection *K*, we can verify whether *K* is an autocatalytic circuit by performing simple logical operations.

Note that if *K* is not an autocatalytic circuit, it is still possible to have a proper subset *K*′ ⊂ *K* such that *K*′ is autocatalytic. On the other hand, if *K* is autocatalytic, it is also possible to have a proper subset *K*′ ⊂ *K* which is autocatalytic. The following simple algorithm allows us to examine whether a flux collection *K* contains any autocatalytic circuit *K*′ ⊆ *K*. We denote 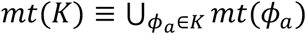 and 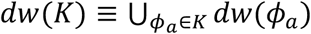.

###### Algorithm 6.3

Consider a flux network (*x*, *ϕ*, *J*) and an initial flux set *K*_1_.

1. For *j* ≥ 1, calculate *mt*(*K*_*j*_) and *dw*(*K*_*j*_). If *mt*(*K*_*j*_) ⊆ *dw*(*K*_*j*_), an autocatalytic circuit *K*_*j*_ is found and the algorithm terminates. If *mt*(*K*_*j*_) ⊈ *dw*(*K*_*j*_), examine each reaction *ϕ*_*a*_ ∈ *K*_*j*_ and test if *mt*(*ϕ*_*a*_) ∈ *dw*(*K*_*j*_). Remove every reaction *ϕ*_*a*_ such that *mt*(*ϕ*_*a*_) ∉ *dw*(*K*_*j*_), and obtain a smaller reaction set *K*_*j*+1_.
2. Repeat the above procedure until either *K*_*j*_ is empty or an autocatalytic circuit is found. If the algorithm reaches an empty set, then there is no autocatalytic circuit in *K*_1_.

*Proof*: Suppose *K* has no autocatalytic subset, the above algorithm will yield an empty set. Suppose *K* has an autocatalytic circuit *K*′, we need to show every reaction *ϕ*_*a*_ ∈ *K*′ will not be removed by this algorithm. Since *mt*(*ϕ*_*a*_) ⊆ *mt*(*K*′) ⊆ *dw*(*K*′), *ϕ*_*a*_ will not be removed at step *j* = 1. Suppose that we have *K*′ ⊆ *K*_*j*_ up to step *j*. We have *mt*(*ϕ*_*a*_) ⊆ *mt*(*K*′) ⊆ *dw*(*K*′) ⊆ *dw*(*K*_*j*_) and hence *ϕ*_*a*_ will not be removed in step *j*. Therefore, *K*′ will be a subset of *K*_*j*+1_. By induction on *j*, the reactions in the subset *K*′ will not be removed by the algorithm. Hence, after at most *m* steps (*m* is the number of reactions in the system), the algorithm reaches a stable reaction set *K*\* with *mt*(*K*\*) ⊆ *dw*(*K*\*) and *K*′ ⊆ *K*\*.

The following Lemma establishes a simple relation satisfied by scalable fluxes, with several corollaries, which will be useful in proving our main result on autocatalytic circuits below (Thm. 6.5).

###### Lemma 6.4

If a node *x*_*p*_ ∈ *mt*(*ϕ*_*a*_), then the flux function takes the form *J*_*a*_(*X*) = *R*_*a,p*_(*Y*)*X*_*p*_ where *R*_*a,p*_(*Y*) is a continuous function defined on Δ^*n*-1^.

*Proof*: We define *R*_*a,p*_(*Y*) ≡ *J*_*a*_(*Y*)/*Y*_*p*_ = *J*_*a*_(*X*)/*X*_*p*_ which is well-defined and continuous for *Y*_*p*_ > 0. Extension of *R*_*a,p*_(*Y*) to the entire simplex is possible only if *J*_*a*_(*Y*) → 0 as *Y*_*p*_ → 0, i.e. if *x*_*p*_ ∈ *mt*(*ϕ*_*a*_), and we define 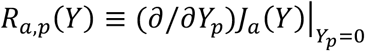 for *Y*_*p*_ where the derivative exists by scalability condition C1.

###### Corollary 6.4a

For a SRN, if *Y*(0) is regular for some ergodic measure *ω* then the node growth rates lim_*t*→∞_(1/*t*)log *X*_*p*_ exist for all nodes *x*_*p*_ of the network.

*Proof:* We can express the dynamics of *X*_*p*_ in terms of the influxes and effluxes of node *x*_*p*_ as 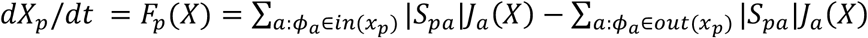. Using Lemma 6.4 for the effluxes, we have

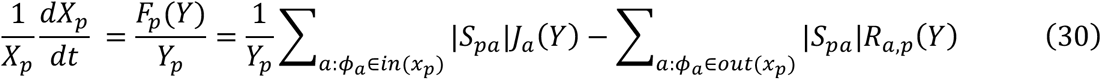

The second summation is a continuous function on the entire simplex hence integrable, while the first summation is non-negative and continuous on the simplex interior; thus the integral of *F*_*p*_(*Y*)/*Y*_*p*_ exists and takes values in ℝ ∪ +∞. We can therefore apply the ergodic theorem (which applies without the *L*^1^ assumption for functions that are bounded from below) to obtain

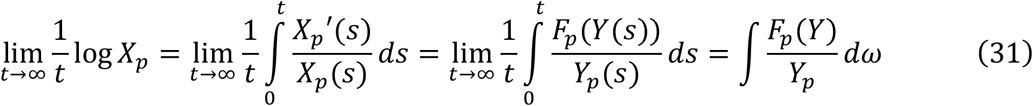

and since node growth rates are bounded by the system’s long-term growth rate *λ*, which exists and is finite by Thm. 2.3, the above limit is finite.

###### Corollary 6.4b

For an SRN with long-term growth rate *λ* > 0, if a flux *J*_*a*_ has growth rate *λ*, then all of its maintenance nodes *x*_*p*_ grow at rate *λ*.

*Proof:* Let *x*_*p*_ be a maintenance node of *J*_*a*_. By Lemma 6.4, *J*_*a*_(*X*) = *R*_*a,p*_(*Y*)*X*_*p*_. Since *R*_*a,p*_(*Y*) is a bounded non-negative function, positive growth of *J*_*a*_ implies that *X*_*p*_ must grow at least as fast as *J*_*a*_. Since *J*_*a*_ is growing at the same rate as the whole system and *X*_*p*_ is a component of the system, *X*_*p*_ must be growing exactly at the rate *λ*.

###### Theorem 6.5

Consider an SRN (*x*, *ϕ*, *J*) with an initial condition *Y*(0), which is regular for an ergodic measure *ω*. If the long-term growth rate *λ* is positive, then there exists at least one autocatalytic circuit *K* in the network such that every node *x*_*j*_ ∈ *mt*(*K*) has a long-term growth rate *λ*.

*Proof:* By Cor. 6.4a, all node growth rates *λ*[*X*_*p*_] exist. We provide a recursive procedure to find an autocatalytic circuit *K* satisfying Eq. (29). Given a network with growth rate *λ* > 0, there is at least one node *x*_*p*_ with *λ*[*X*_*p*_] = *λ*, and it is clear that there exists at least one reaction *ϕ*_(01)_ ∈ *in*(*x*_*p*_) with *λ*[*J*_*a*_] = *λ*. We denote *K*_0_ ≡ {*ϕ*_(01)_}.

Below we provide a procedure that starts with a collection of reactions *K*_*j*_ all of which have growth rate *λ*, and enlarges the collection to *K*_*j*+1_. Let *K*_*j*_ = {*ϕ*_(*j*1)_, …, *ϕ*_(*jn*)_}. For each flux *ϕ*_(*ja*)_, we examine its maintenance set *mt*(*ϕ*_*ja*_(*X*)). If *mt*(*ϕ*_*ja*_) is nonempty, all nodes *x*_*p*_ ∈ *mt*(*ϕ*_*ja*_) have the same growth rate *λ* based on Cor. 6.4b, thus every node *x*_*p*_ has at least one influx with growth rate *λ*. (If there are multiple influxes with growth rate *λ*, we are free to choose any of them.) We denote *ϕ*[*x*_*p*_] to be the influx chosen from *in*(*x*_*p*_), as shown here:

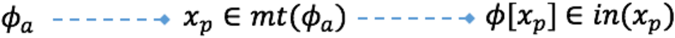

Now, we define

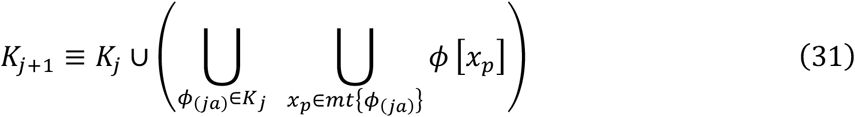

noting that all fluxes in *K*_*j*+1_ have growth rate *λ*. By this procedure, the size {*K*_*j*_} monotonically increases. Since the total number of reactions in the network is finite, there exists a maximal collection *K* such that *K* is invariant under this procedure. Next, we must show that *mt*(*K*) ⊆ *dw*(*K*), i.e., *K* is an autocatalytic circuit.

Given an arbitrary reaction *ϕ*_*a*_ ∈ *K*, suppose that this reaction is recruited into *K* at step *j* for the first time. For each node *x*_*k*_ ∈ *mt*(*ϕ*_*a*_), there exists at least one reaction that will be recruited into *K* at step *j* + 1, denoted as *ϕ*[*x*_*k*_] (if this reaction does not yet belong to *K*_*j*_). Hence, *x*_*k*_ ∈ *dw*(*ϕ*[*x*_*k*_]) ⊂ *dw*(*K*). This shows *mt*(*ϕ*_*a*_) ∈ *dw*(*K*) for an arbitrary reaction *ϕ*_*a*_ ∈ *K*, and therefore, *K* is an autocatalytic circuit.

By construction, every flux *J*_*a*_ associated with *ϕ*_*a*_ ∈ *K* has a growth rate *λ* > 0. By Cor. 6.4b, all maintenance nodes of *K* has a growth rate *λ*.

##### Remark 1

The maximal collection *K* constructed above is not necessarily unique. There may be several redundant autocatalytic circuits in the flux network. Each of them can be constructed by choosing a different initial reaction *ϕ*_(01)_ or a different *ϕ*[*x*_*k*_] during the recursion steps.

##### Remark 2

The condition *λ* > 0 is essential in Theorem 6.5. If the system has zero or negative long-term growth rate, the statements in Propositions 6.4a and 6.4b no longer hold. That is, the growth rate of each node in an autocatalytic circuit does not necessarily need to be the same.

This method is remarkable in the sense that we do not need to solve the differential equation (either analytically or numerically). The procedure only requires that we inspect each flux function and obtain its maintenance set. The remaining computation is purely algebraic. It is the flux network connection and the property of positive growth rate that constrain the growth rate between nodes and fluxes.

###### Example 6.6: Random networks

To characterize random networks, we define *P*_*auto*_ as the probability that a random reaction network is autocatalytic, which depends on its statistical properties. Intuitively, for Eq. 6 to be satisfied, one should decrease the average number of *mt*(*ϕ*_*a*_) and increase the average number of *dw*(*ϕ*_*a*_). Furthermore, given a random reaction *ϕ*_*a*_, the chance of a given node to belong to *mt*(*ϕ*_*a*_) or *dw*(*ϕ*_*a*_) could be different. Intuitively, if some nodes acted as “hubs” to maintain many other nodes, the probability *P*_*auto*_ should be higher (Fig. S6A). In contrast, if a node required many other upstream nodes (Fig. S6B), *P*_*auto*_ should be lower.

To test how network topology affects *P*_*auto*_, we constructed an ensemble of random reaction networks with a fixed number of nodes (*n* = 100), but with different connections of reactions. Consistent with our expectation, we found that *P*_*auto*_ transitions from 0 to 1 as the number of reactions increases in the network (Fig. S6). To see how hub-like connections affect *P*_*auto*_, we randomly sampled the maintenance and downstream nodes of every reaction from power-law distributions with exponents γ_*mt*_ and γ_*dw*_. We found that *P*_*auto*_ is strongly affected by both exponents. For example, network connections with uniform probability (γ_*mt*_ = γ_*dw*_ = 0) usually required 150 or more reactions to be autocatalytic (Fig. S6A). However, for a hub-like maintenance set with skewed probability γ_*mt*_ = 2, the number of reactions required could be reduced to 100. This contrasts with a hub-like downstream set with skewed probability γ_*dw*_ = 1, for which more than 400 reactions were required for the network to be autocatalytic.

## 7. Appendix Existence and uniqueness of solutions for Eq. (19)

In this section, we denote |· | as the *l*^1^ norm in Euclidean space. We verify the following conditions in Ref(47),

(A1) |*F*^(*Ito*)^(*X*)| + |*H*(*X*)| ≤ *C*_1_(1 + |*X*|)

(A2) |*F*^(*Ito*)^(*X*) − *F*^(*Ito*)^(*X*′)| + |*H*(*X*) − *H*(*X*′)| ≤ *C*_2_|*X* − *X*′|for all *X*, *X*′ ∈ *Q*^*n*^ and positive constants *C*_1_, *C*_2_.

The conversion between Ito and Stratonovich integrals is described by Eq. (20). To show (A1), we obtain |*F*_*k*_(*X*)| = |*X*||*F*_*k*_(*Y*)| because of the scalable condition (C3). According to the continuous differentiability condition (C1) and the compactness of *Δ*^*n*-1^, there exists a constant *B*_*F*_ > 0 such that |*F*_*k*_(*Y*)| ≤ *B*_*F*_ for all *Y* ∈ *Δ*^*n*-1^ and for every *k*. The function *H*_*k*_(*X*) is bounded similarly by a constant *B*_*H*_.

Next, by the scalable property (N3), the value of 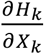 is scale-invariant (i.e., 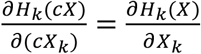 for all *c* > 0). Hence, this value is specified by 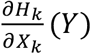 on the unit simplex *Δ*^*n*-1^. Since *H* is continuously differentiable on the simplex, this term is again bounded by a constant. Hence, the drift correction term 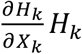 is bounded by *B*_*drift*_|*X*| for all *X* and all *k*, with constant *B*_*drift*_ > 0. Together, we can choose *C*_1_ = *n*(*B*_*F*_ + *B*_*drift*_ + *B*_*H*_) and condition (A1) is satisfied.

To show (A2), we use the mean value theorem to get *F*_*k*_(*X*) − *F*_*k*_(*X*′) = *∇F*_*k*_(*X*\*)(*X* − *X*′) with some *X*\* at the interior of segment 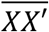. Note that the gradient of the scalable function is scale-invariant. Hence, because of the continuity condition (C1), there is a positive constant *B*_*∇F*_ as upper bound of 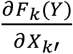 for all *Y* ∈ *Δ*^*n*-1^ and for all *k*, *k*′. Hence, we have

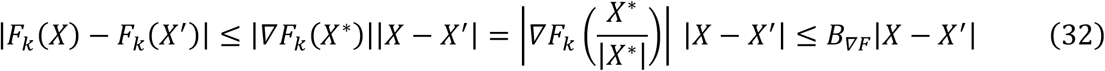

A similar argument holds for *H* For the drift correction term, note that

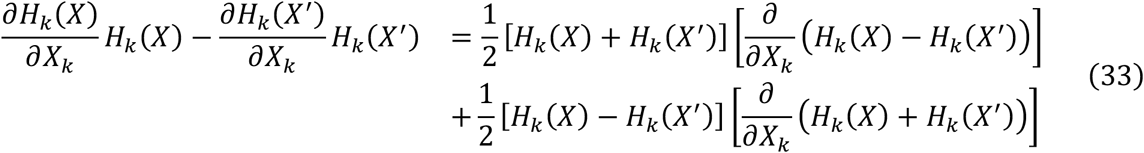

Using similar arguments to those mentioned above, the difference of the drift-correction term can be bounded by *B*_*drift*′_|*X* − *X*′| with a constant *B*_*drift*′_ > 0. Together, we can choose *C*_2_ = *n*(*B*_*∇F*_ + *B*_*drift*′_ + *B*_*∇H*_) and condition (A2) is satisfied.

### Proof of Proposition 3.2

We will use the following two theorems. We denote *Q*_*k*_ ≡ {*X*: *X*_*k*_ = 0} be a plane in ℝ^*n*^ and 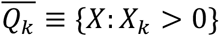.

#### Theorem 3.2.1

[Ref(48), Thm 1.1]: For an SDE with the diffusion operator 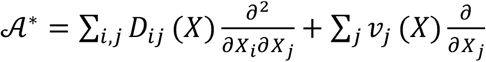, if there exists a real-valued function *V*(*X*) that satisfies the following conditions

1. there exists a constant *C* > 0 such that 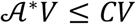 for all 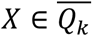
2. *V*(*X*) → ∞ whenever *dist*(*X*, *Q*_*k*_) → 0,

then a trajectory starting with *X*_*k*_ > 0 cannot reach *Q*_*k*_ at a finite time.

The next theorem states that for an SDE satisfying the above criteria, the function *V*(*X*) in the above theorem can be found explicitly.

#### Theorem 3.2.2

[Ref(48). Thm 1.1, Ref(49), Thm 9.4.1]: Consider the SDE in Theorem 3.2.1, with diffusion operator *D*_*ij*_(*X*) and deterministic flow *v*_*k*_(*X*). The function *V*(*X*) can be chosen as 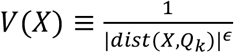 for a fixed *ϵ* > 0, if the following criteria are satisfied:

(A1) 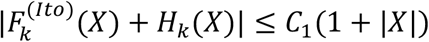 for a constant *C*_1_ > 0 and for all *X*.
(A2) 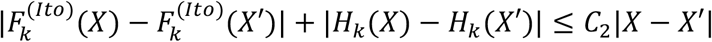 for a constant *C*_2_ > 0 and for all *X*, *X*′
(A3) If *X* ∈ *Q*_*k*_, then *D*_*kk*_(*X*) = 0.
(A4) If *X* ∈ *Q*_*k*_ then 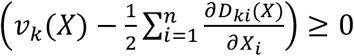

The quantity within the parenthesis in (A4) is called the *Fichera drift*.

To proceed, we show that the conditions (A1-A4) hold for the scalable SDE (19). Conditions (A1) and (A2) were demonstrated to hold above. To verify (A3) and (A4), we calculate the diffusion matrix and drift vector by standard SDE rules ([Ref(47)]):

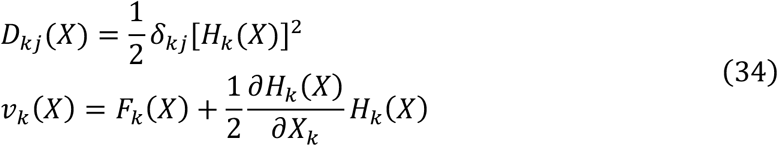

Since the matrix *D* is diagonalized, we have *D*_*kk*_(*X*) = [*H*_*k*_(*X*)]^2^ and condition (N2) implies condition (A3). For condition (A4), we first notice that the upstream-limited condition of flux (C2) implies that *F*_*k*_(*X*) ≥ 0 whenever *X*_*k*_ = 0. Second, *H*_*k*_(*X*) = 0 on *Q*_*k*_, while 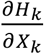 is bounded by a constant. Together, we have *v*_*k*_(*X*) ≥ 0 on *Q*_*k*_. Finally, we obtain

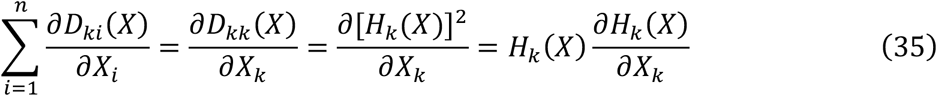

As above, this term is zero on *Q*_*k*_. This shows that the Fichera drift on *Q*_*k*_ satisfies the condition in (A4).

We have shown that for the scalable SDE, conditions (A1) to (A4) are satisfied. By Theorems 3.2.1 and 3.2.2, for any trajectory *X*(*t*) with *X*_*k*_(0) > 0, the system does not reach the plane *Q*_*k*_ at a finite time. This implies that *N* = *X*_0_+… +*X*_2_ does not reach the origin at a finite time. Together, the trajectory *Y*(*t*) = *X*(*t*)/*N*(*t*) with *Y*_*k*_(0) > 0 does not reach a simplex boundary at a finite time. Along the proof, we have shown that *v*_*k*_(*X*) ≥ 0 and the component *X*_*k*_ becomes deterministic on *Q*_*k*_. Hence the trajectory *Y*(*t*) is confined in the unit simplex.

## Methods: Simulation and analysis of scalable reaction networks

### 1. Numerical solution of ordinary differential equations

All ordinary differential equations (ODE) were integrated by MATLAB function *ode45* with relative tolerance *RelTol* = 10^−3^ and absolute tolerance *RelTol* = 10^−6^. In Fig. 2b, c, and g, the trajectory *Y*(*t*) was also calculated by Mathematica function *NDSolve* with default accuracy, yielding consistent results. In Fig. S2, the trajectory *Y*(*t*) was integrated by a log-transform ODE for improving the accuracy when *Y*_*k*_ is close to zero. For calculating the long-term growth rate *λ*, we simulated the trajectory *Y*(*t*) for a sufficiently long time such that *Y*(*t*) converges to its attractor. After convergence, we averaged the instantaneous growth rate *μ*(*Y*(*t*_*j*_)) with time step = 0.01 (time unit) to obtain growth rate *λ*. To simulate a system with noise, the differential equation was integrated with an additional Gaussian noise term for every integration time step.

### 2. Calculation of Poincare section and bifurcation analysis

The Poincare section and bifurcation analysis in Fig. S3A were calculated using the following procedure:

1. A Poincare section plane {*Y*^*P*^ = *c*^*P*^} and a projection coordinate *Y*_*a*_ are chosen.
2. The ODE system is integrated from time 0 to a large time *T*_*Pre*_ = 2000 to allow the trajectory to converge to the attractor. The final position *Y*^(*eq*)^ of the trajectory is used as the initial condition in the next step.
3. The ODE system is integrated from time 0 to a large time *T*^*L*^ = 1000, with initial condition *Y*(0) = *Y*^(*eq*)^ and a given parameter *κ*. This generates a trajectory {*Y*(*t*_*j*_)} with time step size Δ*t* = 0.02. When a time point *t*_*j*_ satisfied

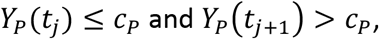

the coordinate *Y*_*a*_(*t*_*j*_) is recorded. This procedure gave a list of scalars {*Y*_*a*_(*t*_*j*1_),…, *Y*_*a*_(*t*_*jC*_)}.
4. The same procedure as in 3 is repeated with a different parameter *α*. For each *α*, a list of points 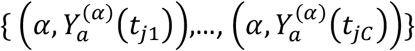 is obtained on the *α* − *Y*_*a*_ plane. The two-dimensional scatter plot shown in Fig. S3B was generated by using all *α* values and all points in their lists.

### 3. Calculation of the largest Lyapunov exponent

The largest Lyapunov exponent (LLE) in Fig. S3B was calculated by the following procedure:

1. The ODE system is integrated for a long-time interval from time 0 to 2000 to allow the trajectory to converge to the attractor. The final position *Y*^(*eq*)^ of trajectory is used as the initial condition in the next step.
2. We set *Y*_*start*_ ← *Y*^(*eq*)^

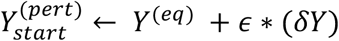

where (*δY*) is an isotropic random vector with |*δY*| ≤ 0.1 and ∑_*j*_(*δY*)_*j*_ = 0.
3. The system with the initial condition *Y*_*start*_ is integrated for a short time interval [0, *τ*] with *τ*=20 to obtain *Y*(*τ*). Similarly, the system with the initial condition 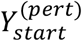 is integrated for a short time interval [0, 20] to obtain *Y*^(*pert*)^(*τ*). Using these results, the exponent of trajectory divergence *E*_1_ is calculated as follows:

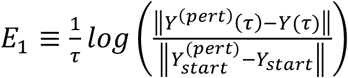
4. The parameter is reset as *Y*_*start*_ ← *Y*^(*τ*)^

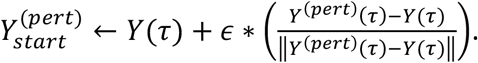

with *ϵ* = 0.1 and the procedure described by step 3 is repeated to obtain another pair of trajectories *Y*(*τ*) and *Y*^(*pert*)^(*τ*). The trajectory divergence *E*_2_ is calculated similarly. This aforementioned procedure is repeated 50 times to obtain a list of values {*E*_1_, …, *E*_50_}. The average divergence *E* is calculated by the algebraic averaging of *E*_*j*_.
5. The procedure in steps 2 to 4 is repeated five times, where each repeat has different random perturbations and yields a different exponential divergence *E*^(*k*)^. The LLE shown in Fig. S3B is calculated by averaging *E*^(*k*)^ from all simulations.

Note that for choosing the proper values for constants *s* and *τ*, they should satisfy

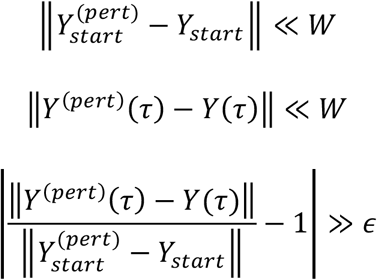

where *W* is the characteristic length scale of the attractor and *ϵ* is the numerical accuracy of the computation. (For the autocatalytic double repressilator, we chose *τ* = 20 and *s* = 0.1.)

### 4. Mathematical models of scalable reaction networks

Parameters shown in blue are the ones we varied in this study.

#### (i) Autocatalytic single-repressilator model

The autocatalytic single-repressilator model has four nodes *X*_1_ to *X*_4_ (see Fig 3A and the main text), with the following reactions:

| Reaction | Flux function |
| --- | --- |
| $\phi_1: \{\text{env}\} \rightarrow x_1$ | $J_1 = b_2X_2 + b_3X_3 + b_4X_4$ |
| $\phi_2: x_1 \rightarrow x_2$ | $J_2 = \frac{c_{syn}X_1}{1 + KY_3^\theta}$ |
| $\phi_3: x_1 \rightarrow x_3$ | $J_3 = \frac{c_{syn}X_1}{1 + KY_4^\theta}$ |
| $\phi_4: x_1 \rightarrow x_4$ | $J_4 = \frac{c_{syn}X_1}{1 + KY_2^\theta}$ |
| $\phi_5: x_2 \rightarrow \{\emptyset\}$ | $J_5 = d_2X_2$ |
| $\phi_6: x_3 \rightarrow \{\emptyset\}$ | $J_6 = d_3X_3$ |
| $\phi_7: x_4 \rightarrow \{\emptyset\}$ | $J_7 = d_4X_4$ |

Parameters used in simulations were as follows:

|  |  |
| --- | --- |
| $b_2 = 0.5$ (1/time) | $d_2 = 0.25$ (1/time) |
| $b_3 = 0.4$ (1/time) | $d_3 = 0.3$ (1/time) |
| $b_4 = 0.3$ (1/time) | $d_4 = 0.2$ (1/time) |
| $c_{syn} = 20$ (1/time) | $K = 10 - 10000$ |
| | $\theta = 1 - 4$ |

#### (ii) Autocatalytic double-repressilator model

The autocatalytic double-repressilator model has seven nodes *X*_1_ to *X*_7_ (see Fig. 3F and the main text), with the following reactions:

| Reaction | Flux function |
| --- | --- |
| $\phi_1: \{\text{env}\} \rightarrow x_1$ | $J_1 = b_2X_2 + b_3X_3 + b_4X_4 + b_5X_5 + b_6X_6 + b_7X_7$ |
| $\phi_2: x_1 \rightarrow x_2$ | $J_2 = \frac{cX_1}{1 + K_a Y_3^\theta} \frac{1}{1 + MY_5^\phi}$ |
| $\phi_3: x_1 \rightarrow x_3$ | $J_3 = \frac{cX_1}{1 + K_a Y_4^\theta}$ |
| $\phi_4: x_1 \rightarrow x_4$ | $J_4 = \frac{cX_1}{1 + K_a Y_2^\theta}$ |
| $\phi_5: x_1 \rightarrow x_5$ | $J_5 = \frac{cX_1}{1 + K_b Y_6^\theta} \frac{1}{1 + MY_2^\phi}$ |
| $\phi_6: x_1 \rightarrow x_6$ | $J_6 = \frac{cX_1}{1 + K_b Y_7^\theta}$ |
| $\phi_7: x_1 \rightarrow x_7$ | $J_7 = \frac{cX_1}{1 + K_b Y_5^\theta}$ |
| $\phi_8: x_2 \rightarrow \{\emptyset\}$ | $J_8 = d_2 X_2$ |
| $\phi_9: x_3 \rightarrow \{\emptyset\}$ | $J_9 = d_3 X_3$ |
| $\phi_{10}: x_4 \rightarrow \{\emptyset\}$ | $J_{10} = d_4 X_4$ |
| $\phi_{11}: x_5 \rightarrow \{\emptyset\}$ | $J_{11} = d_5 X_5$ |
| $\phi_{12}: x_6 \rightarrow \{\emptyset\}$ | $J_{12} = d_6 X_6$ |
| $\phi_{13}: x_7 \rightarrow \{\emptyset\}$ | $J_{13} = d_7 X_7$ |

Parameters used in simulations are as follows:

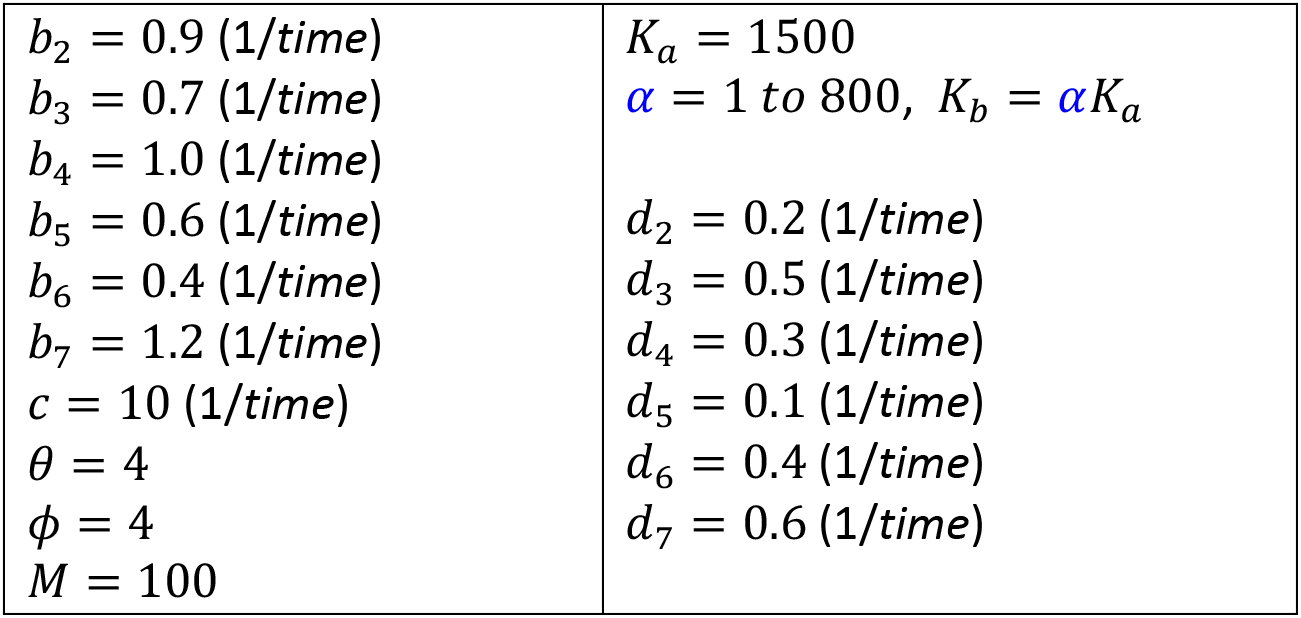

#### (iii) Ecosystem model of three competing and cross-feeding species

##### Cross-feeding model for a turbidostat-like system (unlimited resource)

The three-species model has six nodes: *x*_1_ to *x*_3_ represent three species and *m*_1_ to *m*_3_ represent three metabolites, each secreted by one species (see Fig. 4A in the main text). *X*_1_, *X*_2_, *X*_3_, *M*_1_, *M*_2_, *M*_3_ denote the total amount of biomass of species and metabolites and *N* is defined by *X*_1_ + *X*_2_ + *X*_3_ + *M*_1_ + *M*_2_ + *M*_3_. The variables 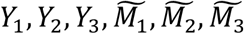 represent the relative amount of biomass of species *x*_1_ to *x*_3_ (normalized as *Y*_*j*_ = *X*_*j*_/*N* and 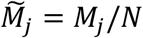).

For each species, the influxes from the environment (*ϕ*_1_ to *ϕ*_3_) are modeled by a Lotka-Volterra-like quadratic function. We modified the function to be scalable. The production of secreted metabolites are linear fluxes (*ϕ*_4_ to *ϕ*_6_) that are proportional to the species abundance. There is a total of six cross-feeding fluxes, with each metabolite being able to feed one or both species that do not produce this metabolite. The cross-feeding fluxes are Michaelis-Menten type (*ϕ*_12_, *ϕ*_13_, *ϕ*_21_, *ϕ*_23_, *ϕ*_31_, *ϕ*_32_).

| Reaction | Flux function |
| --- | --- |
| $\phi_1: \{\text{env}\} \rightarrow x_1$ | $J_1 = r_1 X_1 \left( 1 - \sum_{j=1}^3 a_{1j} Y_j \right)$ |
| $\phi_2: \{\text{env}\} \rightarrow x_2$ | $J_2 = r_2 X_2 \left( 1 - \sum_{j=1}^3 a_{2j} Y_j \right)$ |
| $\phi_3: \{\text{env}\} \rightarrow x_3$ | $J_3 = r_3 X_3 \left( 1 - \sum_{j=1}^3 a_{3j} Y_j \right)$ |
| $\phi_4: x_1 \rightarrow m_1$ | $J_4 = b_1 X_1$ |
| $\phi_5: x_2 \rightarrow m_2$ | $J_5 = b_2 X_2$ |
| $\phi_6: x_3 \rightarrow m_3$ | $J_6 = b_3 X_3$ |
| $\phi_{12}: m_2 \rightarrow x_1$ | $J_{12} = c_{12} X_1 \frac{M_2}{KN + M_2}$ |
| $\phi_{13}: m_3 \rightarrow x_1$ | $J_{13} = c_{13} X_1 \frac{M_3}{KN + M_3}$ |
| $\phi_{21}: m_1 \rightarrow x_2$ | $J_{21} = c_{21} X_2 \frac{M_1}{KN + M_1}$ |
| $\phi_{23}: m_3 \rightarrow x_2$ | $J_{23} = c_{23} X_2 \frac{M_3}{KN + M_3}$ |
| $\phi_{31}: m_1 \rightarrow x_3$ | $J_{31} = c_{31} X_3 \frac{M_1}{KN + M_1}$ |
| $\phi_{32}: m_2 \rightarrow x_3$ | $J_{32} = c_{32} X_3 \frac{M_2}{KN + M_2}$ |

The system dynamics is governed by the following differential equations:

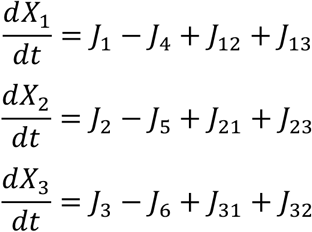

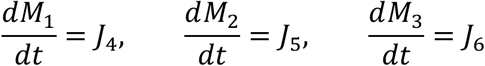

Parameters used in simulations are as follows:

|  |
| --- |
| $r_1, r_2, r_3 = 1.0 \text{ (1/time)}$<br>$b_1, b_2, b_3 = 0.2 \text{ (1/time)}$<br>$K = 0.2$<br>$a_{ij} = 0 - 0.8 \text{ (scalar)}$<br>$c_{ij} = 0 - 0.5 \text{ (1/time)}$ |

For all simulations, the coefficients *a*_*ij*_ were sampled from a uniform distribution between [0, 0.8].

For weak and strong cross-feeding, the coefficients *c*_*ij*_ were sampled from uniform distributions between [0, 0.1] and [0, 0.5], respectively. For the no cross-feeding cases, the coefficients *c*_*ij*_ were set to 0.

Each condition (no cross-feeding, weak cross-feeding and strong cross-feeding) were simulated with a 50,000 random parameter set. For each parameter set, five random initial conditions (uniformly sampled from a simplex space) were simulated. A customized script was developed to classify if the normalized system dynamics (*Y*(*t*)) reaches a fixed point. Systems with nontrivial dynamics in the long term were subjected to manual inspection and classified as fixed points, limit cycles or heteroclinic cycles. For most of random parameter sets, the model converged to a fixed point. About 0.5% of the cases converged to a heteroclinic cycle, while the limit cycle cases were even rarer (2 cases from the 150,000 random parameter sets).

##### System parameters for data shown in Fig. 4C

Initial condition: (*X*_1_, *X*_2_, *X*_3_, *M*_1_, *M*_2_, *M*_3_) = (0.25, 0.25, 0.4, 0.02, 0.03, 0.05)

|  |  |
| --- | --- |
| $a_{ij} = \begin{pmatrix} 0.536 & 0.486 & 0.214 \\ 0.266 & 0.055 & 0.763 \\ 0.774 & 0.654 & 0.182 \end{pmatrix}$ | $c_{ij} = \begin{pmatrix} 0 & 0.044 & 0.098 \\ 0.251 & 0 & 0.212 \\ 0.436 & 0.426 & 0 \end{pmatrix}$ |

##### System parameter set for data shown in Fig. 4D

Initial condition: (*X*_1_, *X*_2_, *X*_3_, *M*_1_, *M*_2_, *M*_3_) = (0.1, 0.1, 0.7, 0.05, 0.03, 0.02)

|  |  |
| --- | --- |
| $a_{ij} = \begin{pmatrix} 0.553 & 0.694 & 0.061 \\ 0.408 & 0.348 & 0.531 \\ 0.771 & 0.396 & 0.069 \end{pmatrix}$ | $c_{ij} = \begin{pmatrix} 0 & 0.098 & 0.108 \\ 0.378 & 0 & 0.165 \\ 0.030 & 0.108 & 0 \end{pmatrix}$ |

##### Cross-feeding model for a chemostat-like system (resource in limited amount)

The simulations in Fig. 4B-D were performed using usual SRNs in which there are unlimited upstream resources. For a chemostat-like system in which an essential nutrient is limited, we made several modifications to the turbidostat-like equation in the above section.

We first describe the chemostat equations for the cross-feeding model. Below, we denote the variable [*S*], [*X*_*j*_],[*M*_*j*_] as the *concentrations* (mass per volume) of the external metabolite, the cells, and cross-feeding metabolites. The flux functions are:

| Reaction | Flux function |
| --- | --- |
| $\phi_1: \{\text{env}\} \rightarrow x_1$ | $J_{1c} = \frac{r_{\max}[S]}{K_S + [S]} [X_1] \left(1 - \sum_{j=1}^3 a_{1j}[X_j]\right)$ |
| $\phi_2: \{\text{env}\} \rightarrow x_2$ | $J_{2c} = \frac{r_{\max}[S]}{K_S + [S]} [X_2] \left(1 - \sum_{j=1}^3 a_{2j}[X_j]\right)$ |
| $\phi_3: \{\text{env}\} \rightarrow x_3$ | $J_{3c} = \frac{r_{\max}[S]}{K_S + [S]} [X_3] \left(1 - \sum_{j=1}^3 a_{3j}[X_j]\right)$ |
| $\phi_4: x_1 \rightarrow m_1$ | $J_4 = b_1[X_1]$ |
| $\phi_5: x_2 \rightarrow m_2$ | $J_5 = b_2[X_2]$ |
| $\phi_6: x_3 \rightarrow m_3$ | $J_6 = b_3[X_3]$ |
| $\phi_{12}: m_2 \rightarrow x_1$ | $J_{12} = c_{12}[X_1] \frac{[M_2]}{K + [M_2]}$ |
| $\phi_{13}: m_3 \rightarrow x_1$ | $J_{13} = c_{13}[X_1] \frac{[M_3]}{K + [M_3]}$ |
| $\phi_{21}: m_1 \rightarrow x_2$ | $J_{21} = c_{21}[X_2] \frac{[M_1]}{K + [M_1]}$ |
| $\phi_{23}: m_3 \rightarrow x_2$ | $J_{23} = c_{23}[X_2] \frac{[M_3]}{K + [M_3]}$ |
| $\phi_{31}: m_1 \rightarrow x_3$ | $J_{31} = c_{31}[X_3] \frac{[M_1]}{K + [M_1]}$ |
| $\phi_{32}: m_2 \rightarrow x_3$ | $J_{32} = c_{32}[X_3] \frac{[M_2]}{K + [M_2]}$ |

The modified system ODE is:

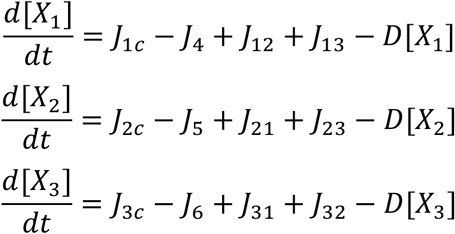

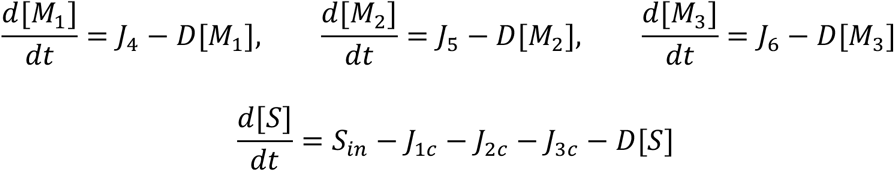

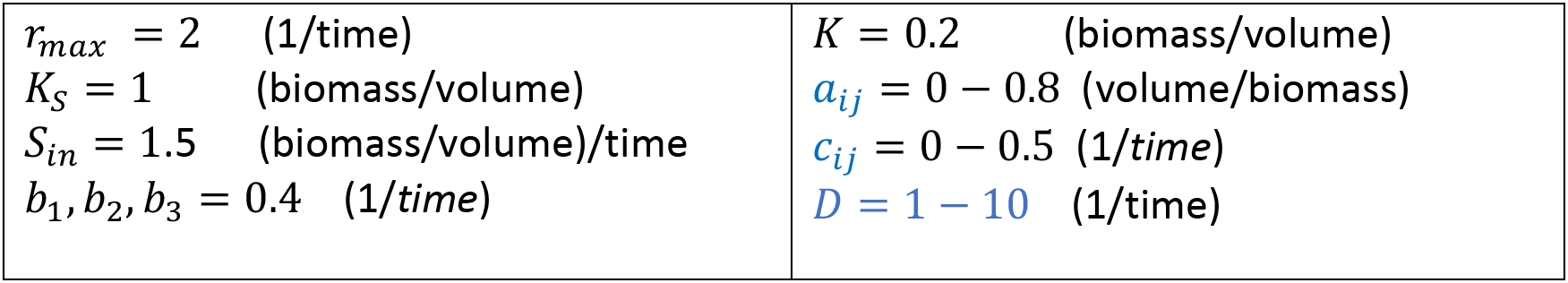

|  |  |
| --- | --- |
| $r_{max} = 2$ (1/time) | $K = 0.2$ (biomass/volume) |
| $K_S = 1$ (biomass/volume) | $a_{ij} = 0 - 0.8$ (volume/biomass) |
| $S_{in} = 1.5$ (biomass/volume)/time | $c_{ij} = 0 - 0.5$ (1/time) |
| $b_1, b_2, b_3 = 0.4$ (1/time) | $D = 1 - 10$ (1/time) |

Next, we convert the above chemostat equations into SRN-equivalent equations. We include the “solvent node” *w* with the solvent concentration [*W*] and calculate the total volume of the chemostat system:

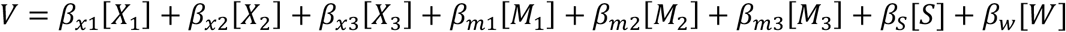

where 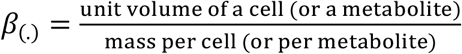

Since the chemostat has a constant volume, we have *V* as a time-independent constant. Now, we can define (*X*_1_, *X*_2_, *X*_3_, *M*_1_, *M*_2_, *M*_3_, *S*, *W*) as the total mass of cells or molecules in the chemostat. This gives the relation *X*_*j*_ = *V*[*X*_*j*_], *M*_*j*_ = *V*[*M*_*j*_] and *S* = *V*[*S*]. The SRN-type flux functions are calculated using the following equations:

| Reaction | Flux function |
| --- | --- |
| $\phi_1: \{\text{env}\} \rightarrow x_1$ | $J_{1c} = \frac{r_{max}(S/V)}{K_S + (S/V)} X_1 \left( 1 - \sum_{j=1}^3 a_{1j}(X_j/V) \right)$ |
| $\phi_2: \{\text{env}\} \rightarrow x_2$ | $J_{2c} = \frac{r_{max}(S/V)}{K_S + (S/V)} X_2 \left( 1 - \sum_{j=1}^3 a_{2j}(X_j/V) \right)$ |
| $\phi_3: \{\text{env}\} \rightarrow x_3$ | $J_{3c} = \frac{r_{max}(S/V)}{K_S + (S/V)} X_3 \left( 1 - \sum_{j=1}^3 a_{3j}(X_j/V) \right)$ |
| $\phi_4: x_1 \rightarrow m_1$ | $J_4 = b_1 X_1$ |
| $\phi_5: x_2 \rightarrow m_2$ | $J_5 = b_2 X_2$ |
| $\phi_6: x_3 \rightarrow m_3$ | $J_6 = b_3 X_3$ |
| $\phi_{12}: m_2 \rightarrow x_1$ | $J_{12} = c_{12} X_1 \frac{(M_2/V)}{K + (M_2/V)}$ |
| $\phi_{13}: m_3 \rightarrow x_1$ | $J_{13} = c_{13} X_1 \frac{(M_3/V)}{K + (M_3/V)}$ |
| $\phi_{21}: m_1 \rightarrow x_2$ | $J_{21} = c_{21} X_2 \frac{(M_1/V)}{K + (M_1/V)}$ |
| $\phi_{23}: m_3 \rightarrow x_2$ | $J_{23} = c_{23}X_2 \frac{(M_3/V)}{K + (M_3/V)}$ |
| $\phi_{31}: m_1 \rightarrow x_3$ | $J_{31} = c_{31}X_3 \frac{(M_1/V)}{K + (M_1/V)}$ |
| $\phi_{32}: m_2 \rightarrow x_3$ | $J_{32} = c_{32}X_3 \frac{(M_2/V)}{K + (M_2/V)}$ |

The equivalent SRN follows the ODE:

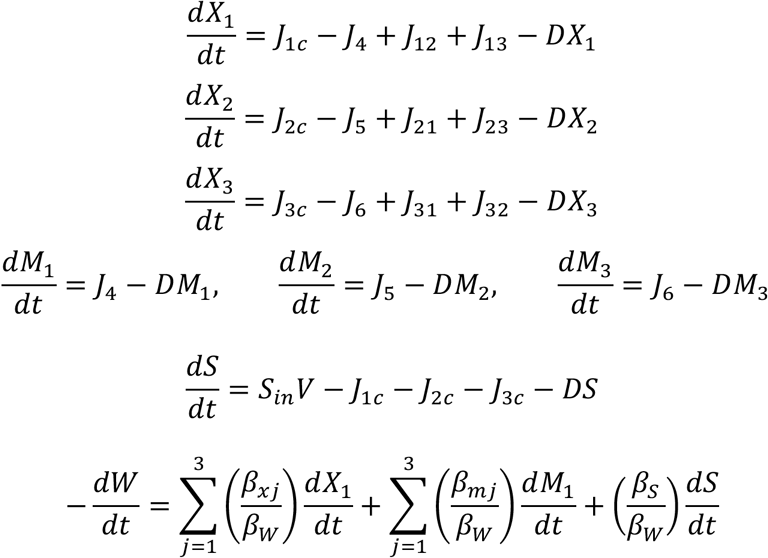

By choosing the proper volume unit, we can set *V* = 1 in the numerical simulation. In this case, the flux functions of the SRN-equivalent model become identical to those of chemostat model. In our simulation, we choose *V* = 1 liter and measure the total mass of *X*_*j*_, *M*_*j*_, *S*, *W* in grams. The parameters of the SRN-equivalent model are:

|  |  |
| --- | --- |
| $r_{max} = 2$ (1/time) | $K = 0.2$ (g/L) |
| $K_S = 1$ (g/L) | $a_{ij} = 0 - 0.8$ (L/g) |
| $S_{in} = 1.5$ (g/L)/time | $c_{ij} = 0 - 0.5$ (1/time) |
| $b_1, b_2, b_3 = 0.4$ (1/time) | $D = 1 - 10$ (1/time) |

We performed simulations using the above equations and the random coefficients *a*_*ij*_ and *c*_*ij*_, where *a*_*ij*_ was sampled from a uniform distribution on [0,0.8] and *c*_*ij*_ was sampled from a uniform distribution on [0,0.1]. For most random parameters, the model converged to a fixed point. In about 0.2% of the cases the model converged to a heteroclinic cycle, while the limit cycle cases were again rare (3 out of 120,000 random parameter sets).

##### System parameters of Fig. 4E

Initial condition: (*X*_1_, *X*_2_, *X*_3_, *M*_1_, *M*_2_, *M*_3_, *S*) = (0.018, 0.012, 0.038, 0.592, 0.296, 0.044, 0.500)

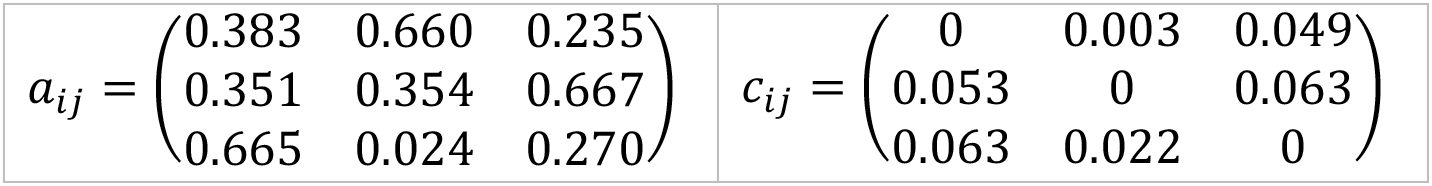

#### (iv) Autocatalytic biosynthesis model (see Fig. 5A and the main text)

##### (a) Quasi-steady-state formula for composite Michaelis-Menten equations

To construct the autocatalytic biosynthesis network model, we used a class of nonlinear flux functions based on quasi-steady-state Michaelis-Menten models (50). All models contained “catalytic enzymes” and one or more “reactants”. In a coarse-grained spirit, we did not include all intermediate metabolites in the model; instead, we used the end product formation flux (the irreversible step in the Michaelis-Menten model) as the flux function. In the following section, we describe the flux functions used in the model. We denote *E*_*total*_ as the total amount of enzyme *E*, including all possible enzyme states (free, bound, etc.).

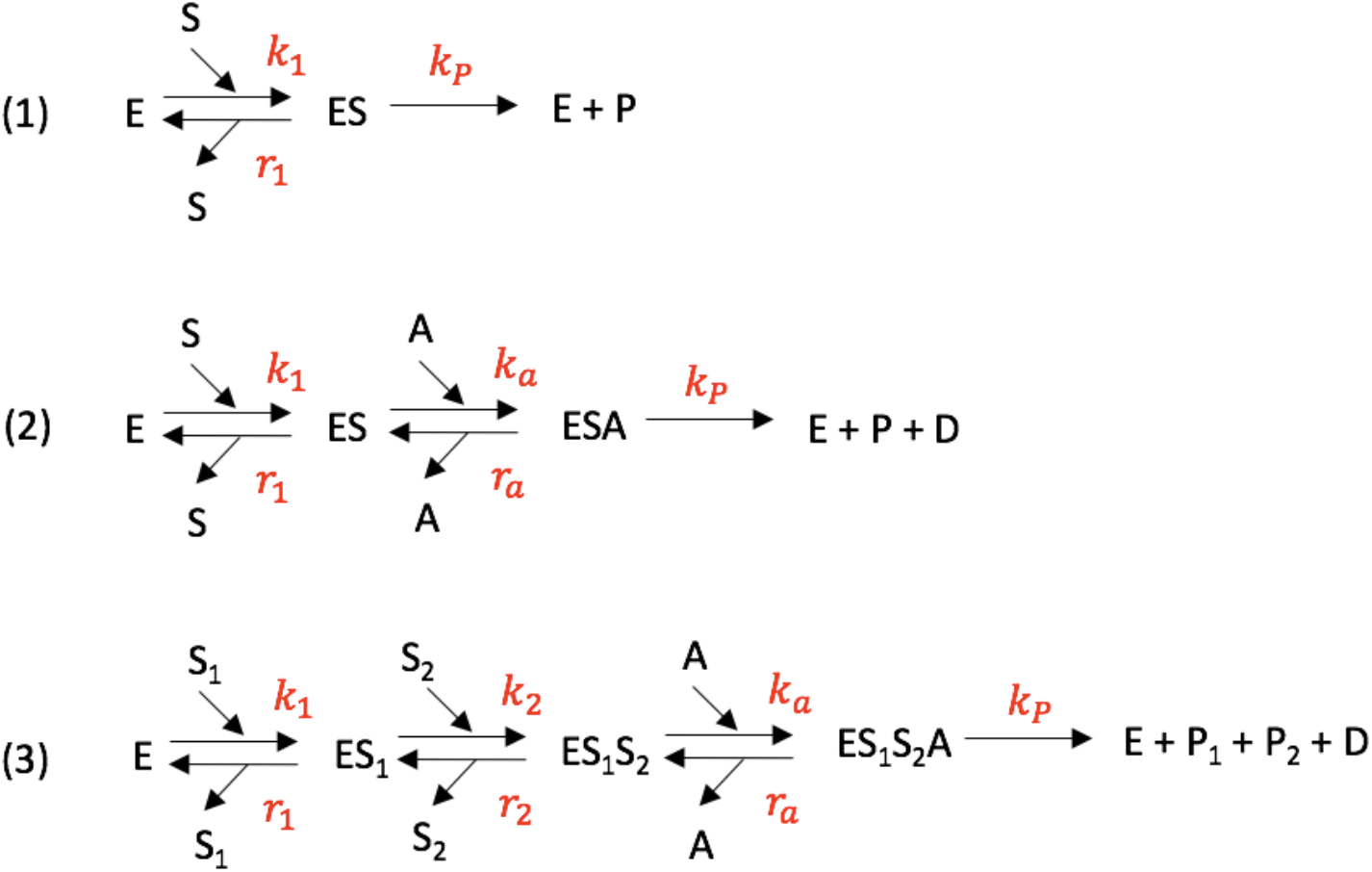

1. One-substrate Michaelis-Menten equation (assuming quasi-steady state on the [ES] complex): The flux function is

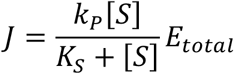

with 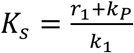.
2. Two-substrate Michaelis-Menten equation (assuming detailed balance between reversible reactions and quasi-steady state on the [ESA] complex): The flux function is

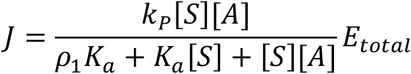

with 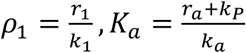.
3. Three-substrate Michaelis-Menten equation (assuming detailed balance between reversible reactions and quasi-steady state on the [*ES*_1_*S*_2_*A*] complex): The flux function is

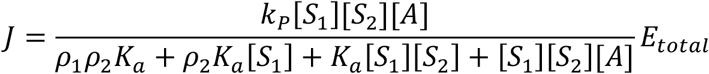

with 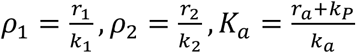.
4. Polymer reaction flux (assuming that the initial complex formation between *E* and *E*_0_to be at equilibrium and that each polymerization step is a two-substrate Michaelis-Menten equation with similar conditions as in (2), for a total of *n* cycles).

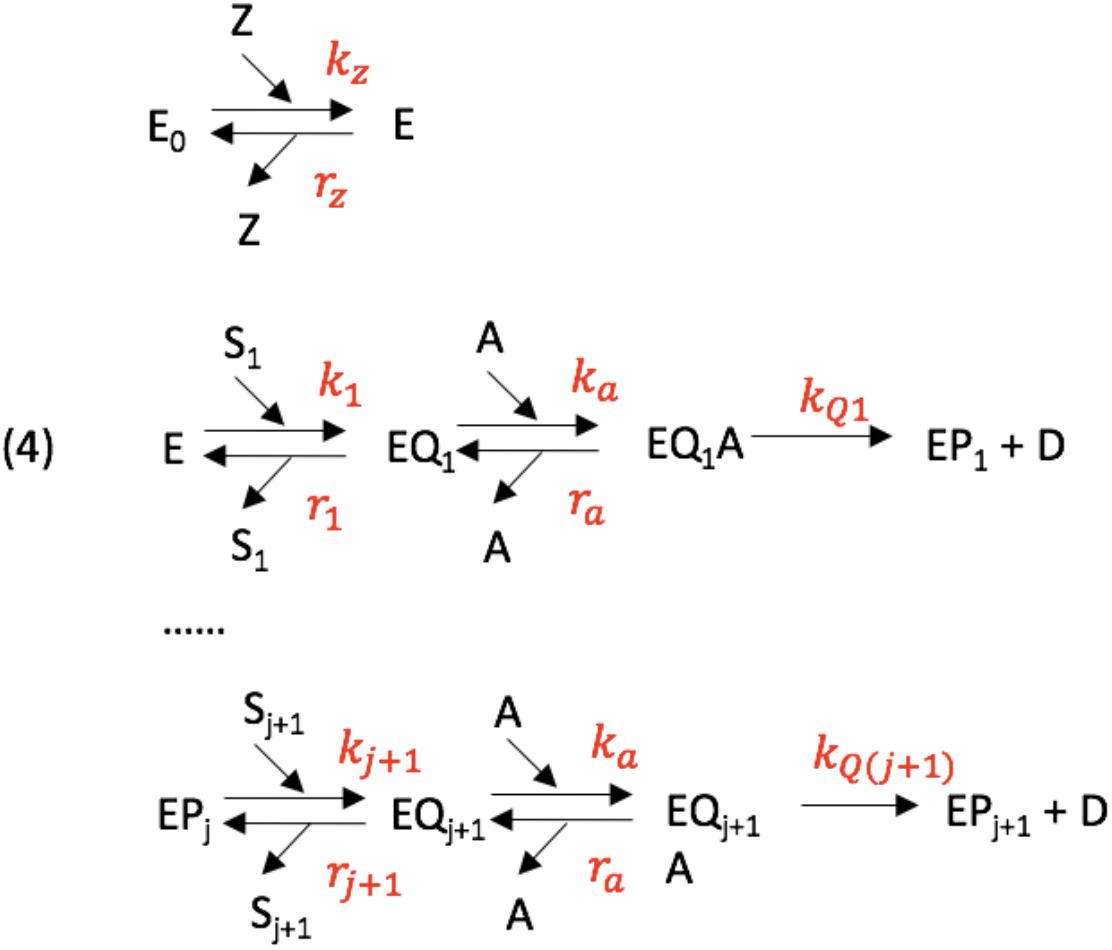

Note that we have the following relation for all *j=*1,…,*n*:

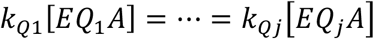

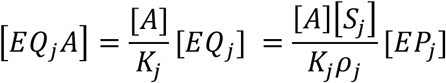

where 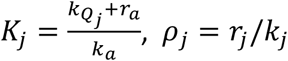.

In particular, 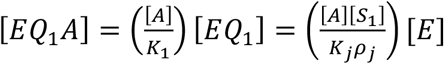. In addition, 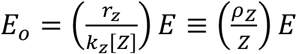.

Therefore, by summing all (bound and unbound) enzymes, we have

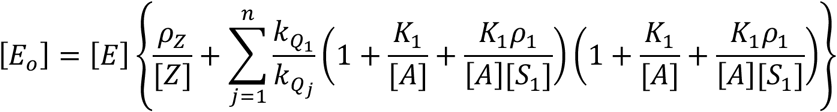

and hence

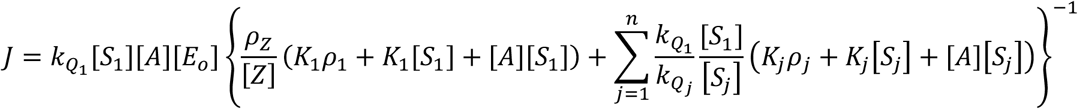

For a polymer requiring {*n*_1_, …, *n*_*m*_} monomers with monomer type 1,…,*m*, the summation can be expressed by

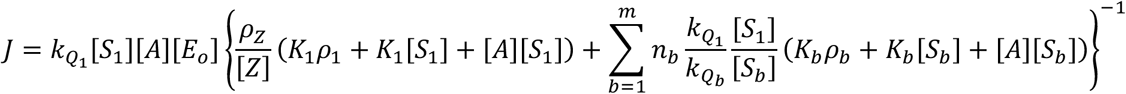

##### (b) Nodes, reactions and flux functions of the autocatalytic biosynthesis model

There are 24 nodes and 40 reactions in our autocatalytic biosynthesis model. All flux functions of the 40 reactions are scalable (see Table S1); their formulas are derived in the previous section. The 24 nodes, including 12 metabolites (red) and 12 polymers (green) are shown in the figure below. The external nutrients *s*_1_, *s*_2_, *s*_3_ belong to the environment and are not part of the system nodes. They are assumed to be maintained at *constant concentration levels* for each simulation condition.

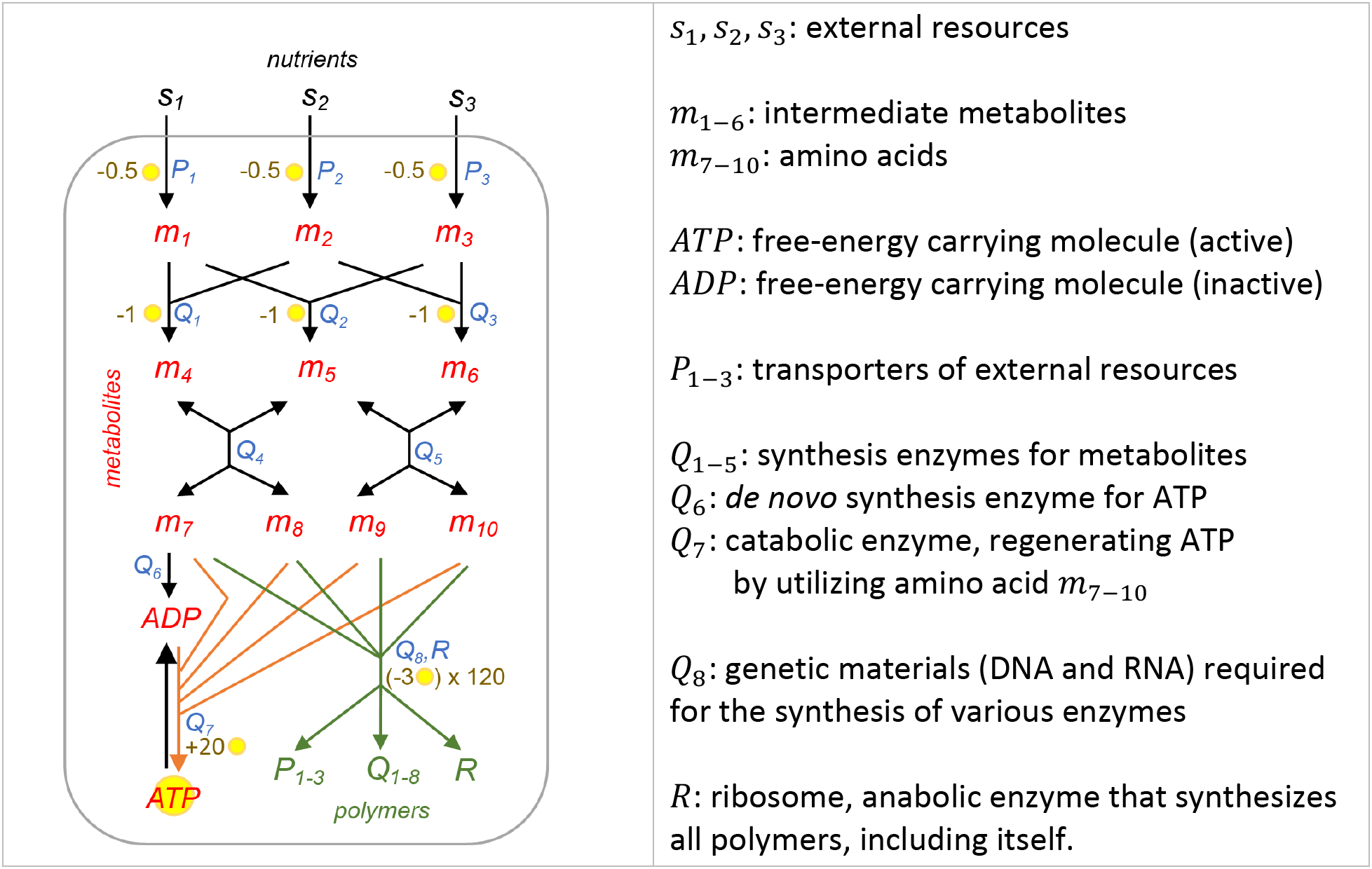

The 40 fluxes are indicated by the arrows in the figure. Note that many fluxes are coupled with ATP → ADP reactions and hence have ATP as an upstream node and ADP as a downstream node. For clarity, all ATP-related arrows are not drawn; instead, the ATP coupling is indicated by the “yellow coin” symbol. The full stoichiometry of the reaction is listed in Table S2.

In addition, many reactions require enzyme catalysis, indicated by blue letters next to the reactions. These enzymes are required for flux, but are not consumed by the reaction. Hence, they do not belong to the upstream or downstream nodes of these reactions. The dependence of these enzymes is shown by the flux function in Table S2.

##### (c) Parameter range of the autocatalytic biosynthesis model

We tested a wide range of parameter values and various initial conditions for the system. Most of the trajectories *Y*(*t*) converged to a fixed point or limit cycle, resulting in convergence of the long-term growth rate. We chose a “standard parameter set” (see Table S3) in which the system trajectory *Y*(*t*) converges to a steady state. Due to the simplicity of our toy model, our parameter set cannot be directly compared to realistic metabolite concentrations. However, we can compare the relative fraction of biomass of each polymer and metabolite between simulations and the cell. Note that all parameters follow the unit system of biomass vector *X*(*t*), which is unitless.

To compare with classical biochemical rate constants, one can use Eq. (5) presented in the main text. We found that under our standard parameter set (Table S3), the 12 metabolites in our toy model account for 4.3% of biomass, while the 12 polymers account for more than 95% of the biomass. Among the polymers, the ribosomes (*Y*_*R*_) and the genetic material 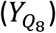 account for ~33% and ~8% of biomass, respectively (see Table S3).

This biomass partition is physiologically realistic for *Escherichia coli*. In this bacterium, macromolecules account for ~95% of the biomass [Ref(51)], with ribosomes occupying 20% to 40% of the biomass of the polymer fraction [Ref(51)]. Polymers that are not ribosomal RNAs (rRNAs) (including mRNA, tRNA and DNA) account for ~7% of the biomass [Ref(52)]. In *E. coli*, the ATP concentration is about 2 mM [Ref(53)], with a molecular weight (MW) = 507 *g*/*mole*.

Suppose the cell is 30% dry weight (i.e., biomass) and the dry weight density is about 1.1 *g*/*cm*^3^[Ref(54)], the biomass fraction of ATP is about 0.33%. In our model we have *Y*^ATP^= 0.45% which is closed to the physiological range. For the energy balance level, the ratio ATP:ADP is about 5-10 folds [Ref(55)]. In our model, *Y*^ATP^: *Y*_ADP_ ~ 4, which is close to the physiological range.

### 5. Analysis of Random networks and autocatalytic circuits

#### (i) Generation of random networks

In the analysis of symbolic reaction networks, an ensemble of random networks is generated according to a given statistical property of the network connections. Each random network is composed of a node set *x* and a reaction set *ϕ*. Each reaction *ϕ*_*a*_ has its associated maintenance set *mt*(*ϕ*_*a*_) ⊆ *x* and downstream set *dw*(*ϕ*_*a*_) ⊆ *x*. We simulated the network as follows:

1. First, we fixed the numbers of nodes and reactions to be *n* and *m* and set two constants *n*_*mt*_, *n*_*dw*_ as the numbers of maintenance nodes and downstream nodes for all reactions. For Fig. S6, we chose *n*_*mt*_ = 3 and *n*_*mt*_ = 2.
2. For each node *x*_*j*_, we assigned a weighting factor 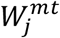 to the node. This weighting factor determines the relative likelihood that node *x*_*j*_ is chosen as the maintenance set of a random reaction. The weighting factors are chosen over the interval (0, *w*_*max*_], with a probability distribution

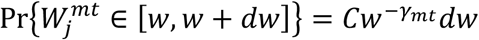

with the normalizing constant 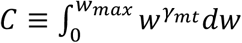. The aforementioned distribution follows a power law with exponent −*γ*_*mt*_ and is truncated in the range (0, *w*_*max*_]. In simulations, *γ*_*mt*_ varied between 0 and 3 and *w*_*max*_ = *γ*_*mt*_|log (10^−4^)| ≈ 9 *γ*_*mt*_. This was sufficient to generate a wide range of values for 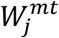 with several orders of magnitude with probability of 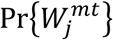.
3. After each node was assigned a weighting factor 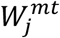, we normalized the weighting factor and acquired a probability function *P*_*mt*_(*j*) by

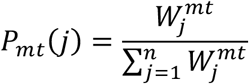

For each reaction and each slot of a maintenance set, a random node was drawn from *x* according to probability *P*_*mt*_(*j*) and assigned to the maintenance node slot. This specified the maintenance set *mt*(*ϕ*_*a*_) for each reaction *ϕ*_*a*_.
4. For each node *x*_*j*_, a similar method is used to generate the node set *dw*(*ϕ*_*a*_) for each reaction *ϕ*_*a*_. We defined 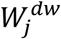 with *γ*_*dw*_ ranging from 0 to 3, similar to 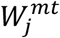. We define 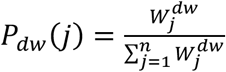. For each reaction and each slot of its downstream set, a random node was drawn from *x* according to probability *P*_*dw*_(*j*) and assigned to the downstream node slot. This specified the downstream node *dw*(*ϕ*_*a*_) for each reaction *ϕ*_*a*_.

#### (ii) Identification of autocatalytic circuits in random networks

To test whether a random network has an autocatalytic circuit, we used the following procedure (for the proof of this algorithm, see the Supplementary Text section in the SI, Algorithm 6.3).

1. Given a collection of reactions *K* ⊆ *ϕ*, we defined

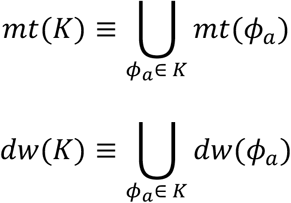
2. We started with a collection of reactions *K*_0_ ≡ *ϕ*. For each *m* ≥ 0, we checked if *mt*(*K*_*m*_) ⊆ *dw*(*K*_*m*_). If true, *K*_*m*_ is an autocatalytic circuit and the network is autocatalytic. If false, we scanned for all reactions and obtained a subset *A*_*m*_: { *ϕ*_*a*_ ∈ *K*_*m*_, *mt*(*ϕ*_*a*_) ∉ *K*_*m*_}. We deleted this reaction subset from *K*_*m*_ and obtained *K*_*m*+1_ ≡ *K*_*m*_\*A*_*m*_.
3. The procedure above was repeated until either *K*_*m*_ became autocatalytic or an empty set. If *K*_*m*_ became an empty set, then the network contained no autocatalytic circuit.

#### (iii) Calculation of the autocatalytic probability for random networks

For Fig. S6, random reaction networks with 100 nodes and various number of reactions were constructed. Each reaction was randomly assigned with three maintenance nodes and two downstream nodes (using sampling with replacement). The probability of assigning a node to a maintenance or downstream set followed a power law exponent *γ*_*mt*_ and *γ*_*dw*_.

For each condition, 500 random networks were generated and tested for their autocatalytic ability using the algorithm shown in the previous section. The autocatalytic probability *P*_*auto*_ was calculated as follows:

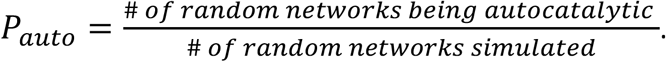

---

**Figure S1:**
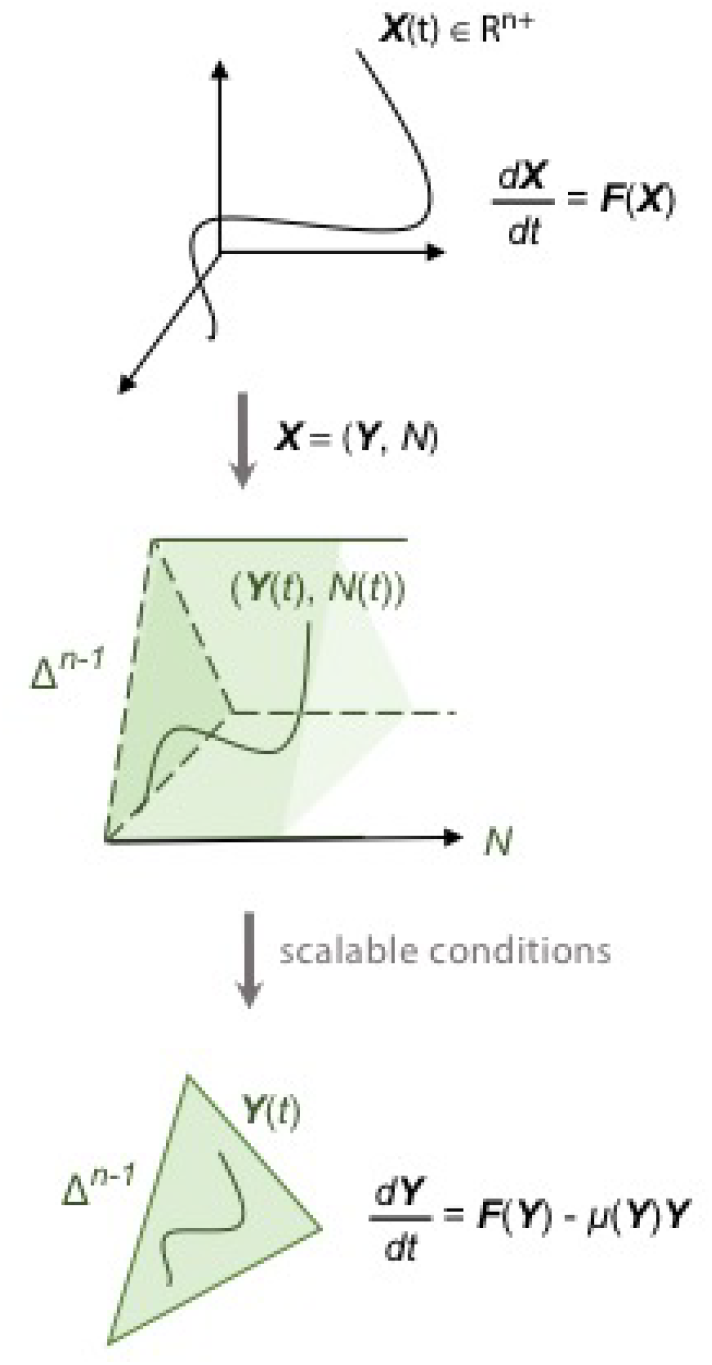
Transformation of a scalable reaction network. Top: System trajectory *X*(*t*) in non-negative quadrant *Q*^*n*^: {*X* ∈ ℝ^*n*^, *X*_*j*_ ≥ 0}. Middle: System trajectory (*Y*(*t*), *N*(*t*)) after transformation from *X* to (*Y*, *N*) coordinate. The state space is Δ^*n*−1^ × [0, ∞), where Δ^*n*−1^:{*X* ∈ ℝ^*n*+^, *X*_1_ + ⋯ + *X*_*n*_ = 1} is the (*n*-1)-dimensional unit simplex. Bottom: For a scalable reaction network, the trajectory *Y*(t) is confined in Δ^*n*−1^ and the system dynamics is fully determined by Eq. (3) in the main text.

**Figure S2:**
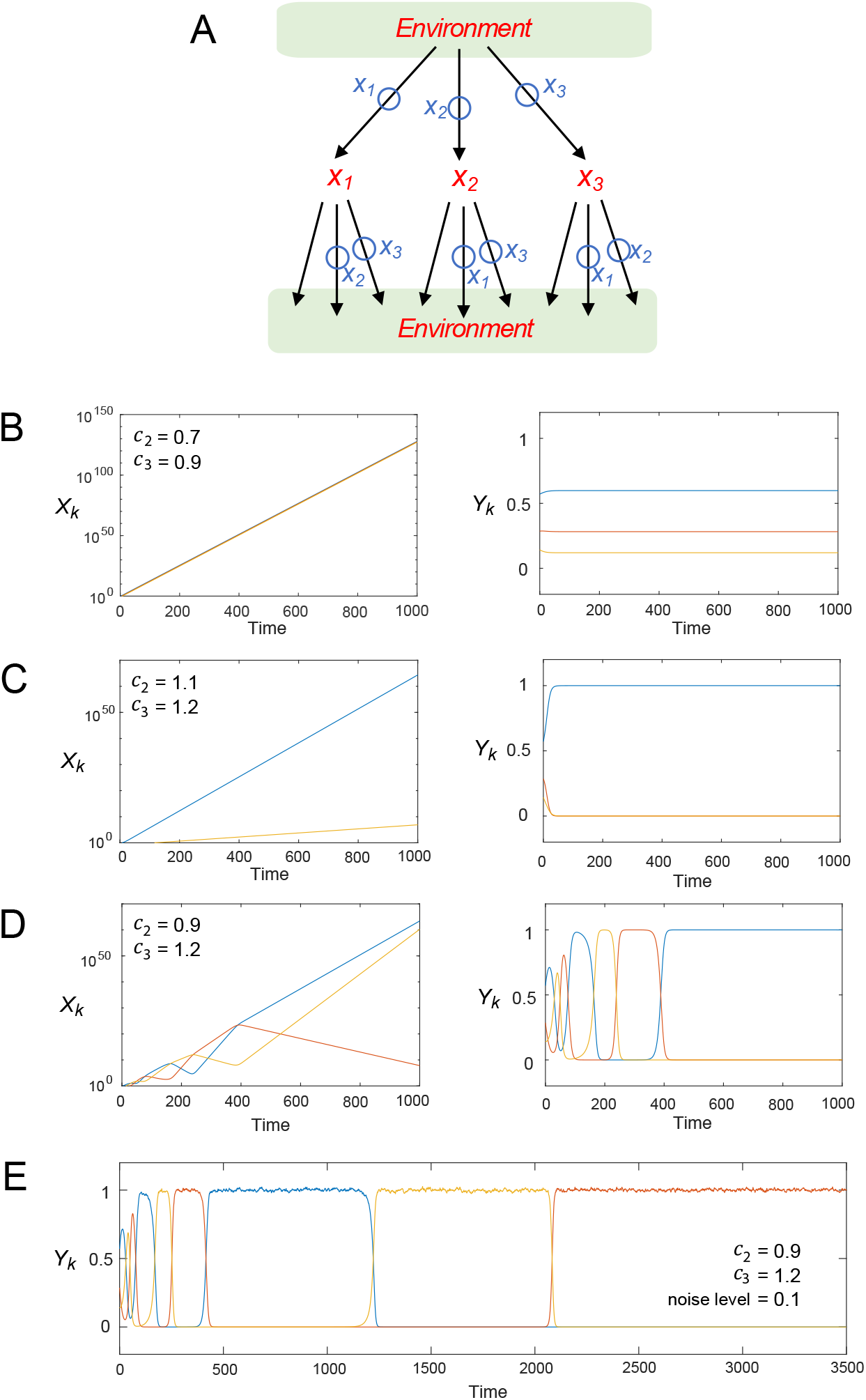
Modified May-Leonard system. **A**, Flux diagram of the system. Blue circles on the flux reactions indicate the nodes that promote the corresponding reaction. For the nonlinear flux function, see example 2.5 in the Supplementary Materials. **B**, System trajectories with *c*_2_ = 0.7, *c*_3_ = 0.9. The blue, orange and yellow lines represent *X*_1_, *X*_2_ and *X*_3_ in the left panel, while representing *Y*_1_, *Y*_2_ and *Y*_3_ in the right panel. In the long-term, *Y*(*t*) converges to a steady-state, with the three species coexisting. **C**, Same as **B** except that *c*_2_ = 1.1, *c*_3_ = 1.2. In the long-term, *Y*(*t*) converges to a steady-state, with a single species dominating. **D**, Same as **B** except that *c*_2_ = 0.9, *c*_3_ = 1.2. In the long-term, *Y*(*t*) converges to a heteroclinic cycle and exhibits oscillations with increasing period length. **E**, Same as **D** except that a scalable noise term *h* ∘ *dB*_*k*_ is added for *k*=1,2,3 with noise level *h* = 0.01. This shows that the heteroclinic cycle (in **C**) is robust under noise perturbation.

**Figure S3:**
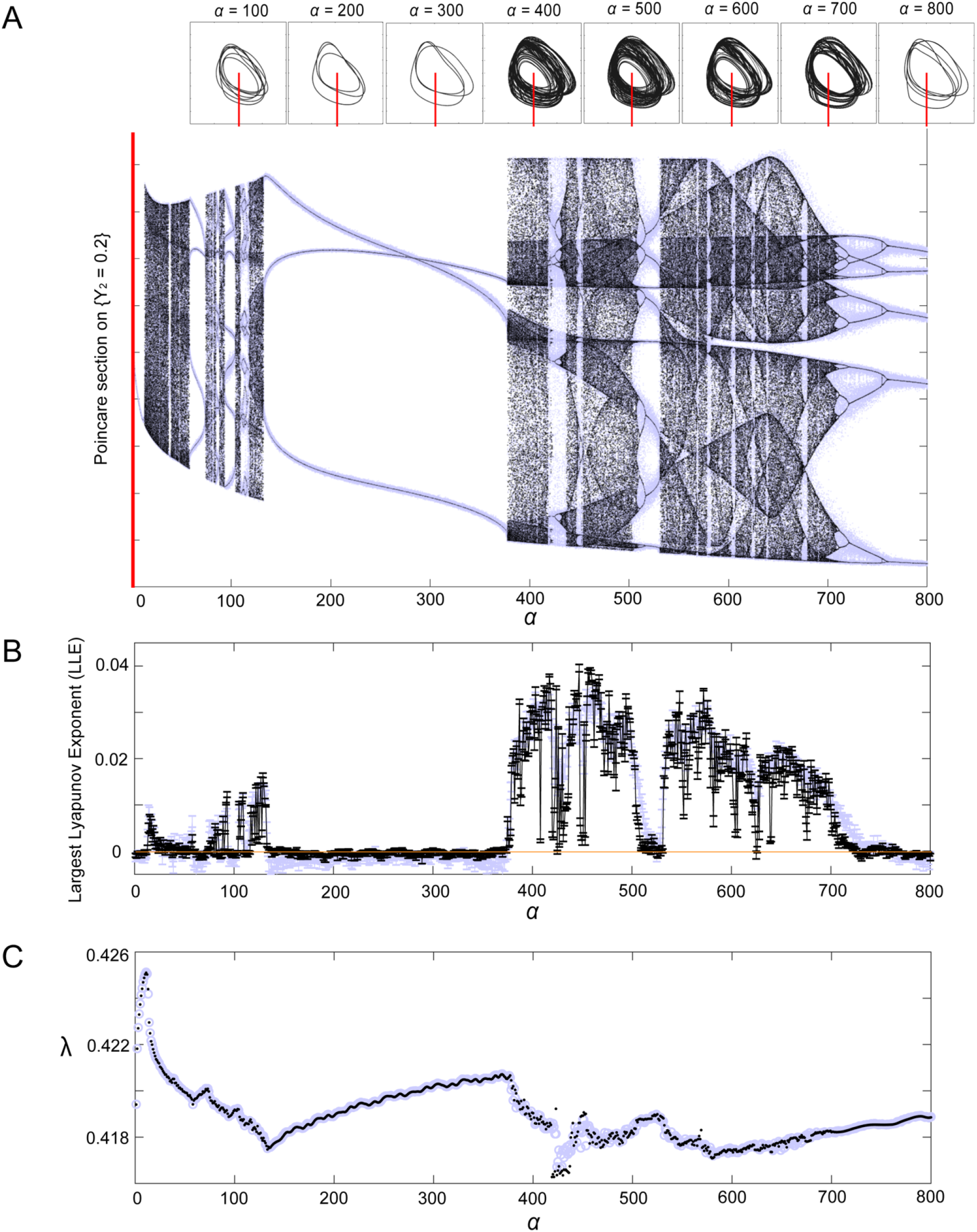
Characterization of the long-term behavior of the double-repressilator model. The analysis is focused on varying parameter *α* = *K*_*a*_/*K*_*b*_, the relative repression strength between the two repressilators. For simulation parameters, see “Methods: Simulation and analysis of scalable networks” in the SI. **A**, Bifurcation diagram of the Poincare section on omega-limit set of *Y*(*t*), with various values of *α*. Upper panel: Trajectory *Y*(*t*) projected on *Y*_2_ − *Y*_3_ subspace, indicating various types of omega-limit sets of *Y*(*t*). Red lines indicate the Poincare section used in the lower panel. Lower panel: Bifurcation diagram on the Poincare section {*Y*_2_ = 0.2} (indicated in the top panel). Black dots show simulation results from a deterministic system, while blue dots are the simulation results from the same system, but with small scalable noise (see SI, Supplementary Text, section 4). **B**, Largest Lyapunov exponent (LLE) of *Y*(*t*) as a function of *α*. Each data point is the average of ten simulation results. The error bars indicate the standard error of the simulated LLE. The orange horizontal line indicates LLE = 0. The regime with positive LLE corresponds to a chaotic regime, which can be compared to **A**. The simulation results from a deterministic system are shown in black, while the ones with small scalable noise are shown in blue. **C**, Long-term growth rate as a function of *α*. The simulation results from a deterministic system are shown in black, while the ones with small scalable noise are in blue.

**Figure S4:**
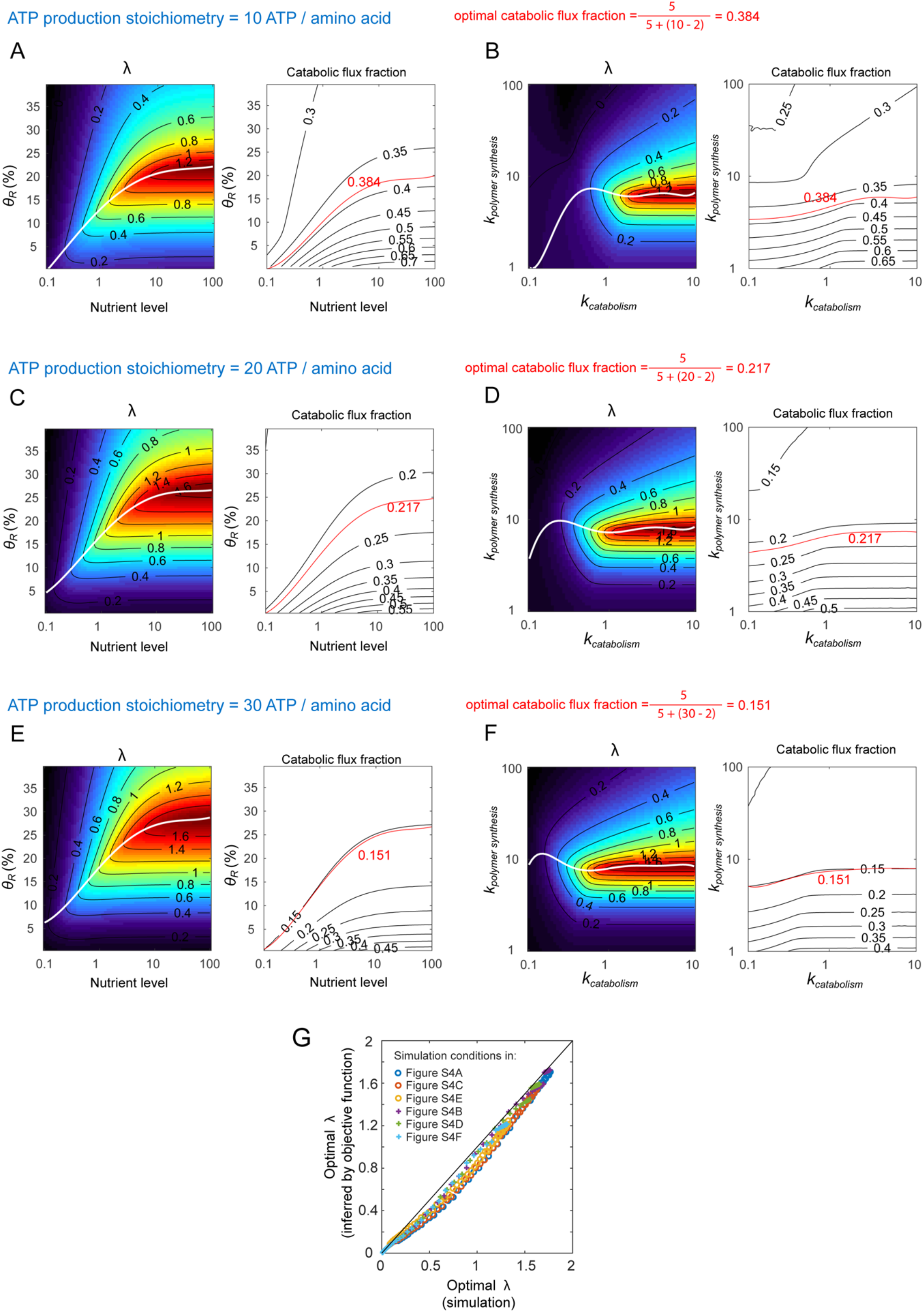
Growth rate optimization under different ATP production stoichiometries. **A**, Growth rate and catabolic flux fraction, with the ATP production stoichiometry set to 10. Left: Growth rate with various *θ*_*R*_ (ribosomal synthesis strength) under various external nutrient levels. The white line indicates the maximal growth rate under each nutrient level. Right: Catabolic flux fraction with various *θ*_*R*_ under various external nutrient levels. The red line indicates the contour of the optimal catabolic flux fraction. **B**, Growth rate and catabolic flux fraction, with the ATP production stoichiometry set to 10. Left: Growth rate with various *k*_*polymer synthesis*_ (polymer synthesis rate constant) under various *k*_*catabolic*_ (catabolic rate constant). The white line indicates the maximal growth rate under each value of *k*_*catabolic*_. Right: catabolic flux fraction with various *k*_*polymer synthesis*_ under various *k*_*catabolic*_. The red line indicates the contour of the optimal catabolic flux fraction. **C**, Same as **A**, except that the ATP production stoichiometry is set to 20. **D**, Same as **B**, except that the ATP production stoichiometry is set to 20. **E**, Same as **A**, except that the ATP production stoichiometry is set to 30. **F**, Same as **B**, except that the ATP production stoichiometry is set to 30. **G**, Comparison between the optimal growth rate obtained by simulation and the optimal growth rate inferred from the objective function. This plot contains six different types of optimization conditions in which ATP production stoichiometry is 30, 20 and 10 per amino acid (*m*_7-10_), respectively.

**Figure S5:**
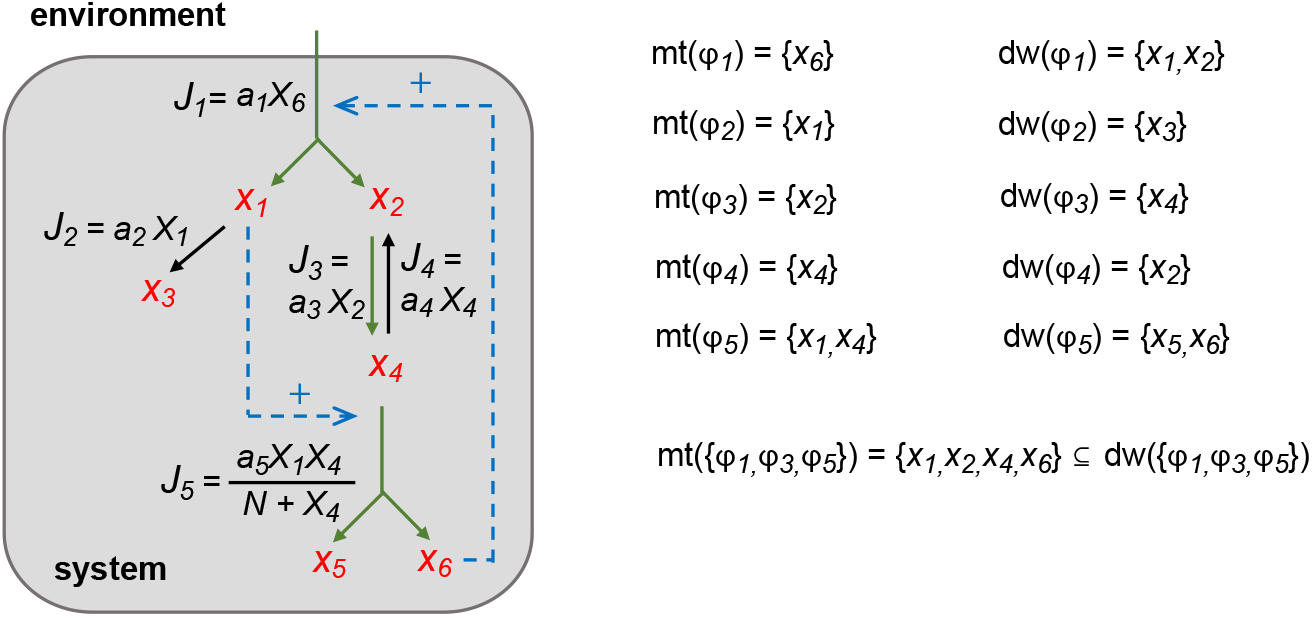
Example of an autocatalytic circuit in a scalable reaction network. In this system, there are five reactions with associated flux function forms specified in the figure. The maintenance and downstream sets of reactions are listed on the right. A subset of reactions *K* ≡ {*ϕ*_1_, *ϕ*_3_, *ϕ*_5_} satisfies *mt*(*K*) ⊆ *dw*(*K*), indicating that this reaction network is autocatalytic.

**Figure S6:**
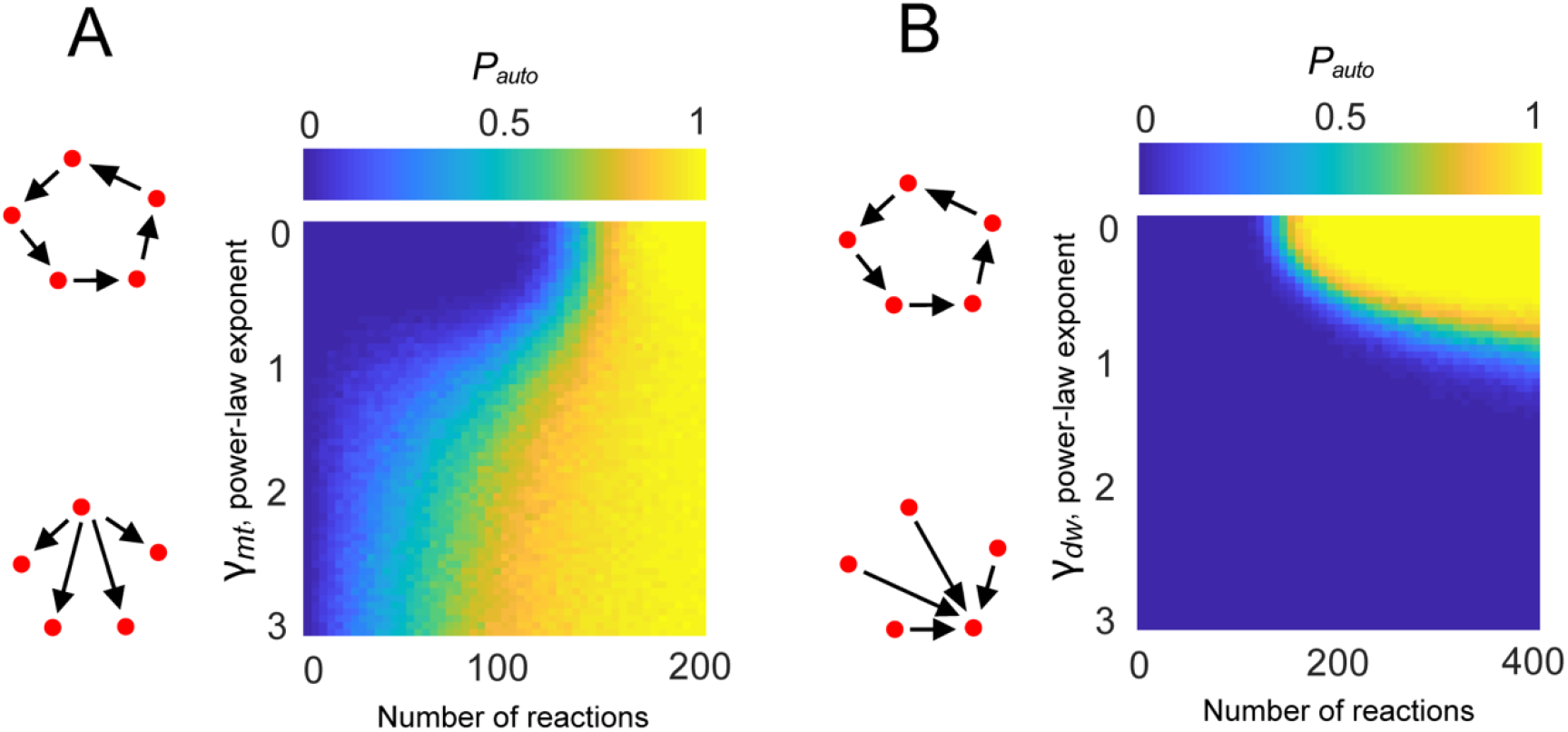
Phase diagram of *P*_*auto*_, the probability of a random network to be autocatalytic. Ensembles of random networks were simulated under different power-law connection probabilities and various numbers of reactions. All random reaction networks have 100 nodes and each reaction has three maintenance nodes and two downstream nodes. **A**, Network with power-law exponent *γ*_*dw*_ = 0 and *γ*_*mt*_ varying from 0 (uniform connection) to 3 (hub-like maintenance nodes). **B**, Network with power-law exponent *γ*_*dw*_ = 0 and *γ*_*dw*_ varying from 0 (uniform connection) to 3 (hub-like downstream nodes).

**Table S1:**
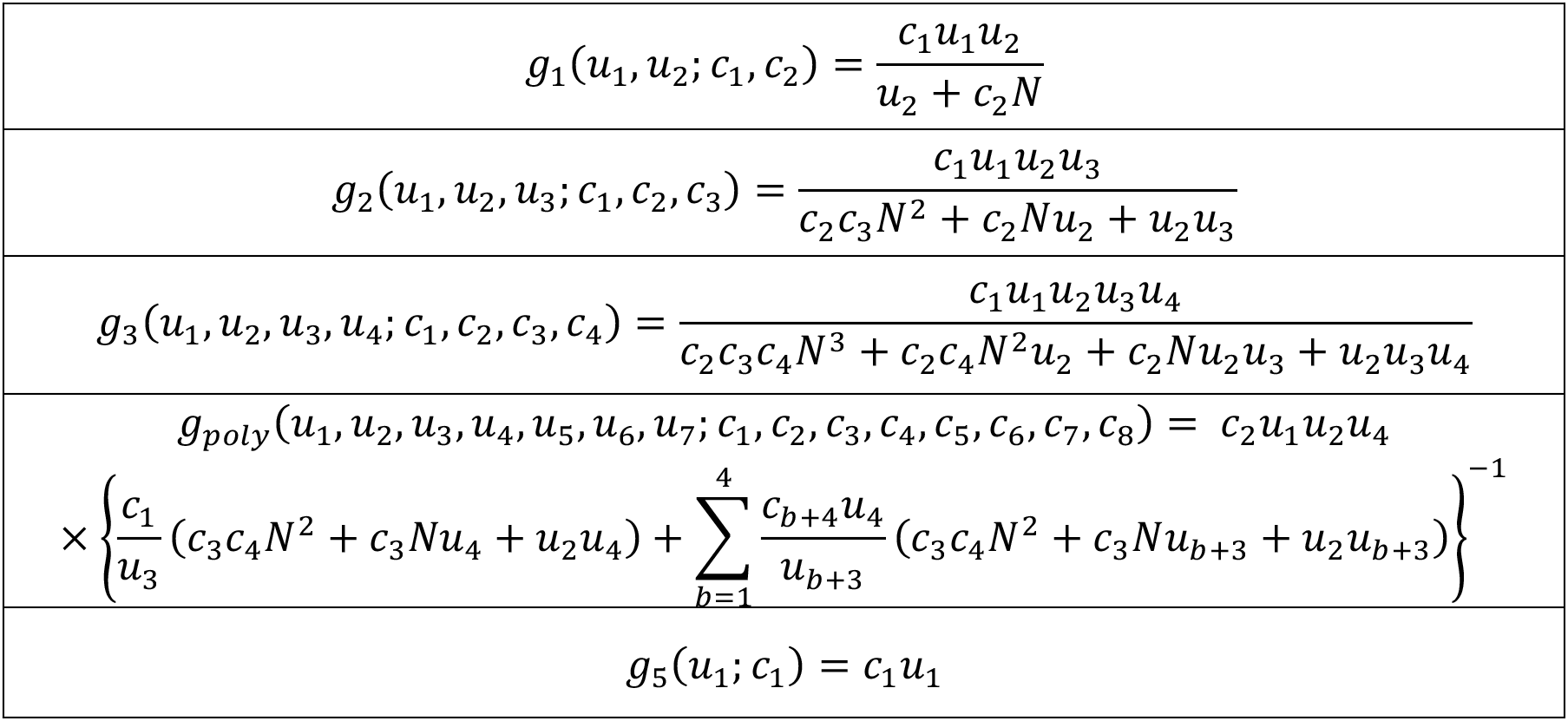
Common flux functions. This table lists the scalable flux functions used in the biosynthesis model. Here, *u*_*j*_ are variables of the scalable flux function and we denote *N* = *u*_1_ + ⋯ + *u*_*n*_. The parameters in the function are denoted as *c*_*j*_. The formulas are derived from Michaelis-Menten kinetics on quasi-steady-state reactions (see Method section: Mathematical models of SRN iv (a)), assuming that the cell volume is proportional to the total biomass *N* during the system growth.

**Table S2:**
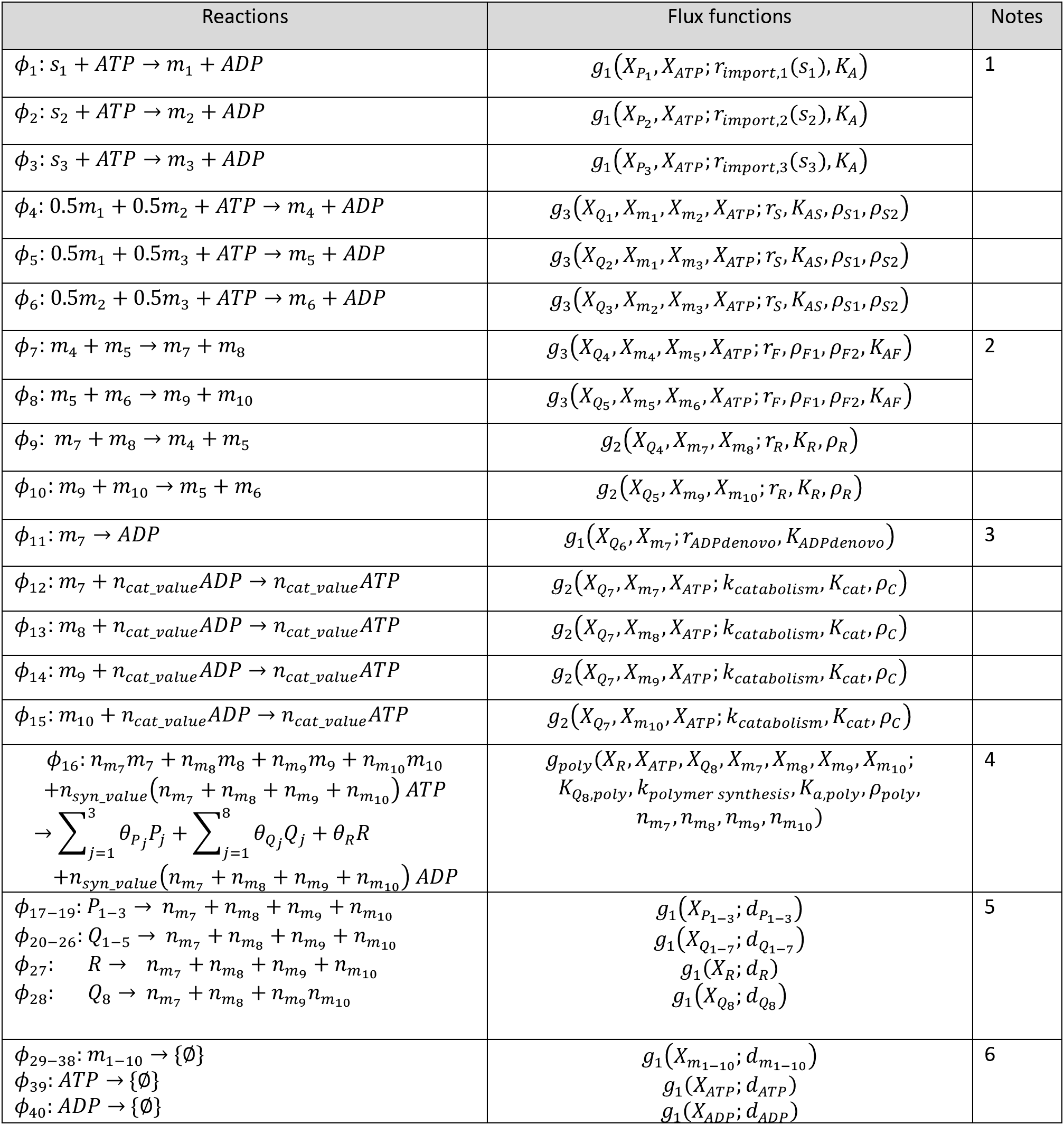
Reactions and flux functions in the autocatalytic biosynthesis model. This table lists 24 reactions in the autocatalytic biosynthesis network, with the corresponding flux function. The mathematical formulas of these flux functions (*g*_1_, *g*_2_, *g*_3_, *g*_*poly*_ and *g*_5_) are described in Table S1. Here, the subscript of the variable *X*_(·)_ denotes the name of the corresponding node.

| Reactions | Flux functions | Notes |
| --- | --- | --- |
| $\phi_1: s_1 + ATP \rightarrow m_1 + ADP$ | $g_1(X_{P_1}, X_{ATP}; r_{import,1}(s_1), K_A)$ | 1 |
| $\phi_2: s_2 + ATP \rightarrow m_2 + ADP$ | $g_1(X_{P_2}, X_{ATP}; r_{import,2}(s_2), K_A)$ | |
| $\phi_3: s_3 + ATP \rightarrow m_3 + ADP$ | $g_1(X_{P_3}, X_{ATP}; r_{import,3}(s_3), K_A)$ | |
| $\phi_4: 0.5m_1 + 0.5m_2 + ATP \rightarrow m_4 + ADP$ | $g_3(X_{Q_1}, X_{m_1}, X_{m_2}, X_{ATP}; r_S, K_{AS}, \rho_{S1}, \rho_{S2})$ | |
| $\phi_5: 0.5m_1 + 0.5m_3 + ATP \rightarrow m_5 + ADP$ | $g_3(X_{Q_2}, X_{m_1}, X_{m_3}, X_{ATP}; r_S, K_{AS}, \rho_{S1}, \rho_{S2})$ | |
| $\phi_6: 0.5m_2 + 0.5m_3 + ATP \rightarrow m_6 + ADP$ | $g_3(X_{Q_3}, X_{m_2}, X_{m_3}, X_{ATP}; r_S, K_{AS}, \rho_{S1}, \rho_{S2})$ | |
| $\phi_7: m_4 + m_5 \rightarrow m_7 + m_8$ | $g_3(X_{Q_4}, X_{m_4}, X_{m_5}, X_{ATP}; r_F, \rho_{F1}, \rho_{F2}, K_{AF})$ | 2 |
| $\phi_8: m_5 + m_6 \rightarrow m_9 + m_{10}$ | $g_3(X_{Q_5}, X_{m_5}, X_{m_6}, X_{ATP}; r_F, \rho_{F1}, \rho_{F2}, K_{AF})$ | |
| $\phi_9: m_7 + m_8 \rightarrow m_4 + m_5$ | $g_2(X_{Q_4}, X_{m_7}, X_{m_8}; r_R, K_R, \rho_R)$ | |
| $\phi_{10}: m_9 + m_{10} \rightarrow m_5 + m_6$ | $g_2(X_{Q_5}, X_{m_9}, X_{m_{10}}; r_R, K_R, \rho_R)$ | |
| $\phi_{11}: m_7 \rightarrow ADP$ | $g_1(X_{Q_6}, X_{m_7}; r_{ADPdenovo}, K_{ADPdenovo})$ | 3 |
| $\phi_{12}: m_7 + n_{cat\_value}ADP \rightarrow n_{cat\_value}ATP$ | $g_2(X_{Q_7}, X_{m_7}, X_{ATP}; k_{catabolism}, K_{cat}, \rho_C)$ | |
| $\phi_{13}: m_8 + n_{cat\_value}ADP \rightarrow n_{cat\_value}ATP$ | $g_2(X_{Q_7}, X_{m_8}, X_{ATP}; k_{catabolism}, K_{cat}, \rho_C)$ | |
| $\phi_{14}: m_9 + n_{cat\_value}ADP \rightarrow n_{cat\_value}ATP$ | $g_2(X_{Q_7}, X_{m_9}, X_{ATP}; k_{catabolism}, K_{cat}, \rho_C)$ | |
| $\phi_{15}: m_{10} + n_{cat\_value}ADP \rightarrow n_{cat\_value}ATP$ | $g_2(X_{Q_7}, X_{m_{10}}, X_{ATP}; k_{catabolism}, K_{cat}, \rho_C)$ | |
| $\phi_{16}: n_{m_7}m_7 + n_{m_8}m_8 + n_{m_9}m_9 + n_{m_{10}}m_{10}$<br>$+ n_{syn\_value}(n_{m_7} + n_{m_8} + n_{m_9} + n_{m_{10}})ATP$<br>$\rightarrow \sum_{j=1}^3 \theta_{P_j}P_j + \sum_{j=1}^8 \theta_{Q_j}Q_j + \theta_R R$<br>$+ n_{syn\_value}(n_{m_7} + n_{m_8} + n_{m_9} + n_{m_{10}})ADP$ | $g_{poly}(X_R, X_{ATP}, X_{Q_8}, X_{m_7}, X_{m_8}, X_{m_9}, X_{m_{10}};$<br>$K_{Q_8,poly}, k_{polymer\ synthesis}, K_{a,poly}, \rho_{poly},$<br>$n_{m_7}, n_{m_8}, n_{m_9}, n_{m_{10}})$ | 4 |
| $\phi_{17-19}: P_{1-3} \rightarrow n_{m_7} + n_{m_8} + n_{m_9} + n_{m_{10}}$<br>$\phi_{20-26}: Q_{1-5} \rightarrow n_{m_7} + n_{m_8} + n_{m_9} + n_{m_{10}}$<br>$\phi_{27}: R \rightarrow n_{m_7} + n_{m_8} + n_{m_9} + n_{m_{10}}$<br>$\phi_{28}: Q_8 \rightarrow n_{m_7} + n_{m_8} + n_{m_9} + n_{m_{10}}$ | $g_1(X_{P_{1-3}}; d_{P_{1-3}})$<br>$g_1(X_{Q_{1-7}}; d_{Q_{1-7}})$<br>$g_1(X_R; d_R)$<br>$g_1(X_{Q_8}; d_{Q_8})$ | 5 |
| $\phi_{29-38}: m_{1-10} \rightarrow \{\emptyset\}$<br>$\phi_{39}: ATP \rightarrow \{\emptyset\}$<br>$\phi_{40}: ADP \rightarrow \{\emptyset\}$ | $g_1(X_{m_{1-10}}; d_{m_{1-10}})$<br>$g_1(X_{ATP}; d_{ATP})$<br>$g_1(X_{ADP}; d_{ADP})$ | 6 |

Note 1: The external nutrient level *s*_*j*_ affects the parameter *r*_*import,j*_, with the Michaelis-Menten form 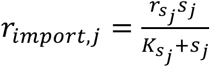 for *j* = 1,2,3.

Note 2: The reactions *ϕ*_7_ and *ϕ*_8_ do not consume ATP, but require ATP as an intermediate in the flux function (i.e., ATP acts as a cofactor for the enzymes *Q*_4_ and *Q*_5_).

Note 3: Reaction *ϕ*_11_ is the *de novo* synthesis of ADP. This step does not involve ATP.

Note 4: Reaction *ϕ*_16_ describes the process of overall polymer synthesis. It uses amino acids *m*_7-10_ and ATP energy, and generates 12 types of polymers with stoichiometry parameters 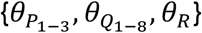. By incorporating the stoichiometry coefficients into the common kinetic parameter *k*_*polyymer synthesis*_, the coefficient *θ*_*α*_ can be normalized such that ∑_*α*_ *θ*_*α*_ = 1. Hence, *θ*_*α*_ represents the fraction of polymer synthesis flux. Biologically, *θ*_*α*_ can be regarded as the (relative) polymer synthesis strength, which is the fraction of the total capacity of polymer synthesis occupied by the polymer type *α*. In the main text, the ribosomal synthesis strength *θ*_*R*_ is set to different values to vary the ribosomal fraction and change the polymer partition.

Note 5: Reactions *ϕ*_17-28_ represent first-order polymer degradation fluxes. The polymer is degraded to regenerate amino acids *m*_7-10_. These reactions are not coupled with ATP consumption.

Note 6: Reaction *ϕ*_29-40_ represent first-order monomer degradation fluxes. These reactions do not have downstream nodes and can be regarded as degradation of metabolites.

**Table S3:** Parameters used in the autocatalytic biosynthesis model. This table shows the parameters used in the simulations. Parameters that were varied in this study are shown in blue.

| Parameter | Value used in the standard parameter set | Reaction involved | Parameter regime tested |
| --- | --- | --- | --- |
| $r_{import,1-3}$ | $r_{import,j} = \frac{r_{sj}s_j}{K_{sj}+s_j} = 54.5 \text{ (1/t)}$<br>$s_1 = s_2 = s_3 = 10 \text{ (mM)}$<br>$r_{sj} = 60 \text{ (1/t)}$<br>$K_{sj} = 1 \text{ (mM)}$ | $\phi_{1-3}$ | $s_j = 0.01 \text{ mM} - 100 \text{ mM}$ |
| $K_A$ | 0.005 | $\phi_{1-3}$ | |
| $r_S$ | 120 (1/t) | $\phi_{4-6}$ | |
| $K_{AS}$ | 0.005 | $\phi_{4-6}$ | |
| $\rho_{S1}$ | 0.01 | $\phi_{4-6}$ | |
| $\rho_{S2}$ | 0.01 | $\phi_{4-6}$ | |
| $r_F$ | 250 (1/t) | $\phi_{7-8}$ | |
| $\rho_{F1}$ | 0.005 | $\phi_{7-8}$ | |
| $\rho_{F2}$ | 0.005 | $\phi_{7-8}$ | |
| $K_{AF}$ | 0.005 | $\phi_{7-8}$ | |
| $r_R$ | 1 (1/t) | $\phi_{9-10}$ | |
| $K_R$ | 0.04 | $\phi_{9-10}$ | |
| $\rho_R$ | 1 | $\phi_{9-10}$ | |
| $r_{ADPdenovo}$ | 1 | $\phi_{11}$ | |
| $K_{ADPdenovo}$ | 0.05 | $\phi_{11}$ | |
| $k_{catabolism}$ | 6 (1/t) | $\phi_{12-15}$ | 0.1-10 (1/t) |
| $K_{cat}$ | 0.001 (Fig. 3B, Table S4)<br>0.005 (Fig. 3F, Fig. S4) | $\phi_{12-15}$ | |
| $\rho_C$ | 0.05 | $\phi_{12-15}$ | |
| $n_{cat \text{ value}}$ | 20 | $\phi_{12-15}$ | 10 - 30 |
| $n_{syn \text{ value}}$ | 3 | $\phi_{16}$ | |
| $K_{Q8,poly}$ | 0.002 | $\phi_{16}$ | |
| $k_{polymer \text{ synthesis}}$ | 10 | $\phi_{16}$ | 1 - 100 |
| $K_{a,poly}$ | 0.0025 | $\phi_{16}$ | |
| $\rho_{poly}$ | 0.005 | $\phi_{16}$ | |
| $n_{m7} - n_{m10}$ | 30 | $\phi_{16-28}$ | |
| $\theta_\alpha$ | $\theta_{P_{1-3}} = 2.0\%$, $\theta_{Q_{1-5}} = 5.2\%$<br>$\theta_{Q_6} = 6.9\%$, $\theta_{Q_7} = 17.3\%$<br>$\theta_{Q_8} = 8.7\%$, $\theta_R = 35\%$ | $\phi_{16}$ | $\theta_R = 0\% - 40\%$ |
| $d_\alpha$ (polymer) | 0.005 | $\phi_{17-28}$ | |
| $d_\beta$ (monomer) | 0.005 | $\phi_{29-40}$ | |

**Table S4:** Steady-state *Y*\* of the standard parameter set. This table lists the steady-state biomass fraction of the autocatalytic biosynthesis model, simulated by the standard parameter set (see Table S3). At this steady state, *λ* = 1.093 (1/*t*).

| Node | % of biomass | Node | % of biomass |
| --- | --- | --- | --- |
| $Y_{m_1}$ | 0.3913 | $Y_{P_1}$ | 1.9136 |
| $Y_{m_2}$ | 0.3913 | $Y_{P_2}$ | 1.9136 |
| $Y_{m_3}$ | 0.3913 | $Y_{P_3}$ | 1.9136 |
| $Y_{m_4}$ | 0.1393 | $Y_{Q_1}$ | 4.9809 |
| $Y_{m_5}$ | 0.1393 | $Y_{Q_2}$ | 4.9809 |
| $Y_{m_6}$ | 0.1393 | $Y_{Q_3}$ | 4.9809 |
| $Y_{m_7}$ | 0.5128 | $Y_{Q_4}$ | 4.9809 |
| $Y_{m_8}$ | 0.5517 | $Y_{Q_5}$ | 4.9809 |
| $Y_{m_9}$ | 0.5517 | $Y_{Q_6}$ | 6.6411 |
| $Y_{m_{10}}$ | 0.5517 | $Y_{Q_7}$ | 16.6028 |
| $Y_{ATP}$ | 0.4510 | $Y_{Q_8}$ | 8.3015 |
| $Y_{ADP}$ | 0.1113 | $Y_R$ | 33.4871 |

